# Controlling membrane tension by dynamic DNA nanorings

**DOI:** 10.64898/2026.09.22.753646

**Authors:** Longfei Liu, Eason Cao, Chunxiang Wu, Abhijith Radhakrishnan, Manindra Bera, Sudhanshu Gautam, Aniruddha Panda, Kallol Gupta, Yong Xiong, Frédéric Pincet, Chenxiang Lin

## Abstract

Membrane tension, defined as the energy cost to expand the membrane surface area, is a fundamental mechanical property of the lipid bilayer and an important modulator of various cellular processes. Liposomes provide model systems to study protein-mediated membrane dynamics, where membrane tension can be controlled by osmotic pressure and micropipette aspiration. However, it is challenging to individually control membrane tension of nanometer-sized liposomes, such as small unilamellar vesicles (SUVs), that are widely used for modeling subcellular membrane structures and drug delivery. To bridge this technological gap, we present reconfigurable DNA origami rings for dynamically controlling membrane tension of sub-100 nm liposomes. The DNA nanorings template the formation of uniformly sized SUVs and undergo trigger-responsive dilation and contraction, thus applying controlled and reversible mechanical stress to the SUV membranes. Electron microscopy analyses show deformed liposomes with ∼2.5% expanded membrane area following ring dilation. The corresponding increase in membrane tension opens a mechanosensitive ion channel MscL with gating tension of ∼10 mN/m. Selectively manipulating a mixture of SUV populations enables timed release of their distinct molecular cargos, either concurrently or one at a time. Furthermore, vesicles with expanded membranes are more conducive to SNARE-mediated fusion. The nanomechanical device thus provides a programmable platform for engineering the mechanics of synthetic nano-vesicles and studying membrane tension modulated processes at the molecular level.

## Introduction

Membrane tension is delicately maintained when cells are at rest and changes drastically when cells engage in energy-demanding activities, including division, migration, and signaling^1,2^. Proteins modulate membrane tension by changing membrane shapes (e.g., budding, fusion) and adjusting lipid composition (e.g., lipid transport and synthesis). In turn, membrane tension regulates protein activities (e.g., actin polymerization and ion channel conductance) as well as the membrane’s susceptibility to remodeling^3^. Therefore, understanding such bidirectional regulation mechanisms, which give rise to diverse and dynamic membrane tension via numerous pathways, is essential to unraveling membrane-tension regulated cell behaviors.

Despite its central role in many cell processes, membrane tension is difficult to precisely control in cells. While it is possible to perturb tension of the plasma membrane by mechanical, biochemical and genetical means, the corresponding molecular events can be obfuscated by the crowded and ever-changing cellular environment. Intracellular membranes pose additional challenges for tension control due to inaccessibility.

Model membranes built with well-defined molecular composition and geometry thus provide a valuable platform to study how protein structure and function and membrane dynamics are influenced by membrane tension. Small unilamellar vesicle (SUV) is a prominent example of model membrane systems. With established methods to produce protein-embedded SUVs in large quantities, these nanometer-sized liposomes (∼20–100 nm in diameter) have been widely used for biochemical and biophysical analyses of lipid bilayers^4,5^ and membrane proteins^6-12^. However, unlike their micrometer-sized counterparts (giant unilamellar vesicles or GUVs), whose membrane tension can be precisely controlled and measured by micropipette aspiration and membrane tether pulling (often by optical tweezers, atomic force microscopy and micropipette techniques)^13-17^, SUVs are difficult to mechanically manipulate using conventional top-down approaches due to their small size. To date, researchers almost exclusively rely on osmotic pressure to adjust membrane tension of SUVs^18^, which lacks the ability to differentially control membrane tension within a mixture (i.e., all liposomes are subject to the same tonicity) and may lead to unintended protein behaviors due to the membrane’s altered physicochemical surroundings (e.g., ionic strength, viscosity).

In recent years, DNA origami^19^ structures have emerged as a versatile toolset for membrane engineering at the nanoscale^20,21^. Equipped with hydrophobic moieties, self-assembled DNA nanostructures can template the formation of size and shape defined liposomes^22,23^ and drive programmable membrane remodeling^24-26^. Reconfigurable DNA nanostructures designed to transform shapes upon receiving specific biochemical inputs have been shown to exert mechanical force on membranes, causing liposome fusion^23^, tubulation^27^, and budding^28,29^. In several instances, the regulatory role of membrane tension in DNA origami mediated membrane deformation was implied^27,30^. The existing DNA-based membrane engineering methods laid the technological foundation but also underscore the need for developing additional molecular tools for controlling membrane tension.

In this work, we built DNA origami rings with dynamically variable diameters to apply tension to SUV membranes. Designed as tensegrity structures that can be toggled between constricted and dilated conformations via DNA strand displacement, these devices reversibly stretched the SUVs they encompassed. This generated programmable membrane tension up to ∼10 mN/m, approaching the lysing tension of SUVs and sufficient to activate mechanically gated ion channel MscL. Stressing specific subpopulation of mixed vesicle species led to selective content release via embedded protein channels. Moreover, we showed that higher membrane tension predisposed SUVs to SNARE-mediated fusion, in line with theoretical predictions. The dilatable DNA rings provide a tool to control membrane tension of nanoscale vesicles in a way that was previously possible only at the micrometer scale, opening opportunities to study the impact of varying membrane tension on membrane protein structure and assembly, protein-membrane interactions, and membrane dynamics.

## Results

### Design and operation of dilatable DNA nanorings

We designed two ring-shaped DNA origami devices differing in size (small and large), each with multiple conformations (free, closed, and open) to control membrane tension of a liposome held within the device (**Figures S1**). The small, free-state DNA nanoring (SRf) consists of two concentric 6-helix-bundle (6hb) rings interconnected by three groups of four 32-nucleotide (nt) single-stranded DNA (ssDNA) tendons (**Figure 1a**, left). The stationary outer ring has a diameter of 68 nm, while the reconfigurable inner ring contains three 6hb arcs of equal lengths bridged by three groups of four 52-nt ssDNA tendons. The counteracting entropic forces exerted by the tendons created a tensegrity structure^31^ where a semi-open inner ring (theoretical inner diameter: 39 nm, see “**Prediction of ‘free’-state DNA nanoring dimensions**” in the Supplementary Materials for details) is ‘suspended’ within the outer ring. The inner arcs can be forced together by converting the tendons connecting the inner and outer rings into anti-parallel double-stranded DNA (dsDNA) zippers, forming a closed ring with inner diameter of 34 nm (SRc, **Figure 1a**, middle). Conversely, the inner arcs can be pushed apart by rigidifying their bridging tendons into 17-nm-long dsDNA struts, thereby transitioning the device to an open state (SRo) with the inner ring diameter of 50 nm (**Figure 1a**, right). To facilitate reversible opening and closure, tendon-hybridizing DNA oligonucleotides (operating strands, dark red and green in **Figure 1a**) were designed with 8-nt overhangs, so that dsDNA struts can return to the single-stranded form via toehold-mediated strand displacement (TMSD). Following the same principle, we designed a larger device (outer ring diameter: 88 nm, **Figure 1c**) featuring a four-arc inner ring with variable diameters of 57 nm (closed state, LRc), 63 nm (free state, LRf), and 73 nm (open state, LRo). Both large and small rings self-assembled with decent yield in the free state (**Figure S2**). Negative-stain transmission electron microscopy (TEM) verified the homogenous geometry of purified free-state rings as well as their dilation and constriction products (**Figures 1b, 1c, S3–S8**). For both devices, the measured inner ring diameters closely matched the designs during two rounds of DNA-triggered reconfiguration (i.e., free→closed→open→closed→open, **Figures 1d, 1e**).

**Figure 1.**
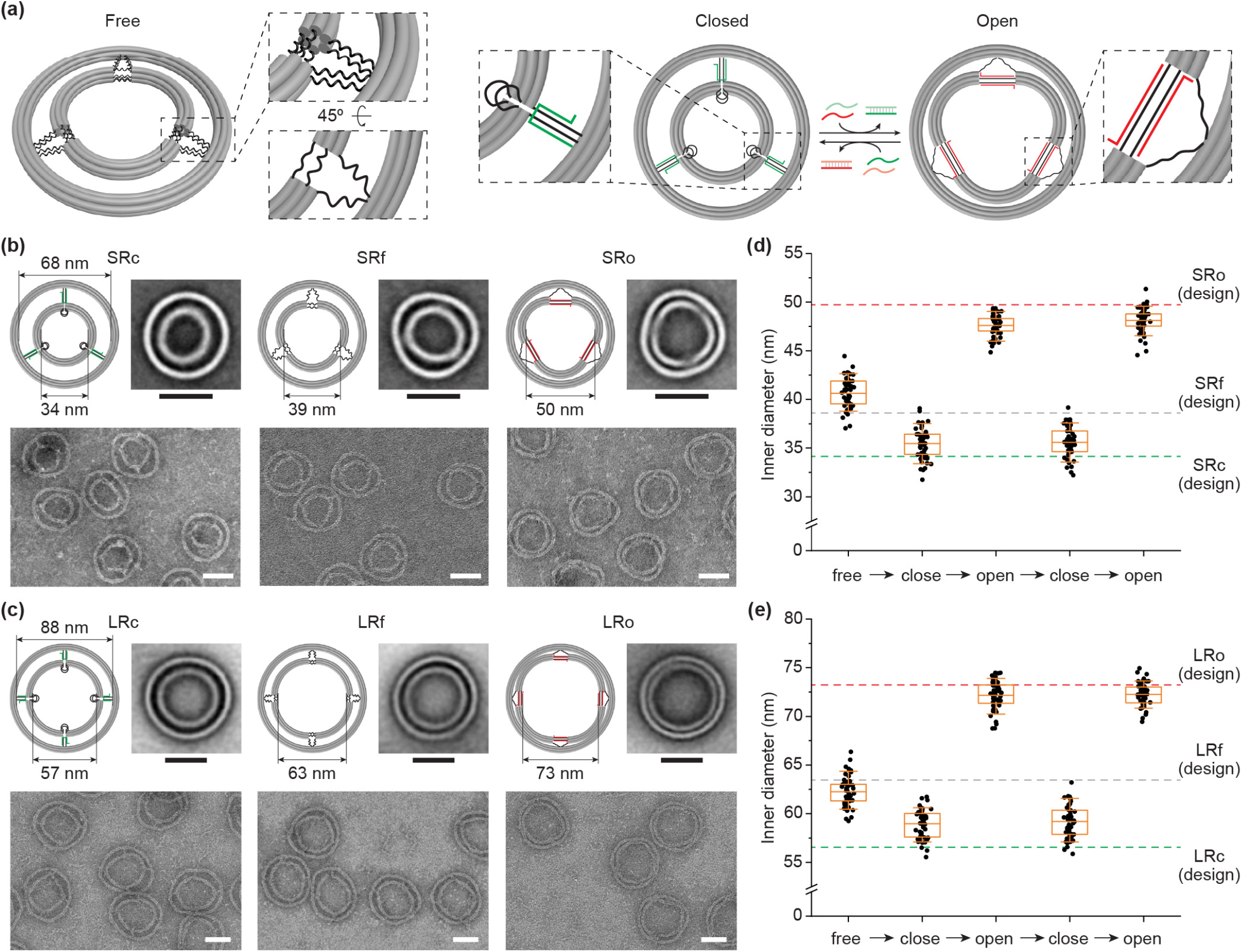
Design and operation of dilatable DNA nanorings. **(a)** Schematics of the dilatable DNA nanoring in the free (left), closed (middle), and open (right) conformations. Reversible switching between the closed and open states is achieved by adding operating strands that zip (dark green) or extend (dark red) the ssDNA tendons (black) and displacement strands (light red and green) that release the counteracting operating strands. **(b)** Negative stain TEM images the small DNA nanoring in three different conformations (SRc, SRf and SRo). Schematics and class-averaged images are shown above the representative micrographs. Scale bars: 50 nm. **(c)** Same as **(b)**, but for the large nanoring. **(d)** Inner ring sizes of the small device during two rounds of reconfiguration measured by TEM (*n* = 50 for each sample). Gray, green, and red dashed lines indicate the designed inner diameters of the free, closed, and open inner ring, respectively. The orange line, box and whisker represent the mean, 25th–75th and 10th–90th percentiles, respectively.

### Dilating DNA nanoring expands liposome

We first used the DNA nanorings in their closed conformation to produce DNA-templated liposomes with defined sizes following an established protocol^22^. Taking the small device as an example (**Figure 2a**), we labeled its inner ring with up to 27 copies of cholesterol via evenly spaced inward-facing DNA handles and similarly grafted up to 16 Alexa Fluor 647 to the exterior of its outer ring (AF647, not shown in **Figure 2** for clarity; see **Figure S1** for full design). After mixing the hydrophobically and fluorescently labeled DNA rings with phospholipids (82.2% DOPC, 15% DOPS, 2% PEG2k-DOPE, 0.8% rhodamine-DOPE, **Table S1**) in the presence of 1% n-octyl-β-D-glucopyranoside (OG), we then dialyzed the mixture overnight against a detergent-free buffer to allow liposome formation and purified the templated SUVs by isopycnic centrifugation. During the detergent removal process, we expected the device’s cholesterol moieties to nucleate lipid bilayer growth and the inner ring to dictate the liposome size, as we had demonstrated in earlier work^22^. Finally, we added appropriate operating strands to dilate the inner ring, pulling on the encircled SUV. Under this condition, we estimated the maximal force sustained by each cholesterol-bearing handle to be 4.6 pN (15.2 pN for the large device) before the device mechanically fails from the buckling of dsDNA struts holding open the ring (**Figure S11**, see “**Maximum forces on DNA handles in open nanorings**” in the Supplementary Materials for details). These values are below or within the range of the forces required to dislodge a DNA handle from the origami structure^32,33^ (>10–30 pN) or to extract a cholesterol molecule from liquid-disordered bilayer membrane^34^ (12–30 pN). On the other hand, the devices’ maximal mechanical work output, or the combined elastic bending energy of the ring-opening struts (∼750 and 1300 kBT for the small and large device, respectively), exceeds the energy needed for the templated SUV to reach its lysing tension (∼15 mN/m, see “**Theoretical work output by dilating DNA nanorings**” in the Supplementary Materials for details). Therefore, we expected the liposome-encircling nanodevices to undergo closed-to-open transition with all handles under mechanical load, leading to substantial membrane expansion.

**Figure 2.**
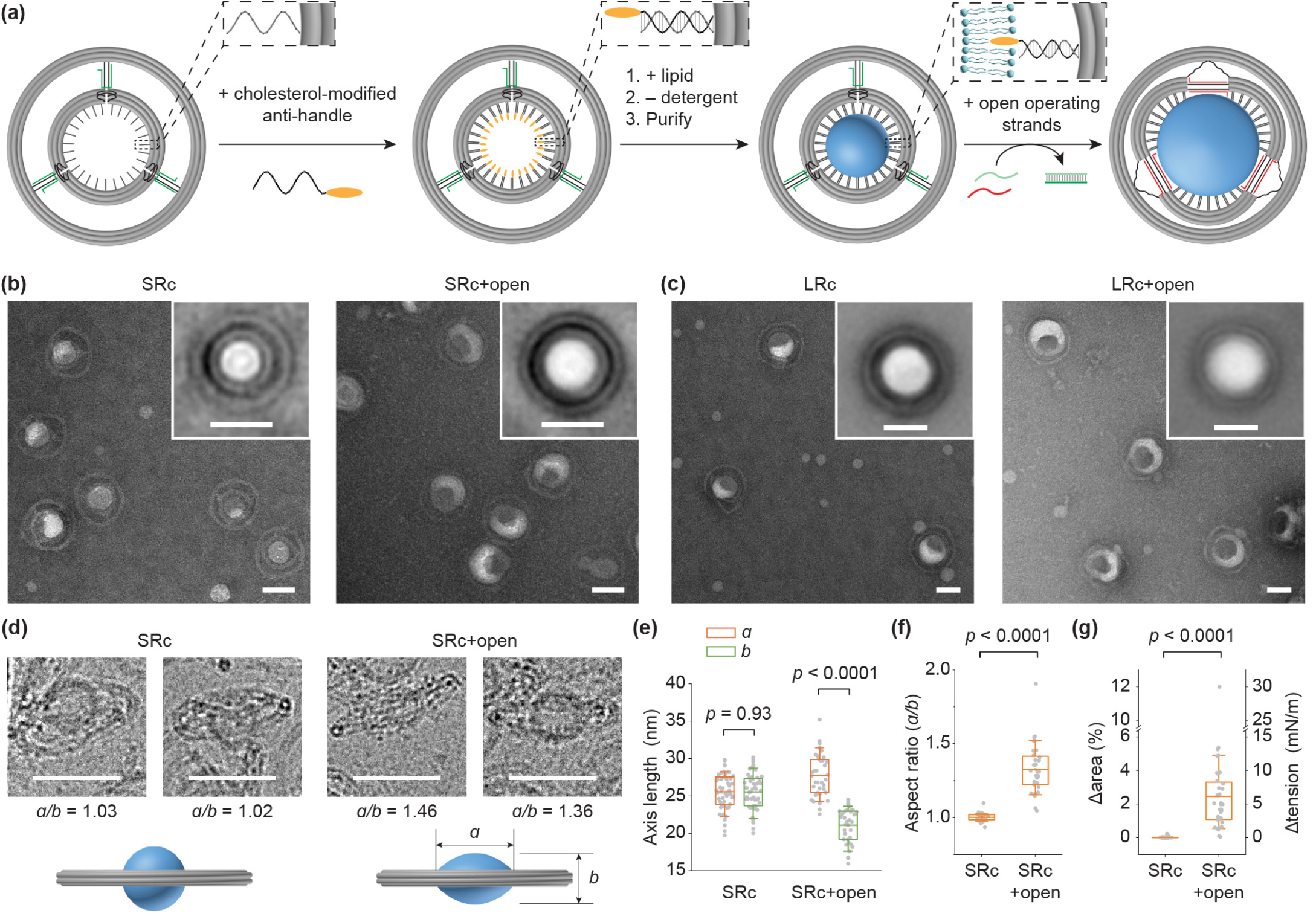
DNA-controlled liposome formation and expansion. **(a)** Schematic of the formation and expansion of DNA-templated liposome using the small device as an example. A constricted DNA nanoring carrying up to 27 (or 56 in the large ring) inward-facing cholesterol moieties (orange ellipse) templates the formation of size-controlled SUVs. Liposome expansion is driven by the subsequent dilation of the inner ring. **(b)** Negative stain TEM images of liposomes in the small DNA nanoring before (SRc) and after ring dilation (SRc+open). Insets are class-averaged TEM images. **(c)** Same as **(b)**, but for liposomes in the large nanoring. **(d)** Representative side views of liposomes encircled by the small nanoring under cryo-EM. The schematics highlight the liposomes’ different aspect ratios before (left) and after (right) ring dilation. **(e)** Dimensions of the liposomes inside the closed (*n* = 52) and open (*n* = 36) small ring measured by cryo-EM. **(f)** Liposome aspect ratio derived from **(e). (g)** Relative changes in liposome surface area and the corresponding increases in membrane tension derived from **(f)**, assuming perfectly spherical liposomes in the tension-free state. Scale bars: 50 nm. The line, box and whisker represent the mean, 25th–75th and 10th–90th percentiles, respectively. *P* values are calculated using an unpaired two-sample t-test with Welch’s correction. Liposomes are made of ∼82.2% 1,2-dioleoyl-sn-glycero-3-phosphocholine (DOPC), 15%1,2-dioleoyl-sn-glycero-3-phospho-l-serine(DOPS),2.0% 1,2-dioleoyl-sn-glycero-3-phosphoethanolamine-N-[methoxy(polyethylene glycol)-2000] (PEG2k-DOPE), and 0.8% 1,2-dioleoyl-sn-glycero-3-phosphoethanolamine-N-(lissamine rhodamine B sulfonyl) (rhodamine-DOPE).

Negative-stain TEM showed that after purification, most closed-state DNA devices held an SUV in the inner ring (**Figures 2b, 2c, S13, S14, S16**). As expected, the templated SUVs were homogenous in size with measured diameters (31.7 ± 2.9 nm in SRc and 52.4 ± 4.1 nm in LRc) slightly below those of the constricted rings. Upon reconfiguration to the open state, the inner rings dilated to nearly the same extent as in the liposome-free devices, accompanied by a corresponding increase of the liposome width (41.4 ± 3.2 nm in SRo and 65.5 ± 4.0 nm in LRo) (**Figures S15, S17, S18**). Most open-state devices kept an SUV in the center, suggesting the force application introduced minimal cholesterol detachment or device damage. However, typical lipid-bilayer membranes can expand only a few percent before rupture, yet the negative-stain TEM observed 20–30% increase in vesicle width without apparent membrane fragmentation. Considering that the negative-stain micrographs granted only the top view of dehydrated liposomes, we examined the small devices by cryo-EM to capture additional view angles of SUVs with well-preserved 3D structure (**Figures 2d, S19, S20**). Remarkably, the spherical liposomes (aspect ratio: 1.00 ± 0.02, *n*=52) in the closed-state devices were significantly flattened (aspect ratio: 1.33 ± 0.16, *n*=36) after DNA ring dilation (**Figure 2e**, see “**Aspect ratio (AR) of liposomes measured by cryo-EM**” in the Supplementary Materials for details). Based on the cryo-EM data, we estimated 0.6–4.9% (mean=2.5%, 10th–90th percentile) membrane expansion caused by pulling on the equatorial plane of vesicles (**Figure 2f**), which corresponds to an increase of membrane tension by 1.4–12.2 mN/m (mean=6.1 mN/m, **Figure 2g**, see “**Calculating membrane tension of expanded liposomes**” in the Supplementary Materials for details). The variation may arise from different membrane tension of SUVs before ring dilation and errors in measuring the dimensions of vesicles that did not give a perfect side view under cryo-EM. Expectedly, the DNA rings and SUVs remained unchanged when treated with an irrelevant DNA strand (a 24-mer poly-thymine or dT24, **Figure S18**). Therefore, the electron microscopy studies unambiguously showed deformed liposomes with expanded membranes following the controlled DNA ring dilation.

### DNA-controlled membrane expansion activates ion channel MscL

To demonstrate the effect of controlled membrane expansion on membrane protein functions, we reconstituted large-conductance mechanosensitive channel (MscL) into SUVs templated by dilatable DNA rings (**Figure 3a**). Functioning as the bacteria’s emergency response to prevent cell membrane lysis under acute turgor pressure increase, activation of MscL opens a non-selective channel as wide as 3 nm and requires extreme membrane tension (10–12 mN/m).^35^ The formation and subsequent stretching of MscL-embedded SUVs in the small and large devices were similar to the protein-free SUVs, as verified by gel electrophoresis (**Figures 3b, S23, S24**) and negative-stain TEM (**Figure S25**). The SUVs, initially formed within constricted rings and loaded with 100 mM calcein (hydrodynamic radius 7.4 Å)^36^, showed nearly constant calcein fluorescence over 23 hours when the devices remained unperturbed in the presence of dT24 (red trace, **Figure 3c**) until vesicles were lysed by 1% OG, causing sudden dilution and fluorescence dequenching of calcein. In contrast, upon adding operating strands to dilate the nanorings, the calcein fluorescence increased steadily for >5 hours, suggesting that increased membrane tension led to gradual release of calcein (black trace, **Figure 3c**). By comparison, DNA-controlled membrane expansion did not result in calcein dequenching for protein-free liposomes (blue trace, **Figure 3c**), confirming that the ion channel activation, rather than membrane rupture, was responsible for the calcein release. For both the small and large devices, calcein fluorescence plateaued at around 60% of the maximum (i.e., detergent-lysed vesicles) following ring dilation (**Figures 3c, S26**); in other words, membrane tension increases triggered MscL gating in most vesicles. Partial opening of the large device by rigidifying 2 or 3 out of the 4 groups of tendons led to smaller fluorescence increments (to ∼10% or 20%, respectively, **Figure S27**), showing that even with reduced energy input, mechanical stress persisted over hours to continuously activate MscL. In contrast, calcein release from MscL-embedded SUVs generally stopped within minutes after hypoosmotic shock due to immediate membrane relaxation after channel opening (**Figure S28**). Furthermore, following the initial DNA ring dilation, we were able to halt/resume calcein release by closing/reopening the DNA device (**Figure 3d**). Finally, to selectively activate MscL embedded in a heterogeneous vesicle population, we mixed calcein-loaded SUVs encircled by the large ring with sulforhodamine B (SRB, hydrodynamic radius 5 Å)^37^ loaded SUVs encircled by the small ring and opened the two devices either independently or concurrently. Release of calcein and SRB, two spectrally distinct fluorescent dyes, strictly followed triggered expansion of rings encircling the dyes’ corresponding SUV containers (**Figure 3e**). Therefore, the DNA devices allowed for programmable, selective, and reversible control of membrane tension.

**Figure 3.**
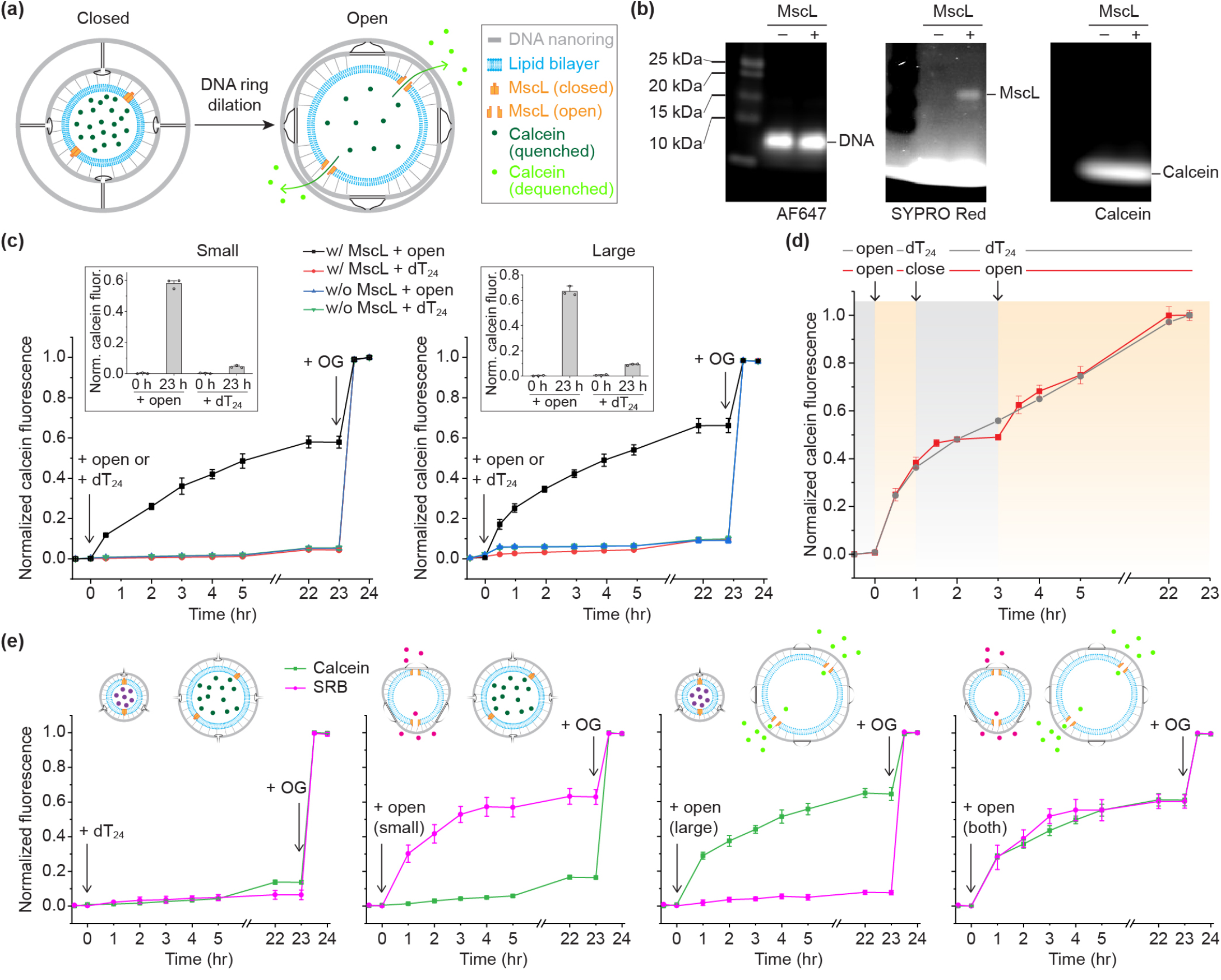
MscL gating triggered by liposome expansion. **(a)** Schematic of the calcein release assay to monitor MscL gating. The release of self-quenched calcein from MscL-embedded liposomes following the membrane expansion triggered by DNA ring dilation is detectable by the increase of calcein fluorescence. **(b)** SDS–PAGE analysis of calcein-loaded liposomes formed in the small DNA device with and without MscL. The gel is imaged in three fluorescent channels: Alexa Fluor 647 (left), SYPRO Red (middle), and calcein (right). **(c)** Traces of calcein fluorescence for liposomes in the small (left) and large (right) nanorings. The ring-opening DNA strands (open) or an irrelevant strand (dT24) are added at time 0 to liposomes with or without MscL. OG (1%) is introduced to all samples after incubating for 23 hr. Fluorescence traces are normalized to the before-reconfiguration (as 0) and after-lysis (as 1) levels. Insets: normalized starting (0 h) and endpoint (23 h) calcein fluorescence. **(d)** Reversible MscL gating monitored by calcein fluorescence. The small DNA device is switched open at time 0 and either closed and reopened (red trace) or treated with dT24 (gray trace) after 1 and 3 hours. For the reversible gating experiment, the periods of time when liposome membranes are expected to be relaxed and expanded are marked by the gray and yellow background, respectively. Fluorescence traces are normalized to the before-reconfiguration level (as 0) and the endpoint (as 1). **(e)** Selective MscL gating in mixed liposomes by independently operated DNA devices. Sulforhodamine B (SRB, loaded in the small device) and calcein (loaded in the large device) fluorescence are monitored after the addition of dT24 (left), ring-open DNA strands targeting the small device (middle left), the large device (middle right), or both devices (right). The plots in (**c**–**e**) show the mean and standard deviation from three technical repeats.

### Heightened membrane tension accelerates SNARE-mediated vesicle fusion

In addition to actuating mechanosensitive membrane proteins, membrane tension is a principal regulator of lipid bilayers’ propensity for remodeling. A typical membrane remodeling event in cells, fusion of vesicles and organelles underlies a broad range of trafficking pathways, nearly all of which require soluble N-ethylmaleimide-sensitive factor attachment protein receptors (SNAREs). Theoretical simulations found that high membrane tension promotes fusion by lowering the initial energy barrier and helping with the subsequent growth of fusion pore^38,39^. Experiments using stretched supported bilayers and surface-adhered GUVs as model membranes corroborated these findings, showing enhanced fusion efficiency when membrane tension increased beyond 4 mN/m.^40^ There, the membranes under tension were virtually flat, mimicking target organelle membranes (e.g., plasma membrane, containing t-SNAREs) rather than vesicles (where v-SNAREs reside). The DNA-ring encircled liposomes are comparable to intracellular vesicles in size and can reach near lytic membrane tension upon expansion, thus enabling us to study the effects of vesicles’ membrane tension on their fusion kinetics.

To this end, we set up ensemble lipid and content mixing assays using tension-controlled SUVs reconstituted with v-SNARE VAMP2 (lipid:SNARE ≈ 200:1) and template-free vesicles with t-SNAREs SNAP25 and syntaxin1 (lipid:SNARE ≈ 400:1). For the lipid-mixing assay, we first formed VAMP2-embedded vesicles in SRc and LRc and doped them with 1.5% NBD-DOPE and 1.5% rhodamine-DOPE (**Figures S29, S30**), creating a FRET pair in the vesicle membranes to quench the NBD fluorescence. We then mixed these SUVs with t-SNARE-embedded liposomes (85% DOPC, 15% DOPS, 45.6 ± 9.7 nm, **Figure S31**) at 37 °C (lipid concentration = 0.3 mM) and monitored the NBD fluorescence. Lipid exchange between liposomes containing complementary SNAREs should separate NBD and rhodamine labeled lipids, leading to increased NBD fluorescence (**Figure 4a**). For both small and large devices, the VAMP2- embedded SUVs within constricted nanorings fused at a rate similar to those without a template (red and purple traces, **Figures 4b, 4c**). Thus, the DNA rings neither promote nor hinder fusion *per se*. However, after dilating the rings to pre-expand VAMP2-SUV membranes, the rate of lipid mixing in the first 4 hours increased by 3–4 folds (black traces, **Figures 4b, 4c**), in line with the theory that heightened vesicle membrane tension accelerates the initial stage of membrane fusion by exposing more hydrophobic lipid acyl chains. This process relied strictly on the formation of trans-SNARE complexes (SNAREpins), as lipid mixing was absent without t-SNAREs (green traces, **Figures 4b, 4c**) or in the presence of the cytosolic domain of VAMP2 (CDV, blue traces, **Figures 4b, 4c**), even after the nanoring dilation. TEM micrographs further illustrated SNAREpin-dependent vesicle interactions, as the apparent docking and merger between ring-encircled vesicles and template-free ones were only found in the presence of all functional SNARE components and without CDV (**Figure S32**). Finally, we probed membrane tension’s effect on SNARE-mediated release of the soluble content in vesicles. For this, we loaded DNA-templated VAMP2-SUVs with 10 mM SRB, such that the self-quenched SRBs would fluoresce upon fusion with t-SNARE-liposomes (**Figure 4d**). Mechanically stressing VAMP2-SUVs by dilating DNA rings caused little change in SRB fluorescence when SNAREpin assembly was inhibited (green and blue traces, **Figures 4e, 4f**), showing good membrane integrity of the tensioned SUVs. For SNARE-mediated vesicle interactions, we again observed faster content mixing when VAMP2-SUVs were under tension (black vs red traces, **Figures 4e, 4f**). Compared to lipid mixing, the tension-modulated effects on content mixing rate were modest (∼2 fold, **Figures 4e, 4f**). We attributed this to quickly reduced membrane tension upon lipid mixing, which tempers the effect of membrane tension on the growth of fusion pore. Indeed, tensioning both t- and v-SNARE bearing liposomes led to an additional ∼1.5-fold increase in the lipid mixing rate (**Figure S33**).

**Figure 4.**
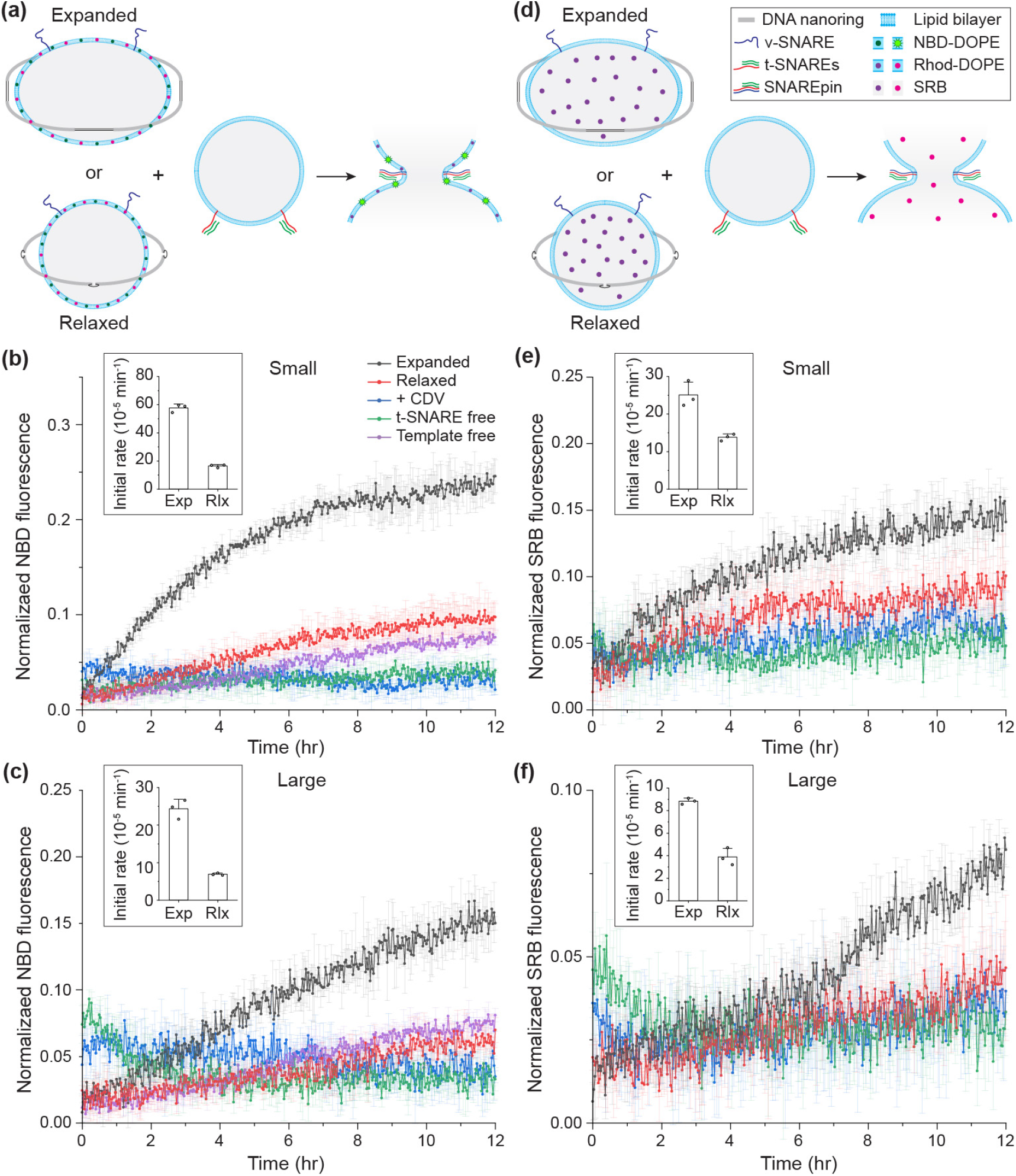
Liposome fusion regulated by membrane tension. **(a)** Schematic of the lipid-mixing assay. Initially quenched NBD dyes (nitrobenzoxadiazole, green dots) fluoresce upon liposome fusion due to reduced FRET with rhodamine dyes (magenta dots). SNARE proteins are depicted as red, green (t-SNAREs) and blue (v-SNARE) curls. Models of liposomes, proteins and dyes are not drawn to scale. Outer rings are omitted for clarity. **(b)** NBD fluorescence traces showing the kinetics of lipid mixing. Liposomes bearing v-SNAREs formed in the small, closed DNA ring, expanded (black) or not (red), are mixed with liposomes bearing t-SNAREs; traces are plotted from the time point when fluorescence increase is detectable (typically ∼30 min after mixing). Addition of CDV (blue) and omission of t-SNAREs (green) serve as controls for expanded v-SNARE liposomes (black), and v-SNARE liposomes formed without DNA template (purple) as a comparison for the templated ones without expansion (red). Inset shows the initial rate of lipid mixing derived from the linear segment of the black (Exp) and red (Rlx) traces. Fluorescence traces are normalized to the measured minimum (as 0) and detergent-lysed (as 1) levels. The plots show the mean and standard deviation from three technical repeats. **(c)** Same as **(b)**, but for v-SNARE liposomes in the large ring. **(d)** Schematic of the content-mixing assay. Self-quenched soluble SRB dyes (magenta dots) fluoresce upon liposome fusion due to dilution. **(e)** and **(f)** are the same as **(b)** and **(c)**, but for the content-mixing assay; traces are plotted from the time point when fluorescence increase is detectable (typically 1– 2 h after mixing).

## Discussion

Controlling the tension of model membranes is fundamental to understanding the influence of membrane mechanics on membrane dynamics and protein activities. Conventional biochemical methods such as lipid extraction and osmotic shock are convenient and scalable but offer limited programmability, whereas existing mechanical tools such as surface expansion and micropipette aspiration are precise but require specialized equipment and do not work for sub-micrometer vesicles. The dilatable DNA-origami rings bridge this technological gap by enabling two-way control over a broad range of membrane tension for sub-100 nm vesicles. Like the current top-down methods, each nanoring stresses and relaxes a vesicle with tunable mechanical energy as it dilates and constricts. However, instead of handling one vesicle at a time, these self-assembled DNA devices work in a massively parallel fashion that make the system amenable to ensemble biochemical analyses, for example the content-release assay for monitoring MscL gating. Using these devices, we find that elevated SUV membrane tension accelerated SNARE-mediated membrane fusion. Recent work showed that purified synaptic vesicles expand nearly 2-fold in membrane area during the uptake of neurotransmitters^41^. Coincidentally, time-resolved cryo-electron tomography on neurons found that large, presumably neurotransmitter-loaded synaptic vesicles rapidly fused with the plasma membrane and shrank to a size equivalent to half of its original membrane area before being recycled^42^.

Taken together, these studies suggest expansion of nanosized vesicles is a mechanism for cells to steer exocytotic fusion, underscoring the physiological relevance of mechanically stressed SUVs as a model system for examining membrane tension regulated fusion kinetics.

The small and large dynamic DNA rings worked on vesicles of different sizes to generate similar effects, showing the versatility of these nanomechanical devices. In this study, we used an archetypical model membrane that mimics the thickness, charge and molecular crowding of plasma membrane^43^. In the future, similar DNA-based devices may be used to control more sophisticated membrane systems including synthetic liposomes and purified vesicles, where membrane tension regulates cooperative actions of multiple protein species, for example the synchronized release of neurotransmitter by synaptic vesicles. Another intriguing application is the structural analysis of proteins and their assemblies with membrane-tension sensitive conformations. Moreover, the ability to differentially stress subpopulations in a mixture of freely diffusing or spatially organized vesicles paves the way to form a membrane tension gradient to investigate bulk lipid transport via non-vesicular pathways^44^.

## Materials and Methods

### Design and assembly of DNA nanorings

DNA nanorings were designed using caDNAno following established principles^31,45,46^ (**Figure S1**). All DNA oligonucleotides were purchased from Integrated DNA Technologies (see **Table S2** for sequence). Unmodified staple strands were supplied in a 96-well plate format with concentrations normalized to 100 µM. Oligonucleotides with fluorescent or cholesterol modifications were HPLC-purified by the vendor. The DNA scaffold strand (8064 nt) was produced using *E. coli* and M13-derived bacteriophages. The small DNA nanoring in the free conformation was assembled from p8064 scaffold strand (50 nM) and a pool of staple strands (300 nM each). The large DNA nanoring in the free conformation was designed to include additional mini-scaffold strands^47^ and assembled from p8064 (20 nM), staple strands (120 nM each), and mini-scaffold strands (240 nM each). All DNA strands were mixed in TE-Mg buffer (5 mM Tris-HCl, 1 mM EDTA, pH 8.0, with 12.5 mM MgCl2) and subjected to a 36-h thermal annealing program: from 80 to 65 °C at ™1°C per 5 min, then from 64 to 24 °C at ™1°C per 50 min, followed by a hold at 12°C. DNA nanorings were purified by rate-zonal centrifugation in glycerol gradients as described previously^48^. After characterized by negative-stain TEM, the purified DNA nanorings were stored at ™20 °C and used within one month. Concentrations of DNA nanorings were determined using a NanoDrop 2000 spectrophotometer (Thermo Fisher). For the initial reconfiguration, DNA nanorings were mixed with appropriate displacement and operating strands at 1:20 molar ratio and incubated at 30°C for 18 h. To induce subsequent conformational changes, counteracting displacement and operating strands were added to achieve 20:1 molar ratio relative to DNA rings after neutralizing the existing displacement and operating strands; the mixture was incubated at 30°C for 1 h.

### Preparing and mechanically stressing DNA nanoring-templated liposomes

Adapting an established protocol^22,23^, unlabeled DNA nanorings were first mixed with cholesterol-modified and AF647-modified DNA strands at handle:anti-handle ratio of 1:1.25 in hydration buffer (25 mM HEPES, 140 mM KCl, 10 mM MgCl2, pH 7.4) containing 1% OG (w/v, 34 mM). The mixture was incubated at 30 °C for 2 h. To form DNA nanoring-templated liposomes, vacuum-dried lipids (Avanti Polar Lipids; composition I, **Table S1**) resuspended in hydration buffer containing 1% OG were added to the cholesterol- and AF647-labeled DNA nanorings (10 nM). The solution was gently shaken at room temperature for 30 min and then transferred to a 7-kDa molecular weight cutoff (MWCO) dialysis cassette (Thermo Fisher) for dialysis against 2 L of hydration buffer for 16 h at 4°C. The dialyzed solutions were subjected to isopycnic ultracentrifugation in iodixanol gradients ranging from 4% to 24% (w/v, 4% intervals, **Figure S13**). Recovered fractions were visualized using a Typhoon FLA 9500 imager (GE Healthcare) to determine their contents. Fractions showing both rhodamine- and AF647-derived fluorescence signals (from liposomes and DNA nanorings, respectively) were combined, characterized by native-stain TEM, stored at 4°C, and used within two days.

To expand liposome membranes, the purified DNA-templated liposomes were mixed with open displacement and operating strands at 1:20 molar ratio and incubated at 30°C for 18 h.

### MscL-gated fluorescent dye dequenching assay

MscL was recombinantly expressed and purified as previously described (**Figure S44**)^49^. DNA nanoring-templated liposomes containing MscL and dye were prepared following the standard protocol described above with slight modifications. Prior to dialysis, purified MscL was added to rehydrated lipids (composition II or III in hydration buffer, **Table S1**) at 1:500 protein-to-lipid molar ratio; calcein disodium salt (VWR) was also added to reach a final dye concentration of 100 mM. The resulting solution underwent a two-step dialysis process at 4°C. First, to remove detergent, the solution was initially dialyzed separately for 16 h against 20 mL of 100 mM calcein in hydration buffer containing 500 mg activated Bio-Beads SM-2 adsorbents (Bio-Rad). Then, to remove unencapsulated calcein, the dialysis cassettes were moved into 1 L of an iso-osmotic dye-free solution (25 mM HEPES, 290 mM KCl, 10 mM MgCl_2_, pH 7.4), with buffer replaced every 12 h for a total of dialysis time of 48 h. SRB (MP Biomedicals, 10 mM) loaded liposomes were made similarly, except an additional 140 mM KCl was introduced to the lipid mixture and the first-step dialysis buffer to compensate for the osmolarity difference with calcein-containing liposomes. DNA-templated liposomes were subsequently purified by isopycnic ultracentrifugation, as described above, and characterized by SDS-PAGE using pre-casted 4–12% gels (Thermo Fisher) or homemade 4%/10% stacking gels stained with SYPRO Red (Thermo Fisher) (**Figures S23, S24**). For fluorescence dequenching analysis, 40 μL of DNA-templated liposomes containing 7 nM for SRc or 1.75 nM for LRc were mixed with open displacement and operating strands or with equivalent molar amount of dT24 at 30 °C for 0.5, 2, 3, 4, 5, 22, and 23 h, at which points fluorescence (excitation/emission: 480/520 nm for calcein, 540/590 nm for SRB) was measured using a BioTek Synergy H1 plate reader (BioTek Instruments). After that, OG was added to a final concentration of 1% to lyse the liposomes, and two additional fluorescence measurements were taken with 30 min intervals.

### SNARE-mediated liposome fusion assay

#### Proteoliposome preparation

SNARE proteins (VAMP2, syntaxin1 and SNAP25) were prepared and purified as previously described^50^. They were reconstituted into DNA-templated liposomes in hydration buffer supplemented with 0.2 mM TCEP, using the standard detergent removal process as described above. Lipid:protein ratios were set at 200:1 for VAMP2-embedded SUVs (composition III for content-mixing assay and IV for lipid-mixing assay, **Table S1**) and 400:1 for t-SNARE-embedded SUVs (composition V, **Table S1**). The resulting DNA nanoring-templated proteoliposomes were purified via isopycnic ultracentrifugation as described above. Recovered fractions were analyzed by SDS-PAGE and visualized using a Typhoon FLA 9500 imager (GE Healthcare) to determine their contents (**Figures S29, S30**). Those containing DNA-templated proteoliopsomes were combined and reconstituted into hydration buffer using Amicon Ultra filtration units (Millipore) with a 30-kDa MWCO. Template-free proteoliposomes were similarly prepared and purified by a 0%/20%/30% (w/v) iodixanol gradient, with the SNARE-embedded liposomes collected from the 0%/20% interface (**Figure S31**).

#### Lipid-mixing assay

DNA nanoring-templated proteoliposomes were expanded by adding the ring-dilating DNA strands as described above. For the relaxed SUV membranes, the correct displacement/operating strands were replaced with equimolar dT24. A typical lipid-mixing assay was initiated by mixing v-SNARE liposomes (lipid concentration: ∼0.03 mM) with t-SNARE liposomes (lipid concentration: ∼0.24 mM) at 37 °C. NBD fluorescence (excitation/emission: 467/540 nm) was recorded every 2 min for 16 h using a Synergy H1 Hybrid Multi-Mode Reader (BioTek Instruments). Then, OG was added to a final concentration of 1% to lyse the liposomes; the maximum fluorescence signal recorded thereafter was used for normalization. Additionally, two negative control experiments were performed using expanded v-SNARE liposomes. In the first control, 10 µM CDV was pre-incubated with v-SNARE liposomes prior to mixing with t-SNARE liposomes, and in the second, t-SNARE liposomes were replaced with protein-free ones of the same lipid concentration. Fluorescence traces were analyzed using Origin (OriginLab).

#### Content-mixing assay

To load liposomes with self-quenching soluble fluorophores, SRB (MP Biomedicals) was added to the rehydrated lipids at a final concentration of 10 mM prior to dialysis. The resulting mixture was subjected to a two-step dialysis procedure. First, to remove detergent, the solution was initially dialyzed for 16 h against 20 mL of an iso-osmotic buffer (25 mM HEPES, 130 mM KCl, 10 mM MgCl_2_, pH 7.4) containing 10 mM SRB and 500 mg activated Bio-Beads SM-2 adsorbents. Then, to remove unencapsulated SRB, the dialysis cassette was moved into 1 L of hydration buffer, with buffer replaced every 12 h for a total dialysis time of 48 h. To measure membrane-fusion induced content mixing, the experimental setup of the lipid-mixing assay was used, except for the fluorescence wavelengths (excitation/emission: 540/590 nm).

### Negative-stain TEM

For DNA-only samples, a drop of the sample (5 µL) was deposited on a glow discharged formvar/carbon-coated copper grid (Electron Microscopy Sciences or Ted Pella), incubated for 1 min, and blotted away. The grid was first rinsed twice with 5 µL of TE buffer containing 12.5 mM MgCl2, washed briefly with 5 µL of 2% (w/v) uranyl formate, and stained for 1 minute with 5 µL of 2% uranyl formate. For samples containing liposomes, a drop of the sample (5 µL) was deposited on a glow discharged formvar/carbon-coated copper grid, incubated for 2 min, and blotted away. The grid was then rinsed with 2% uranyl formate for 10 seconds and stained with 2% uranyl formate for 1 minute. TEM images were acquired on a JEOL JEM-1400Plus microscope (acceleration voltage: 80 kV) with a bottom-mount 4k×3k CCD camera (Advanced Microscopy Technologies). Negative stain 2D class averages were computed using EMAN2.^51^

### Cryo-EM

DNA-templated SUV samples for cryo-EM were first concentrated to ∼40 nM using Amicon Ultra filtration units (Millipore) with a 30 kDa MWCO; iodixanol was removed concurrently. A drop of 4 µL of the concentrated SUVs was applied to a glow discharged lacey carbon 300 mesh grid (Quantifoil Micro Tools) and blotted on a Mark IV Vitrobot (Thermo Fisher Scientific) with blotting time of 4 s and blot force of -4 at 10°C and 100% humidity. For template-free t-SNARE-liposomes, 3 µL of sample was deposited and blotted with force of -1 at 8°C. The cryo-EM images were then taken on a 200 kV Glacios TEM (Thermo Fisher Scientific) with a K3 direct detection camera (Gatan). For DNA-templated SUV samples, images were collected at 45,000× magnification (physical pixel size of 0.86 Å) with a total exposure dose of 50 e^-^/Å^2^; the cryo-EM micrographs were motion corrected with IMOD.^52^ For t-SNARE liposome samples, images were collected at 13,500× magnification (physical pixel size of 3.25 Å) with a total exposure dose of 12 e^-^/Å^2^; the cryo-EM micrographs are motion corrected with MotionCor2.^53^

## Acknowledgment

We thank J.E. Rothman, Y. Zhang, and M. Wu for helpful discussions. This work is supported by National Institutes of Health grants R35-GM149264, R21-GM146105, R21-GM141669 to C.L., R01-AI162260 to C.L. and Y.X, and R35-GM164438-01, RM1-GM149406, and R01-GM141192 to K.G.

## Author contributions

L.L. initiated the project, designed and performed most of the experiments, analyzed data, and prepared the manuscript. E.C. designed the prototype of the small DNA nanoring. C.W., A.R. and S.G. performed the cryo-EM characterization. M.B. prepared SNARE proteins. A.P. prepared MscL under the supervision of K.G. Y.X. interpreted data and supervised C.W. F.P. analyzed and interpreted the membrane tension and liposome fusion data. C.L. initiated the project, designed and supervised the study, interpreted data, and prepared the manuscript. All authors reviewed and approved the manuscript.

## Supplementary Materials Available

Notes, Figures S1–S44, and Tables S1 and S2.

## Data and materials availability

All data needed to evaluate the conclusions in the paper are present in the paper and/or the Supplementary Materials.

