## Supplementary Materials for "Controlling membrane tension by dynamic DNA nanorings"

### NOTES

#### Prediction of ‘free’-state DNA nanoring dimensions

In the free conformation, the inner ring geometry is governed by the tension exerted by ssDNA segments (tendons) of the DNA scaffold strands. We thus evaluated the equilibrium of mechanical forces acting on each arc domain of the inner ring to derive its resting diameter. For both small and large rings, each arc is tethered by 12 ssDNA tendons, 8 of which exert inward-directed force while the other 4 generate outward expansion force (**Figures S9 and S10**). At mechanical equilibrium, these tensile forces balance out, resulting in stable ring geometry with predictable inner diameters (39 nm for SRf and 63 nm for LRf).

The tension exerted by the ssDNA strings was estimated based on the worm-like-chain (WLC) model<sup>1</sup> as

$$F_{WLC}(x) = \frac{k_B \cdot T}{L_{pss}} \cdot \left[ \frac{1}{4 \cdot \left(1 - \frac{x}{L_C}\right)^2} - \frac{1}{4} + \frac{x}{L_C} \right]$$

where $k_B$ is the Boltzmann constant, $T$ is the temperature (297 K), $L_{pss}$ is the persistence length of ssDNA, $L_C$ is the contour length of the ssDNA strings, and $x$ is the extension of the ssDNA strings.

The following Python script (v 3.8 with SciPy<sup>2</sup>) calculates the net radial force on an arc as a function of the angle $\theta$ formed by the arc-to-outer-ring tendon and the radial direction (0°–30°) for the SRf (**Figure S9**). The numerical value of $\theta$ corresponding to zero net force is used to derive the inner diameter (**Figure S9**).

```
from numpy import sin, cos, pi

# All constants are listed below

Lss = 0.63E-9 # Contour length of ssDNA (m/base)
kb = 1.38E-23 #Boltzmann constant (J/K)
T = 297 # Temperature (K)
Lpss = 1.05E-9 # persistence length of ssDNA (m)

# All adjustable parameters are listed below

N1 = 52 # number of nucleotides in ssDNA tether type 1
N1ss = 4 # number of ssDNA tether type 1
N2 = 32 # number of nucleotides in ssDNA tether type 2
N2ss = 2 # number of ssDNA tether type 2
theta_step = 0.1*pi/180 # increase step for theta (0.1 degree)

# Worm-Like Chain Model predicts the force exerted by a ssDNA tether as a function of its end-to-end
distance "x" and number of bases "N"

def F(x,N):
    return((kb*T/Lpss)*(1/(4*((1-x/(N*Lss))**2))-0.25+x/(N*Lss)))

def x(theta):
    return(2*10.1956E-9*sin(theta)/cos(pi/6-theta))

def y(theta):
    return(10.1956E-9/cos(pi/6-theta))

for i in range(300):
    theta = i * theta_step
    print(i/10,N2ss*F(y(theta),N2)*cos(pi/3+theta)-N1ss*F(x(theta),N1)*cos(pi/6))
```

Similarly, the theoretical value of $\theta$ at equilibrium is determined using the following Python script and used to derive the inner diameter of the large DNA ring (LRf, **Figure S10**).

```
from numpy import sin, cos, pi

# All constants are listed below

Lss = 0.63E-9 # Contour length of ssDNA (m/base)
kb = 1.38E-23 #Boltzmann constant (J/K)
T = 297 # Temperature (K)
Lpss = 1.05E-9 # persistence length of ssDNA (m)

# All adjustable parameters are listed below

N1 = 40 # number of nucleotides in ssDNA tether type 1
N1ss = 4 # number of ssDNA tether type 1
N2 = 28 # number of nucleotides in ssDNA tether type 2
N2ss = 2 # number of ssDNA tether type 2
theta_step = 0.1*pi/180 # increase step for theta (0.1 degree)

# Worm-Like Chain Model predicts the force exerted by a ssDNA tether as a function of its end-to-end
distance "x" and number of bases "N"

def F(x,N):
    return((kb*T/Lpss)*(1/(4*((1-x/(N*Lss))**2))-0.25+x/(N*Lss)))

def x(theta):
    return(2*9.7060E-9*sin(theta)/cos(pi/4-theta))

def y(theta):
    return(9.7060E-9/cos(pi/4-theta))

for i in range(450):
    theta = i * theta_step
    print(i/10,N2ss*F(y(theta),N2)*cos(pi/3+theta)-N1ss*F(x(theta),N1)*cos(pi/6))
```

#### Maximum forces on DNA handles in open nanorings

The Euler's buckling load<sup>3</sup> of the DNA struts connecting adjacent arcs provides the upper bound for force applied to the lipid membrane by a nanoring through the DNA handles. The critical buckling force of a DNA strut is calculated using the function:

$$F_{buckling} = \pi^2 \times \frac{k_B T l_p}{L^2}$$

where $k_B$ is the Boltzmann constant, $T$ is the temperature (297 K), $l_p$ is the persistence length of the DNA strut, $L$ is the strut length. Here we use $l_p = 50$ nm for double-stranded DNA (dsDNA).<sup>4</sup>

In SRO, the neighboring arcs are connected by 4 independent 52-bp DNA duplexes ($L=17.42$ nm). The strut buckling force is thus $4\times$ that of an individual DNA duplex ($6.7$ pN) or $26.8$ pN. Assuming the system is at equilibrium right before the struts buckle and each of the 9 DNA handles along the arc bears the same load ($F_{handle}$), we thus obtained $F_{handle} = 4.6$ pN (**Figure S11a**), using the following equation:

$$2 \times F_{buckling} \times \cos 30^\circ = 4 \times F_{tendon} \times \cos 67.7^\circ + \sum_{i=1}^9 F_{handle} \times \cos \alpha_i$$

where $F_{buckling} = 26.8 \text{ pN}$, $F_{tendon} = 5.9 \text{ pN}$ (calculated using the WLC model), $\alpha_i = [40.6^\circ, 30.2^\circ, 20.0^\circ, 9.9^\circ, 0^\circ, -9.9^\circ, -20^\circ, -30.2^\circ, -40.6^\circ]$.

In LRo, each DNA strut is a 13.4-nm-long (40 bp) four-helix bundle (**Figure S1b**). The persistence length of the strut is estimated based on the area moment of inertia relative to dsDNA (**Figure S12**), yielding a critical buckling force of $140.5 \text{ pN}$. Assuming the system is at equilibrium right before the struts buckle and each of the 14 DNA handles along the arc bears the same load ($F_{handle}$), we thus obtained $F_{handle} = 15.2 \text{ pN}$ (**Figure S11b**), using the following equation:

$$2 \times F_{buckling} \times \cos 45^\circ + 4 \times F_{tendon} \times \cos 88.6^\circ = \sum_{i=1}^{14} F_{handle} \times \cos \alpha_i$$

where $F_{buckling} = 140.5 \text{ pN}$, $F_{tendon} = 6.0 \text{ pN}$, $\alpha_i = [34.2^\circ, 28.8^\circ, 23.5^\circ, 18.2^\circ, 13.0^\circ, 7.8^\circ, 2.6^\circ, -2.6^\circ, -7.8^\circ, -13.0^\circ, -18.2^\circ, -23.5^\circ, -28.8^\circ, -34.2^\circ]$.

#### Theoretical work output by dilating DNA nanorings

We assumed the maximal mechanical work ($W$) responsible for membrane expansion is the combined elastic bending energy of the ring-opening struts, which we calculated using a toy model<sup>5</sup>:

$$W = E_{DNA\_bending} = N \times \sum \frac{1}{2} \times B \times \frac{L}{r^2}$$

where $B$ is the bending modulus of dsDNA ($230 \text{ pN} \cdot \text{nm}^2$), $L$ is strut length, $r$ is the radius of a strut bent to $360^\circ$, and $N$ is the total number of arc-bridging DNA duplexes in an inner ring (12).

Therefore, the dilating DNA nanoring devices can theoretically perform mechanical work up to $3127 \text{ pN} \cdot \text{nm}$ (or $763 k_B T$, small ring) and $5421 \text{ pN} \cdot \text{nm}$ (or $1322 k_B T$, large ring).

We then compared $W$ output by the DNA devices to the energy needed to expand the encircled SUVs ($E_{membrane\_exp}$). Taking the small DNA nanoring as an example, the surface area ($S$) of the liposome before DNA ring dilation can be calculated using the function:

$$S = 4\pi R^2 = 2055.7 \text{ nm}^2$$

Where $R$ is the cryo-EM-measured average radius (12.79 nm) of the liposomes before DNA ring dilation.

The membrane tension ($\gamma$) can be described using the function:

$$\gamma(\Delta S) = \kappa \times \frac{\Delta S}{S}$$

Where $\Delta S$ is the change in membrane surface area, $\kappa$ is the elastic modulus ($250 \text{ mN/m}$ for DOPC)<sup>6,7</sup>.

Energy ($E_{membrane\_exp}$) required to achieve a final membrane surface area change ($\Delta S_f$) can be described using the function:

$$E_{membrane\_exp} = \int_0^{\Delta S_f} \gamma(\Delta S) d(\Delta S) = \int_0^{\Delta S_f} \kappa \times \frac{\Delta S}{S} d(\Delta S) = \frac{1}{2} \times \kappa \times \frac{\Delta S^2}{S} \Big|_0^{\Delta S_f}$$

To reach the lytic membrane tension (assuming a lytic tension of $\gamma(lysis) = 15 \text{ pN/nm}$), the required surface area change can be calculated using:

$$\Delta S_f = \frac{\gamma(lysis) \times S}{\kappa} = 123.3 \text{ nm}^2$$

The corresponding mechanical work required to lyse the membrane can then be calculated as:

$$E_{lysis} = \frac{1}{2} \times \kappa \times \frac{\Delta S^2}{S} \Big|_0^{\Delta S_f} = 924 \text{ pN} \cdot \text{nm}$$

Similarly, the mechanical work required to lyse the membrane of the large DNA nanoring-templated liposome is calculated to be $2817 \text{ pN} \cdot \text{nm}$. Note that the radius (22.32 nm) of the liposomes in LRc was estimated by subtracting the DNA handle length from the average inner radius of the templating inner ring.

The calculated work outputs of both devices exceed the mechanical work required for membrane lysis. In practice, however, the actual work output would be smaller than the theoretical maximum and only part of that would be used for membrane expansion.

#### Aspect ratio (AR) of liposomes measured by Cryo-EM

Since a perfect side view of liposomes is rarely captured by cryo-EM, meaning the DNA rings are usually not perfectly perpendicular to the imaging plane, the liposome minor axis ($b_{lipo}$) directly measured from the micrographs would be an overestimation if uncorrected. To better estimate the AR of liposomes, we calibrated the view angle using the DNA outer ring as a reference (**Figure S21**). Briefly, assuming that the DNA outer ring is an ideal circle and that the angle it forms with the EM camera is $\beta$, we obtain:

$$\frac{a'_{DNA}}{b'_{DNA}} = \frac{1}{\cos \beta}$$

where $a'_{DNA}$ and $b'_{DNA}$ are the observed major and minor axis lengths of the outer DNA ring, respectively.

Assuming the liposome under tension adopts the shape of a spherical lens (**Figure S22**), and noting that the orthographic projection of a spherical lens produces projected principal axes that can be well approximated by the corresponding ellipsoidal relation, we treated expanded liposomes as oblate spheroids for the purpose of calculating AR, with $a_{lipo}$ and $b_{lipo}$ denoting the liposome's major and minor axes, respectively. The liposome's actual major axis ($a_{lipo}$) is thus equal to that of the liposome's projection ($a'_{lipo}$), whereas the liposome's actual minor axis ($b_{lipo}$) can be calculated using the function:

$$b'_{lipo} = \sqrt{b_{lipo}^2 \times \sin^2 \beta + a_{lipo}^2 \times \cos^2 \beta}$$

where $a'_{lipo}$ and $b'_{lipo}$ are the liposome's major and minor axis lengths observed by cryo-EM, i.e., those of the projected ellipse.

The liposome's actual AR can then be obtained using the function:

$$AR = \frac{a_{lipo}}{b_{lipo}} = \frac{a'_{lipo} \times \sin \beta}{\sqrt{b'^2_{lipo} - a'^2_{lipo} \times \cos^2 \beta}}$$

#### Calculating membrane tension of expanded liposomes

The membrane tension was calculated based on the increase in liposome surface area, assuming that the liposome volume remains constant after the DNA nanoring dilation. Liposome models shown in **Figure S22** were used to determine the surface area and volume. Before expansion, treating the liposome as a sphere, its initial surface area $S_i$ and volume $V_i$ can be calculated using the functions:

$$S_i = 4\pi R_i^2$$

$$V_i = \frac{4}{3}\pi R_i^3$$

where $R_i$ is the radius of the liposome.

After liposome expansion, treating it as a spherical lens, its final surface area $S_f$ and volume $V_f$ can be calculated using the functions:

$$S_f = 4\pi R_f n = 2\pi(m^2 + n^2)$$

$$V_f = \frac{2}{3}\pi n^2(3R_f - n) = \frac{\pi}{3}n(3m^2 + n^2)$$

Where $R_f$ is the equivalent radius of the spherical lens, and $m$ and $n$ are the semi-major and semi-minor axes, respectively, i.e., $m = a_{lipo}/2$; $n = b_{lipo}/2$.

Assuming that the volume remains unchanged after DNA nanoring dilation ($V_i = V_f$) and using calibrated $a_{lipo}$ and $b_{lipo}$ (see the section above), we can derive the initial radius ($R_i$) of an expanded liposome, hence its initial surface area ($S_i$) before expansion. The membrane tension is then determined using the function:

$$\gamma = \kappa \times \frac{S_f - S_i}{S_i}$$

For comparison, we did the same exercise for liposomes in the closed ring (**Figure 2g**).

#### Initial rate of membrane fusion

In the first 0.5–2 hrs after mixing v-SNARE and t-SNARE bearing SUVs, NBD and SRB fluorescence fluctuates (even for the negative controls), likely due to sample handling. Fluorescence traces measuring vesicle fusion were thus plotted from the time point when fluorescence increase became detectable for both expanded and relaxed membranes. To represent the membrane fusion rate, we applied linear regression to the lipid-mixing traces (**Figures S34–S37**) and content-mixing traces (**Figures S38–S43**). For the initial segments that fit linearly (corrected $R^2$: $\sim 0.5$–$0.9$, $p < 10^{-10}$), their slopes were plotted in **Figure 4** as the initial lipid/content mixing rates. All curve fitting was performed using Origin (OriginLab).

### SUPPLEMENTARY FIGURES

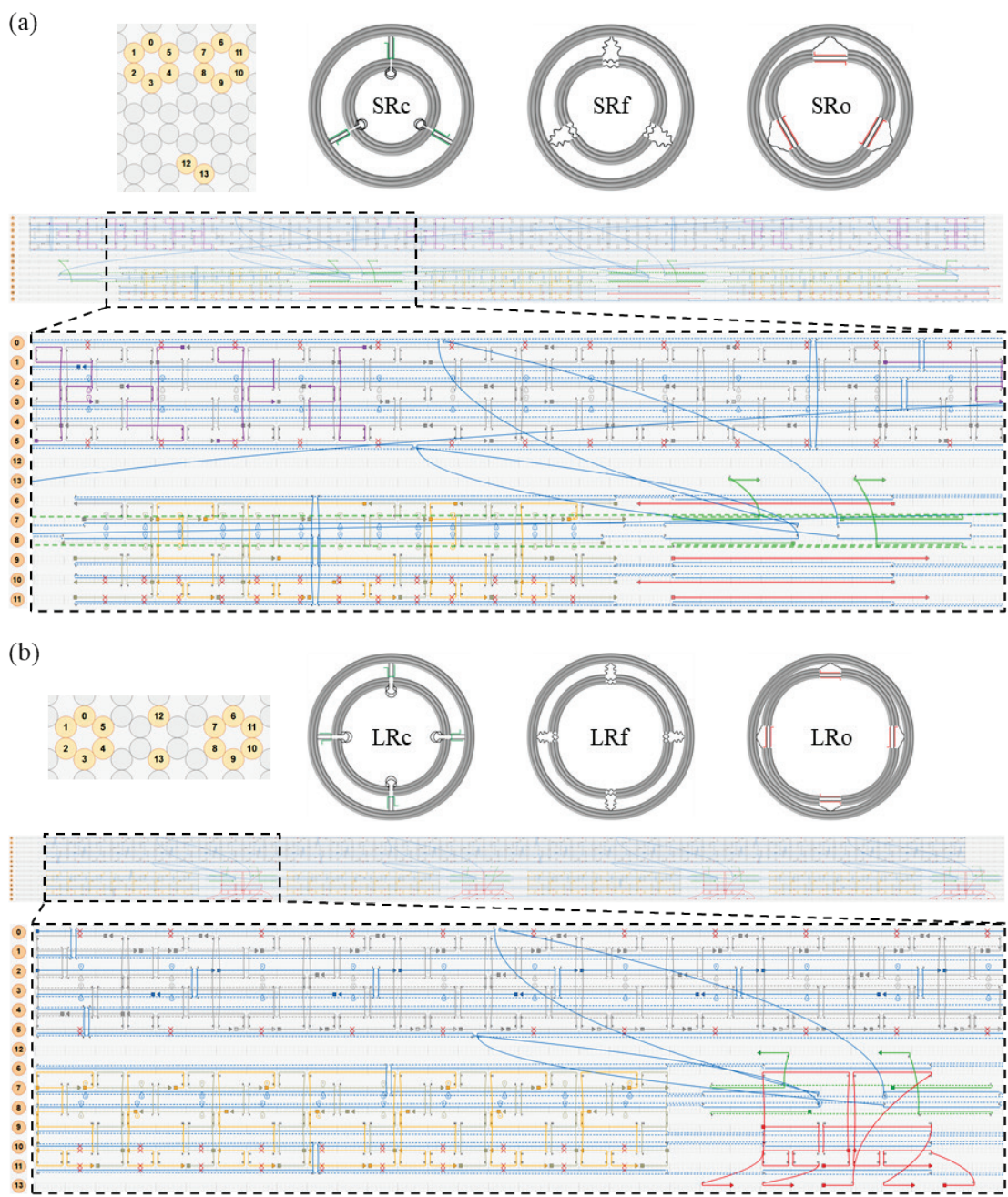

**Figure S1. caDNAno design diagrams of (a) small DNA nanorings and (b) large DNA nanorings.** Orange-colored strands: 3' end extended with the sequence “AAATTATCTACCACAACTCAC” (H1) for hybridizing with cholesterol-modified anti-handles for liposome templating. In small DNA nanorings, purple-colored strands, and in large DNA nanorings, blue-colored strands on helix 2: 3' end extended with the sequence “CTTCACACCACACTCCATCTA” (H2) for hybridizing with Alexa Fluor 647-labeled anti-handles. The conformational states are encoded by the presence or absence of specific staple strands (operating strands): green staples are present only in the closed conformation, red staples are introduced only in the open conformation, and both are absent in the free conformation.

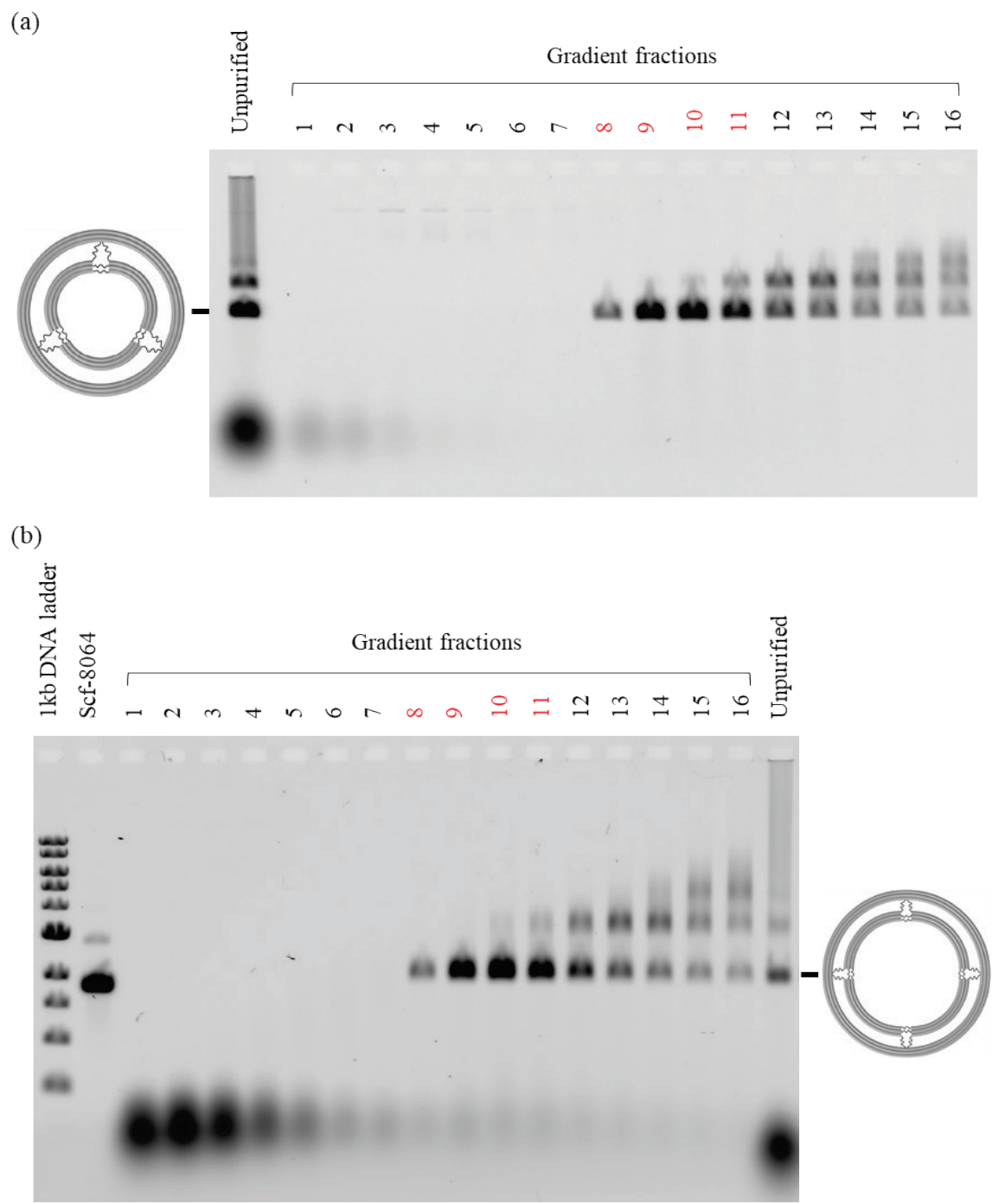

**Figure S2. Agarose gel (1%) analysis of (a) SRf and (b) LRf purified by glycerol-gradient rate-zonal ultracentrifugation. Fractions 8–11 were collected.**

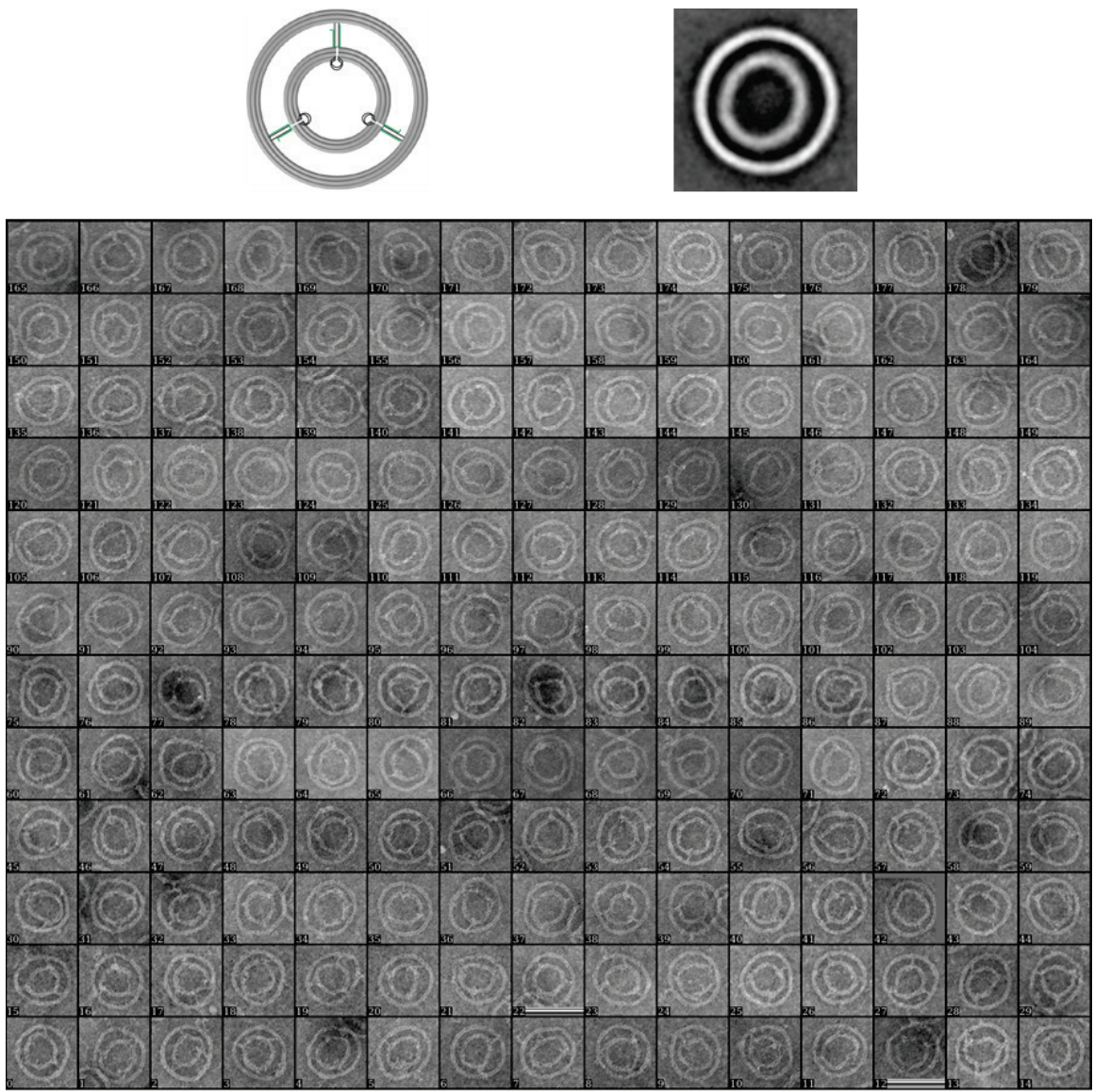

**Figure S3. Class averaging of 180 particles for SRc. Box size: 103 nm.**

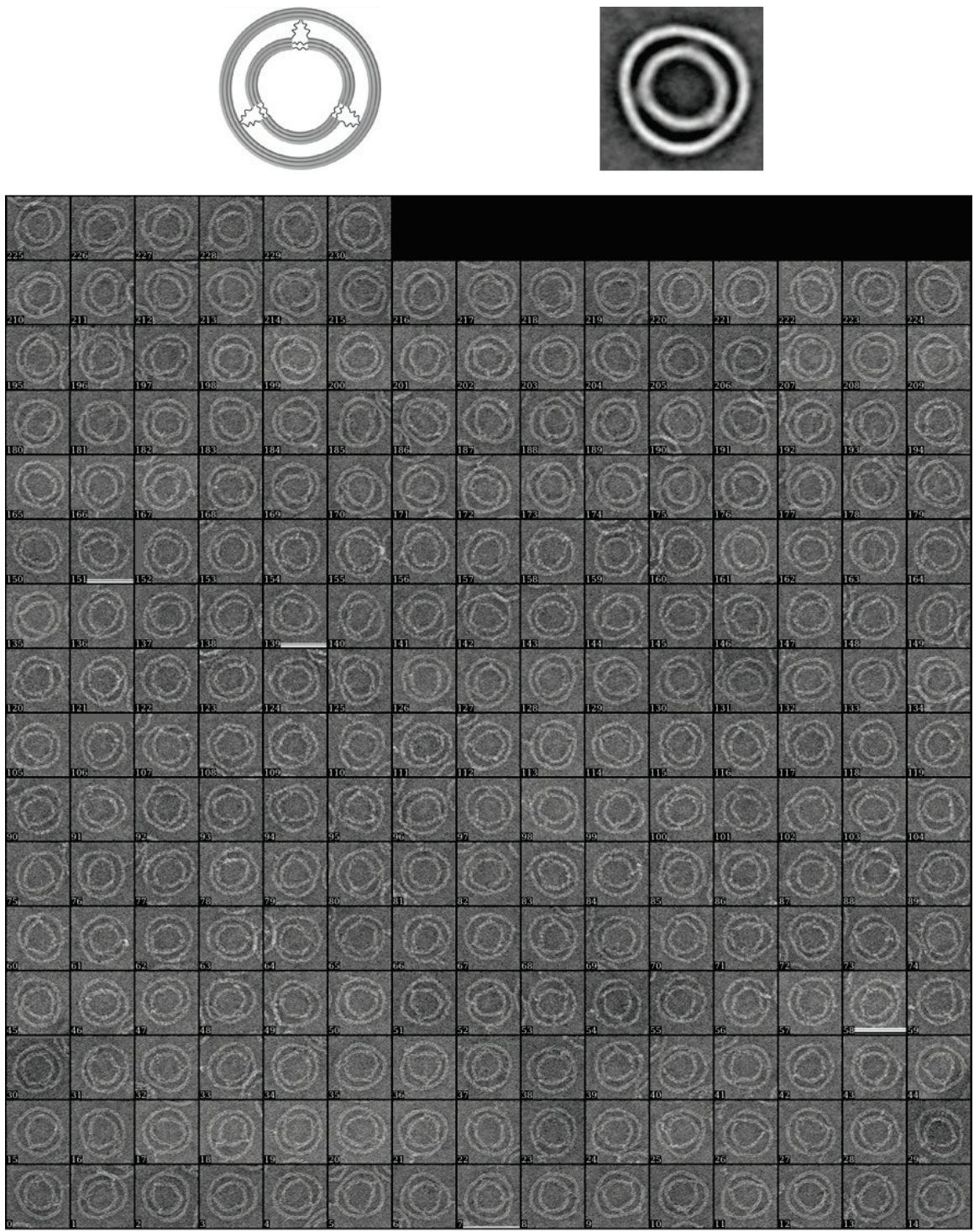

**Figure S4. Class averaging of 231 particles for SRf. Box size: 103 nm.**

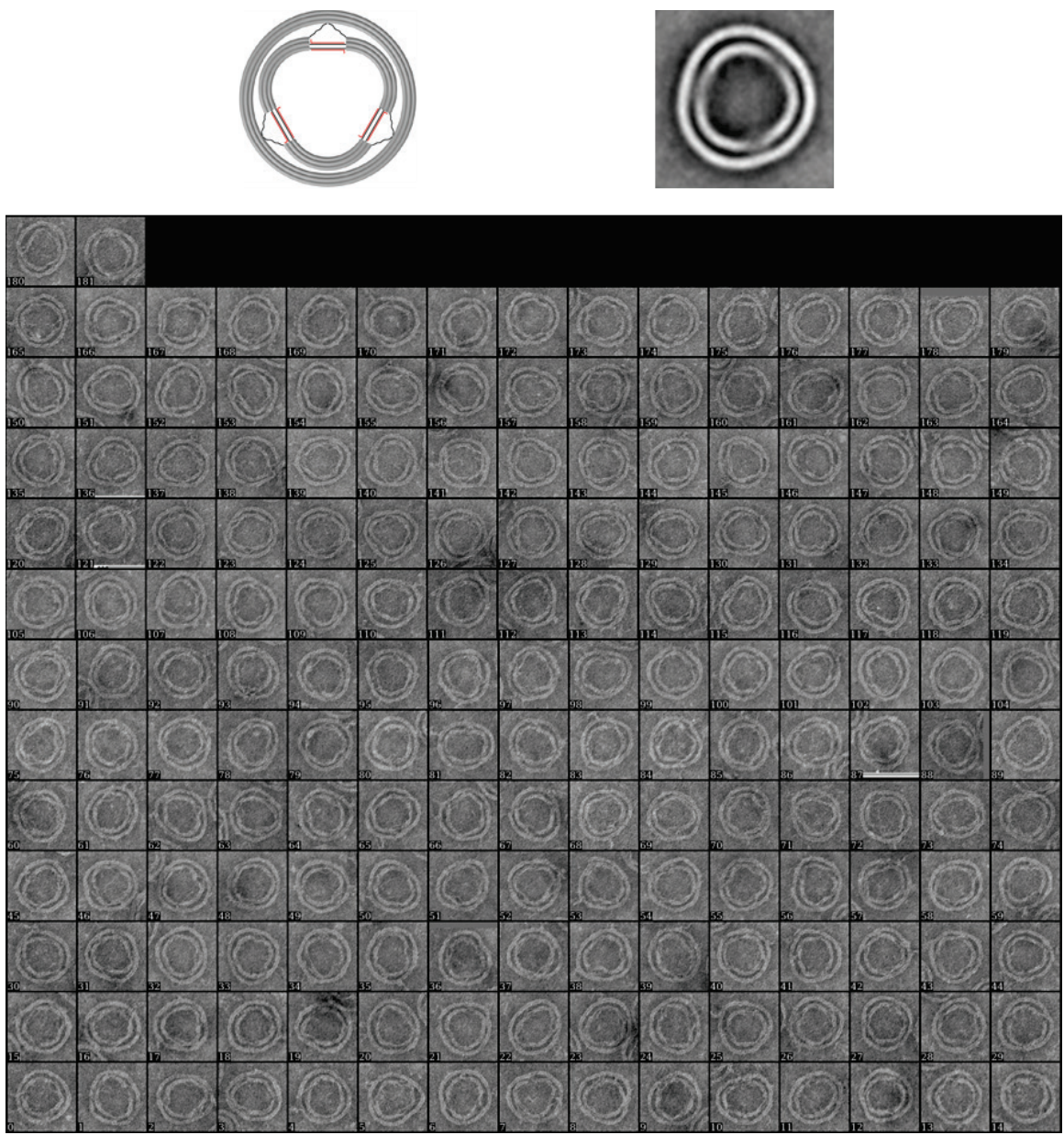

**Figure S5.** Class averaging of 182 particles for SRo. Box size: 103 nm.

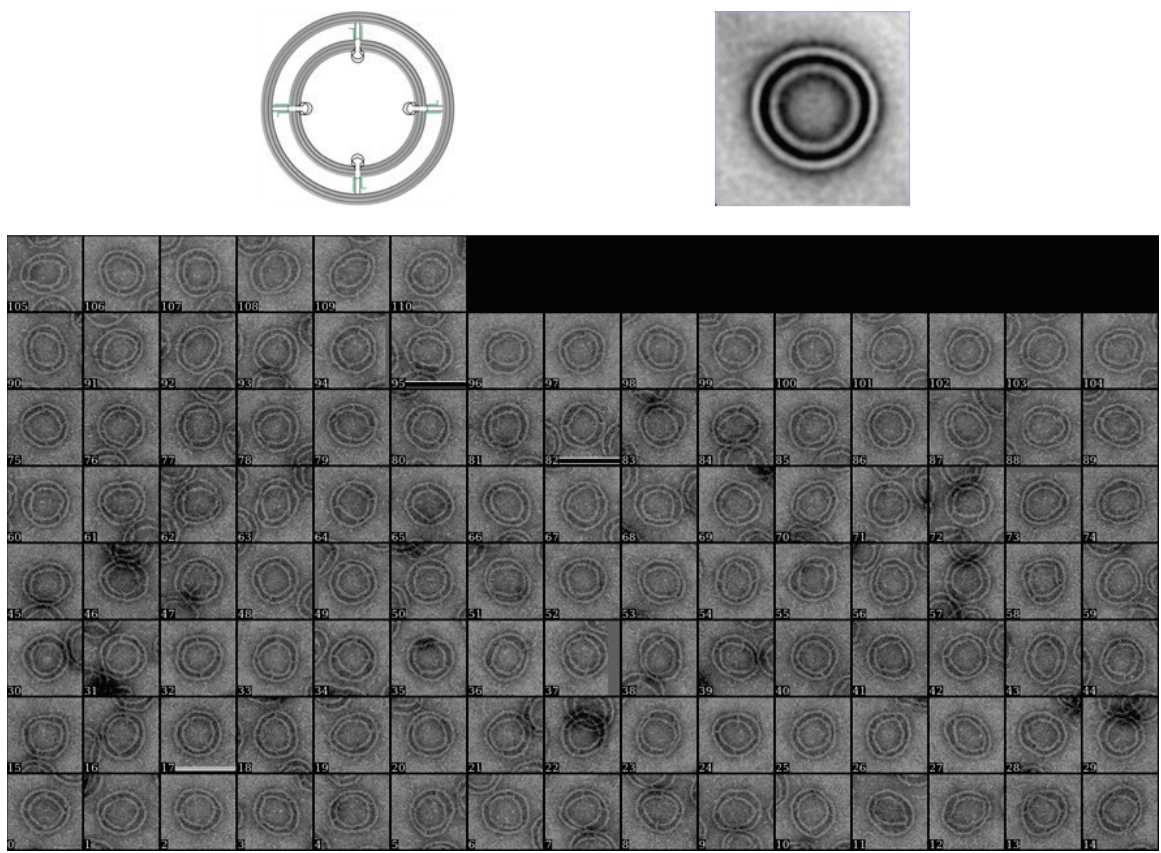

**Figure S6. Class averaging of 111 particles for LRc. Box size: 156 nm.**

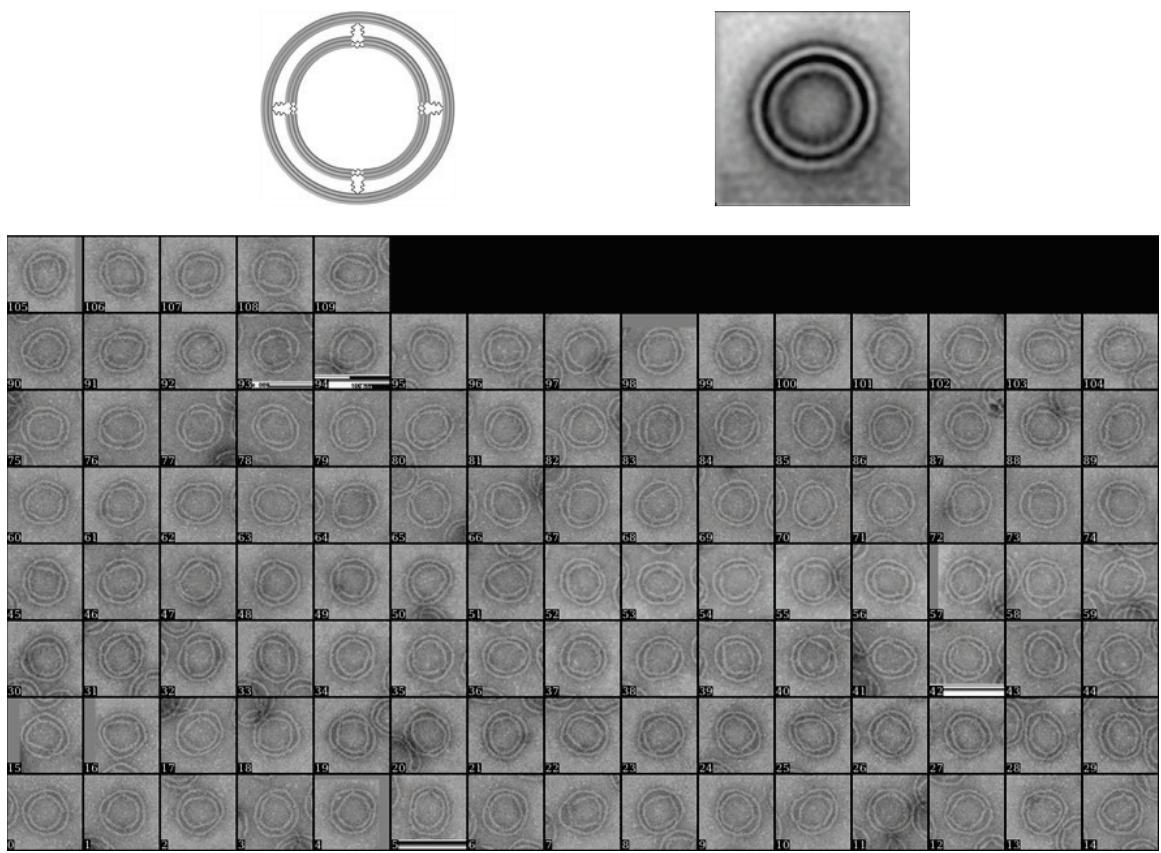

**Figure S7. Class averaging of 110 particles for LRF. Box size: 156 nm.**

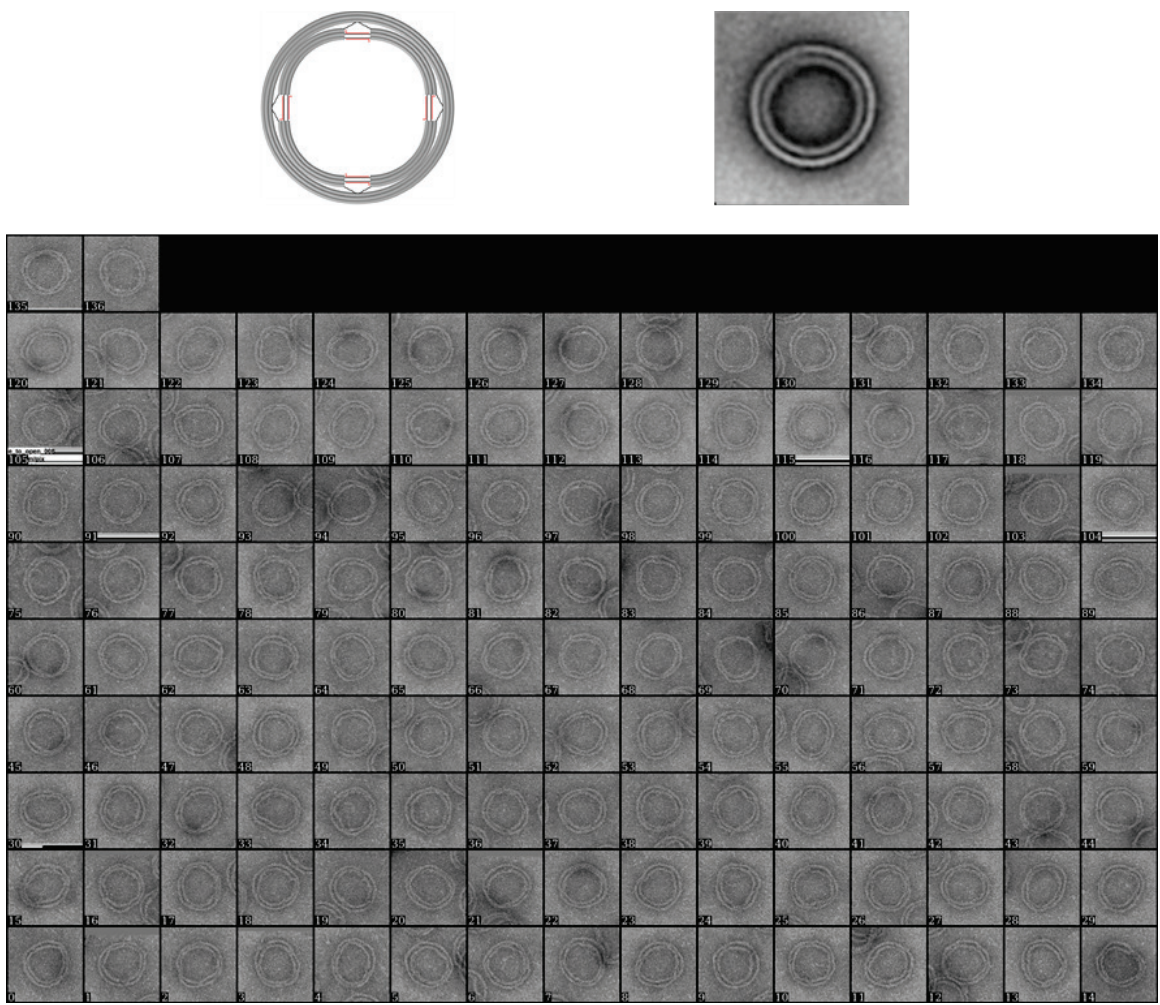

**Figure S8. Class averaging of 137 particles for LRo. Box size: 156 nm.**

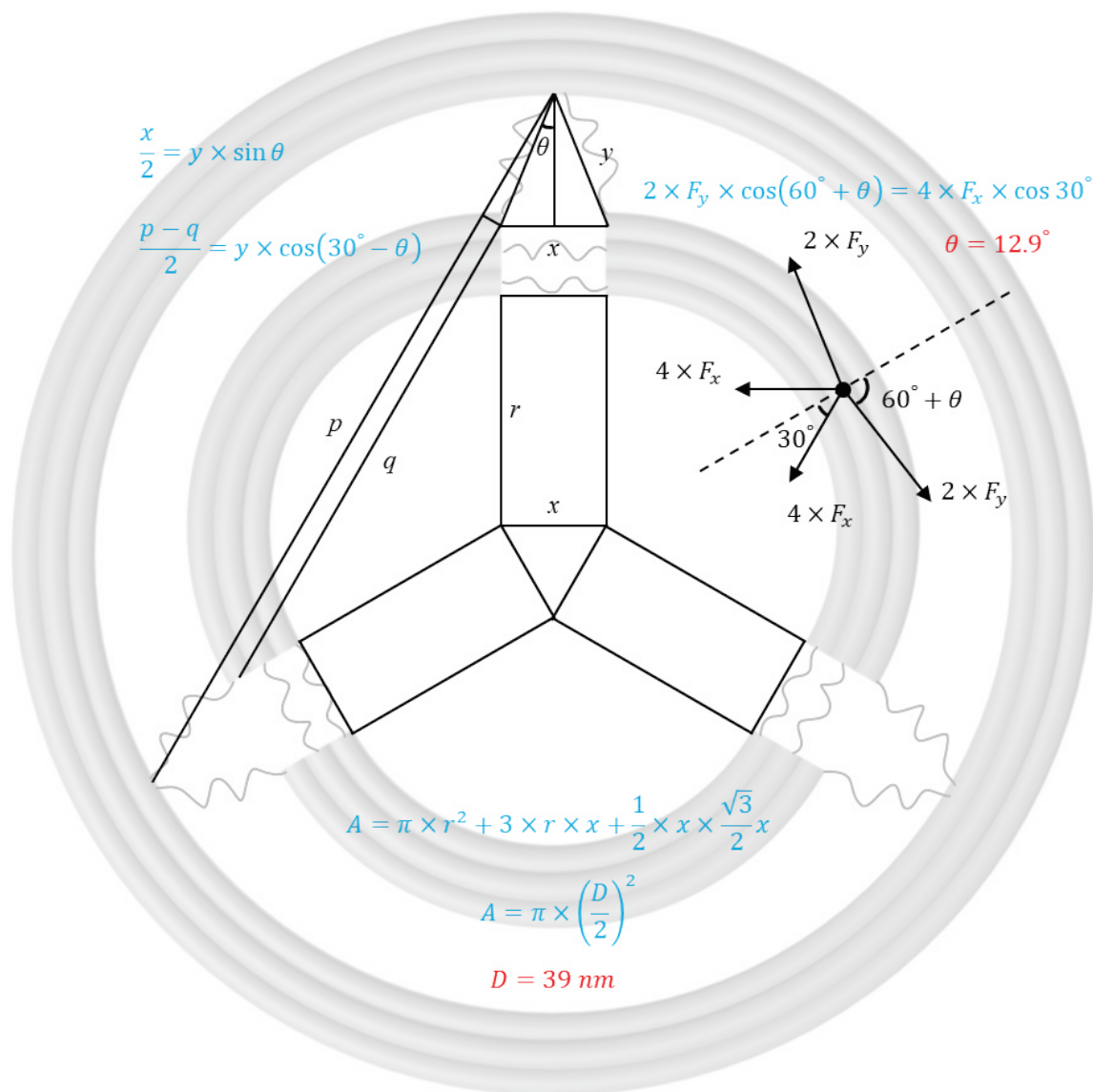

**Figure S9. Evaluation of DNA inner ring dimensions for SRf.** At equilibrium, the forces acting on any inner arc are balanced along the dashed-line direction, resulting in a $\theta$ value of $12.9^\circ$. Thus, the area of the inner ring ($A$) is calculated to derive the inner diameter ($D$) of $\sim 39$ nm, using $A = \pi(D/2)^2$.

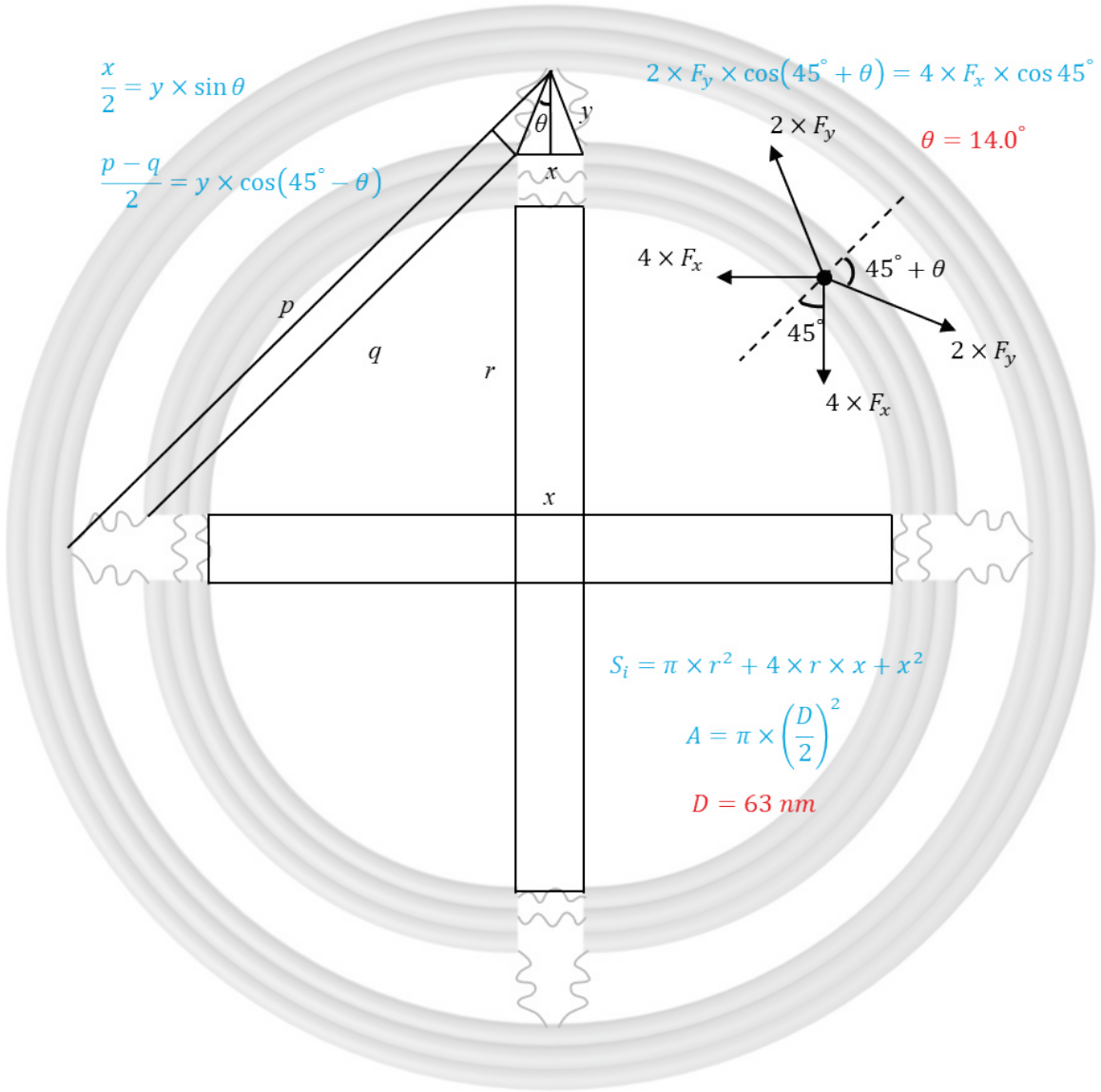

**Figure S10. Evaluation of DNA inner ring dimensions for LRF.** At equilibrium, the forces acting on any inner arc are balanced along the dashed-line direction, resulting in a $\theta$ value of $14.0^\circ$. Thus, the area of the inner ring ($A$) is calculated to derive the inner diameter ($D$) of $\sim 63 \text{ nm}$.

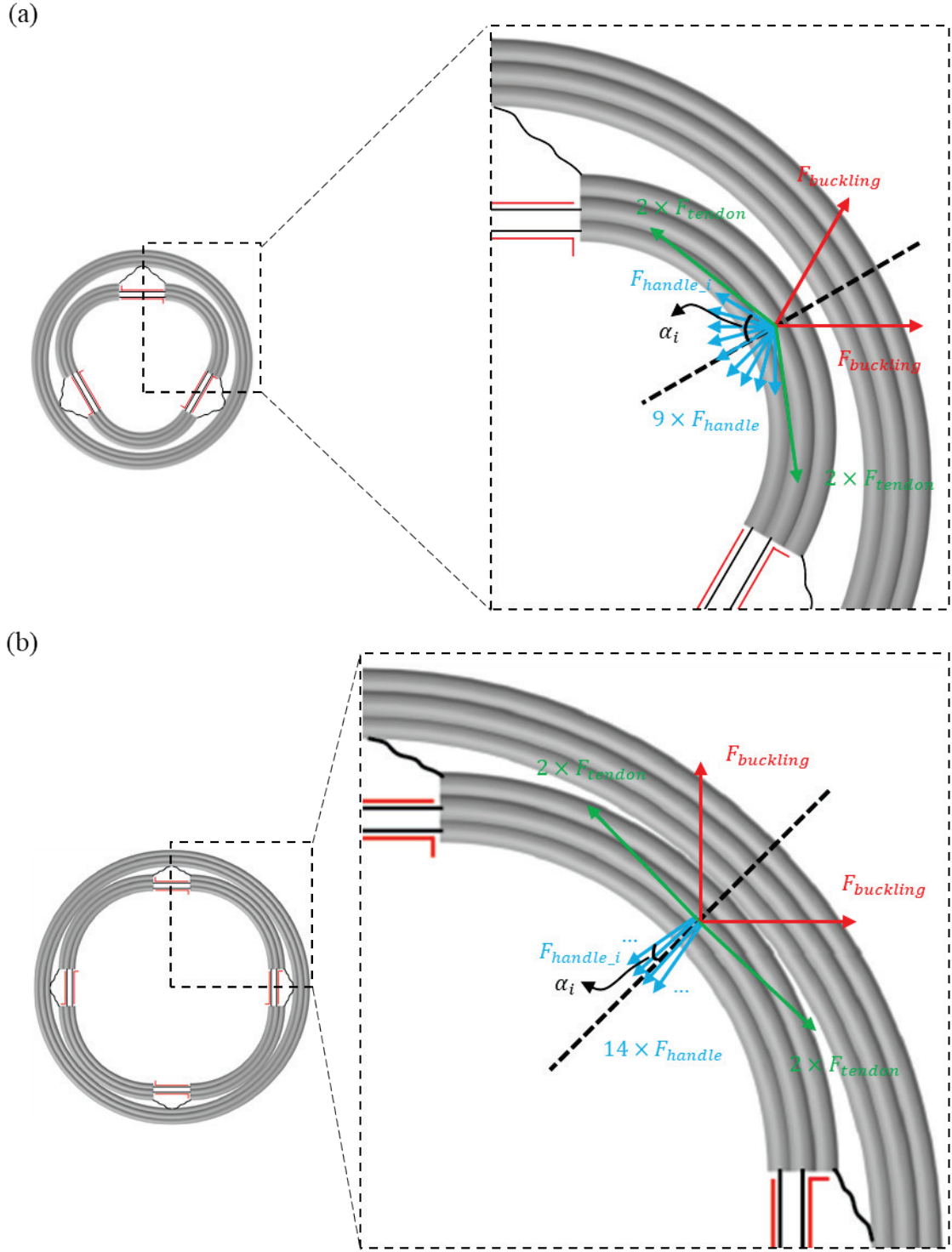

**Figure S11. Pulling forces on DNA handles for (a) SRO and (b) LRO.** The inward pulling forces on handles ($F_{handle}$), the compressive force on DNA struts ($F_{buckling}$), and the tensile forces on DNA tendons ($F_{tendon}$) all act on an inner arc and are balanced along the dashed-line direction at equilibrium.

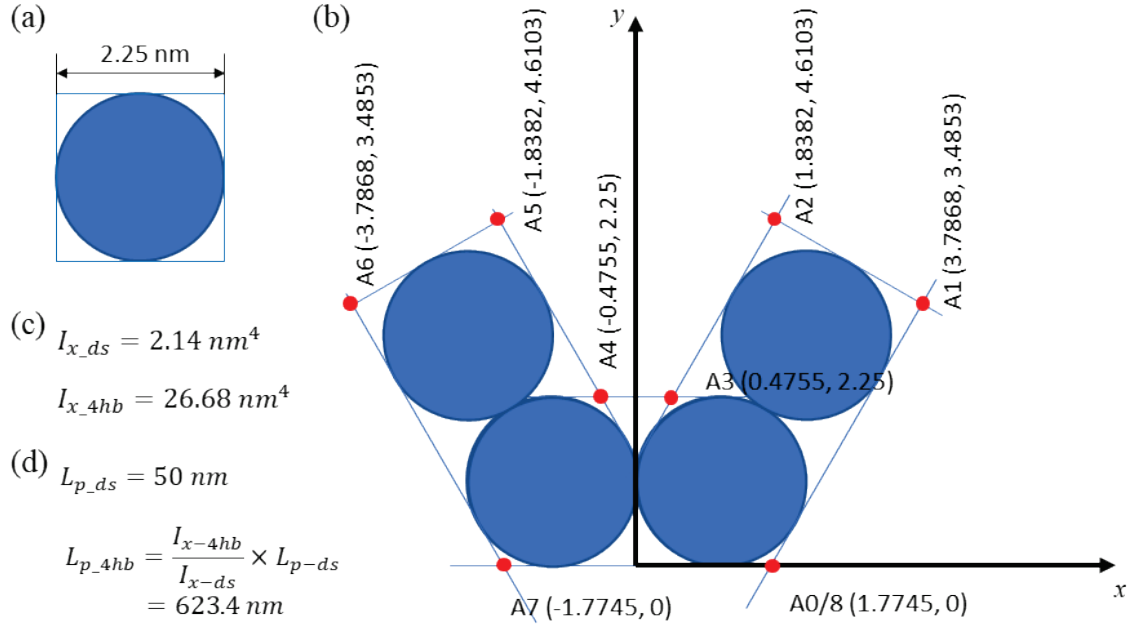

**Figure S12. Persistence length of the four-helix bundle (4hb) struts in LRo.** Polygon models of (a) a double-stranded DNA duplex (ds) and (b) the 4hb strut connecting adjacent inner arcs in LRo. (c) Calculation of the area moment of inertia for (a) ds and (b) 4hb. (d) Estimated persistence length of the 4hb based on that of ds.

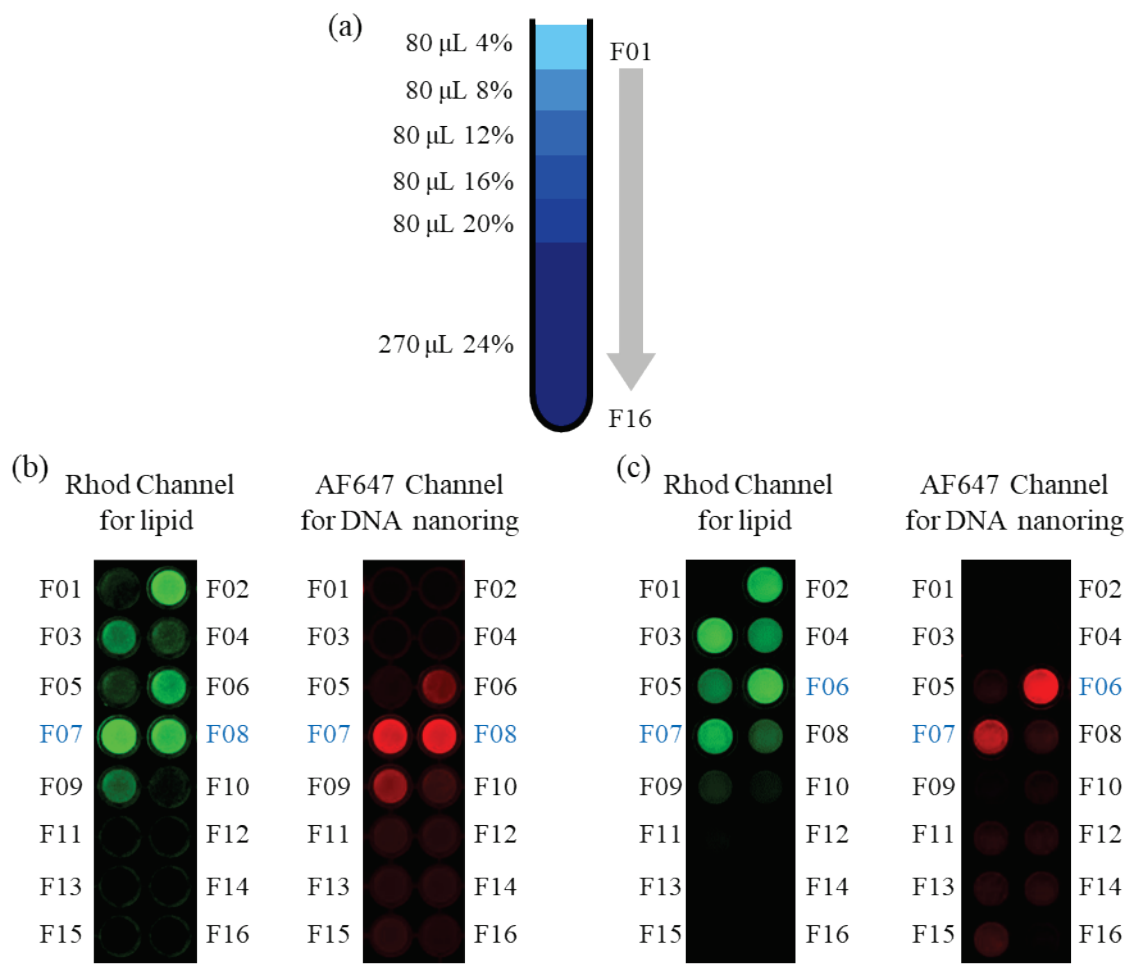

**Figure S13. Purification of DNA-nanoring templated SUVs by isopycnic centrifugation in iodixanol gradients.** (a) Gradient layout. Fractions were sequentially numbered from F01 (top) to F16 (bottom) of the gradient. Fractions with both Rhodamine B (Rhod, green) and Alexa Fluor 647 (AF647, red) fluorescence signals enriched were identified as containing DNA nanoring-templated liposomes: (b) F07–F08 for SRc, (c) F06–F07 for LRc.

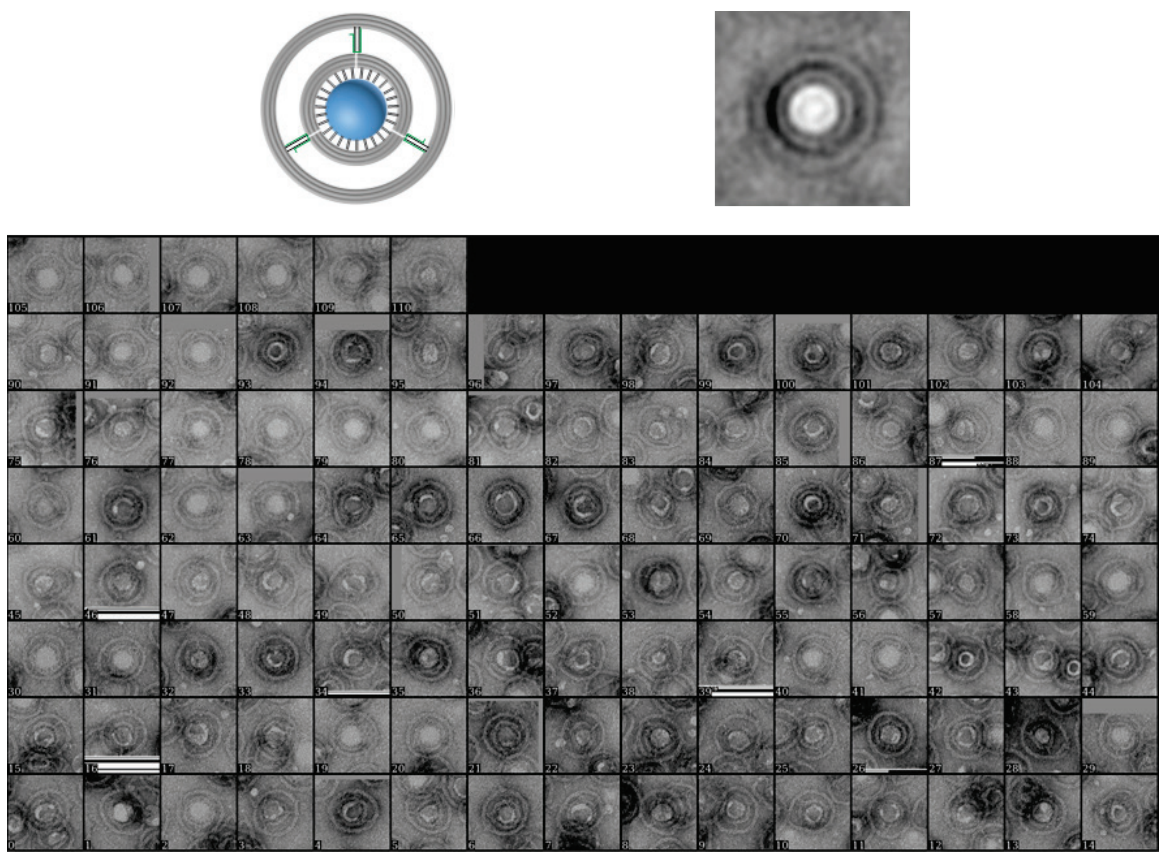

**Figure S14.** Class averaging of 111 particles for SRc-templated liposome. Box size: 125 nm.

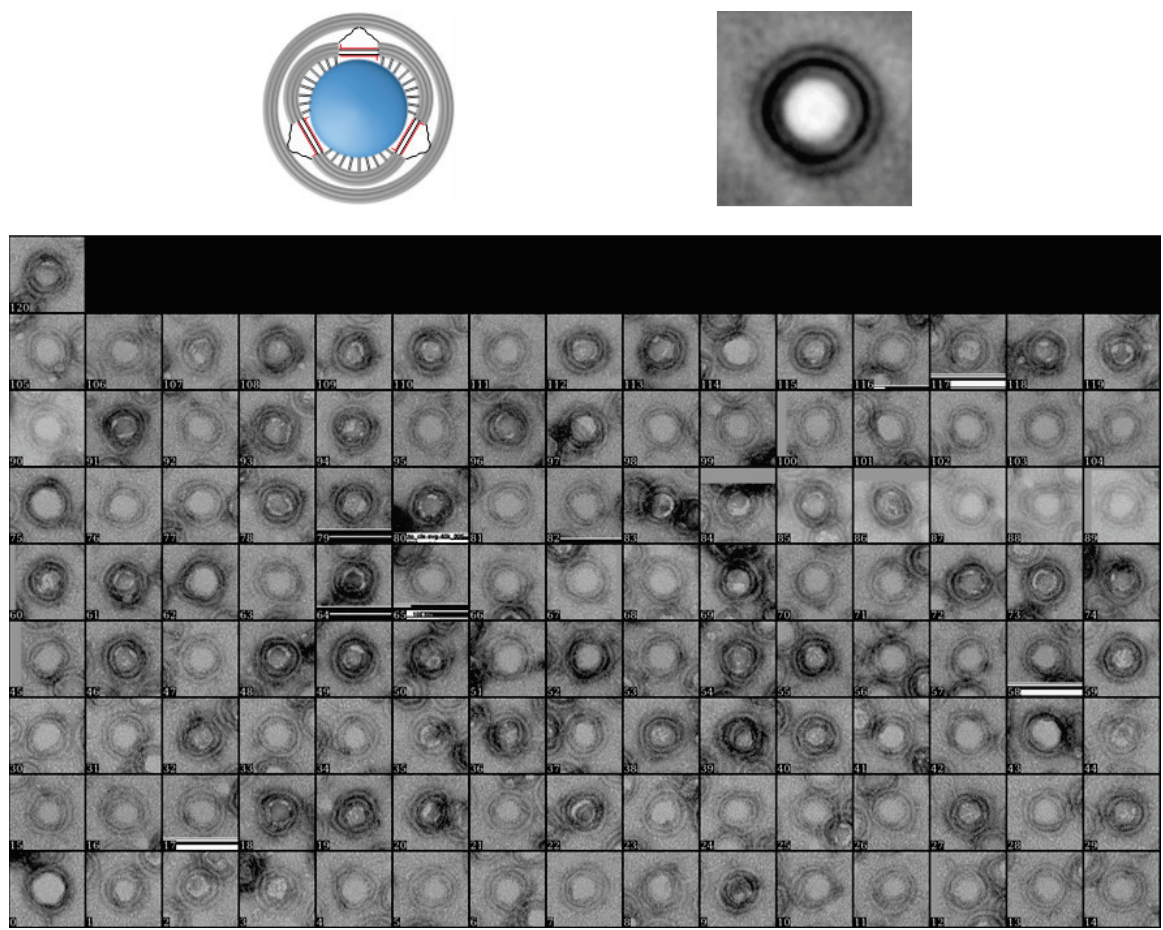

**Figure S15.** Class averaging of 121 particles for liposomes expanded by the small DNA nanoring. Box size: 125 nm.

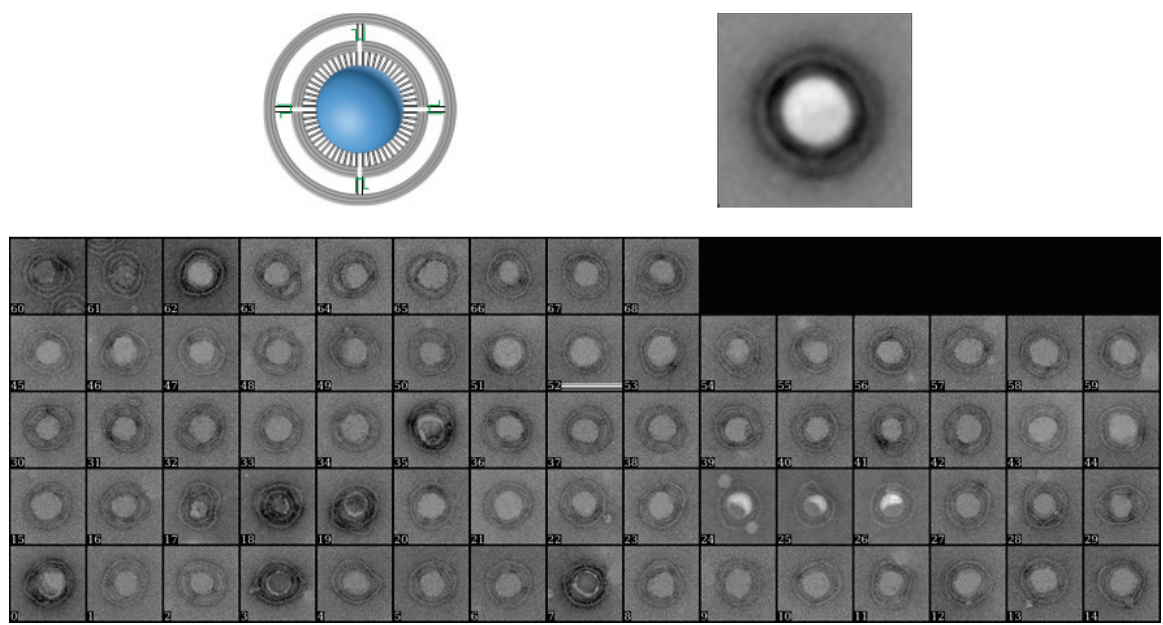

**Figure S16.** Class averaging of 69 particles for LRC-templated liposome. Box size: 156 nm.

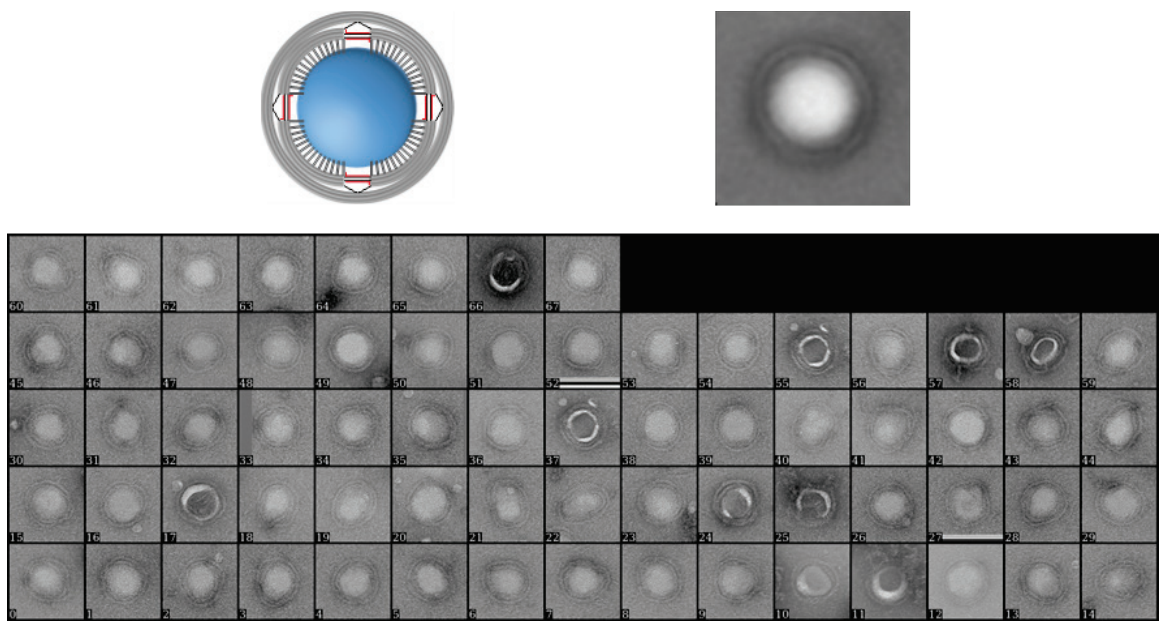

**Figure S17.** Class averaging of 68 particles for liposomes expanded by the large DNA nanoring. Box size: 156 nm.

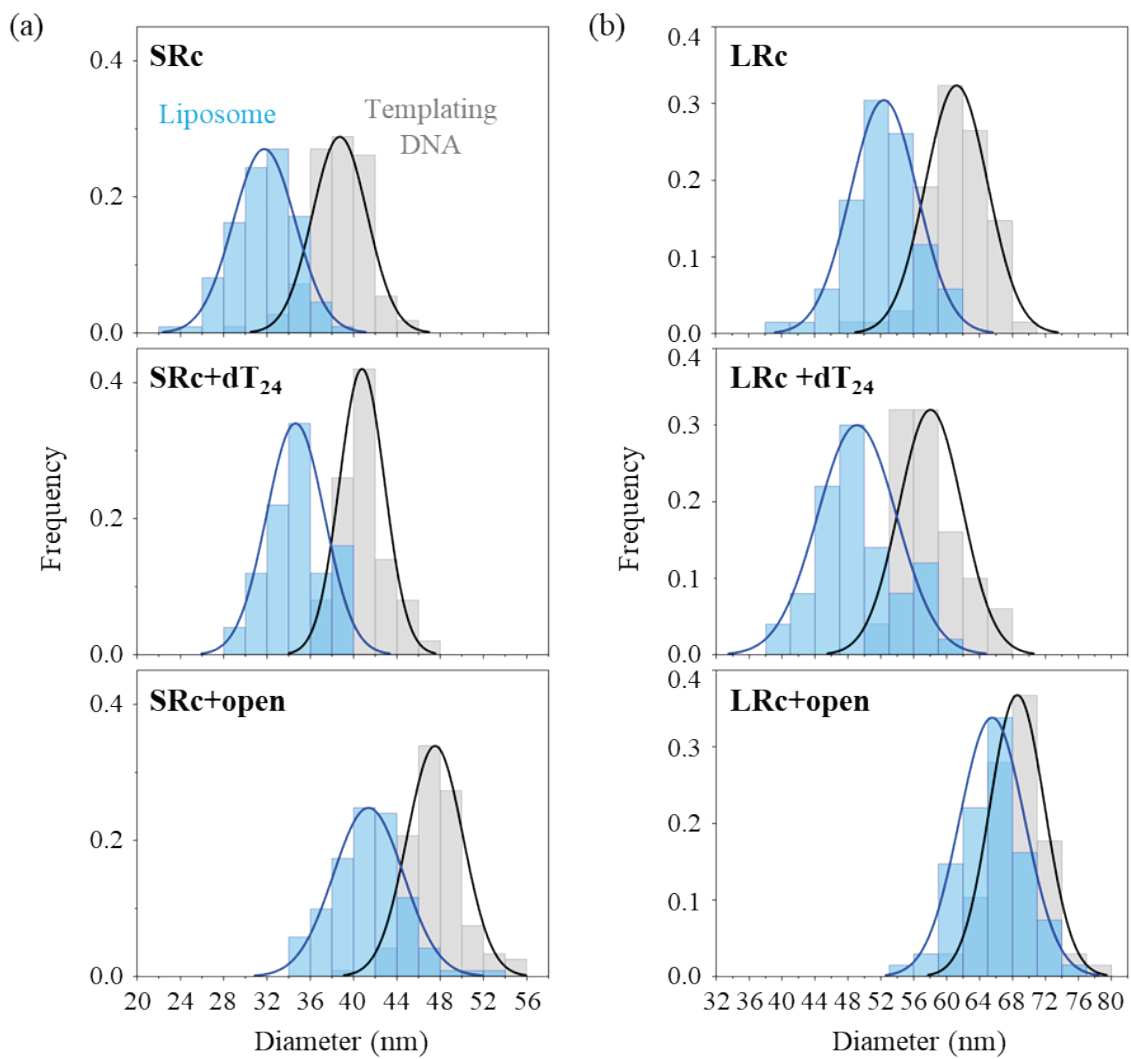

**Figure S18. Size distributions of nanoring-templated liposomes before and after ring dilation.** (a) shows liposomes in the small ring and (b) in the large ring. Blue and grey bars represent measured diameters for liposomes and DNA inner rings, respectively. Top panels: Liposomes formed in the closed DNA rings; middle panels: liposomes in closed rings after incubation with an irrelevant strand (dT<sub>24</sub>); bottom panels: liposomes in closed rings after incubation with ring-opening DNA strands.

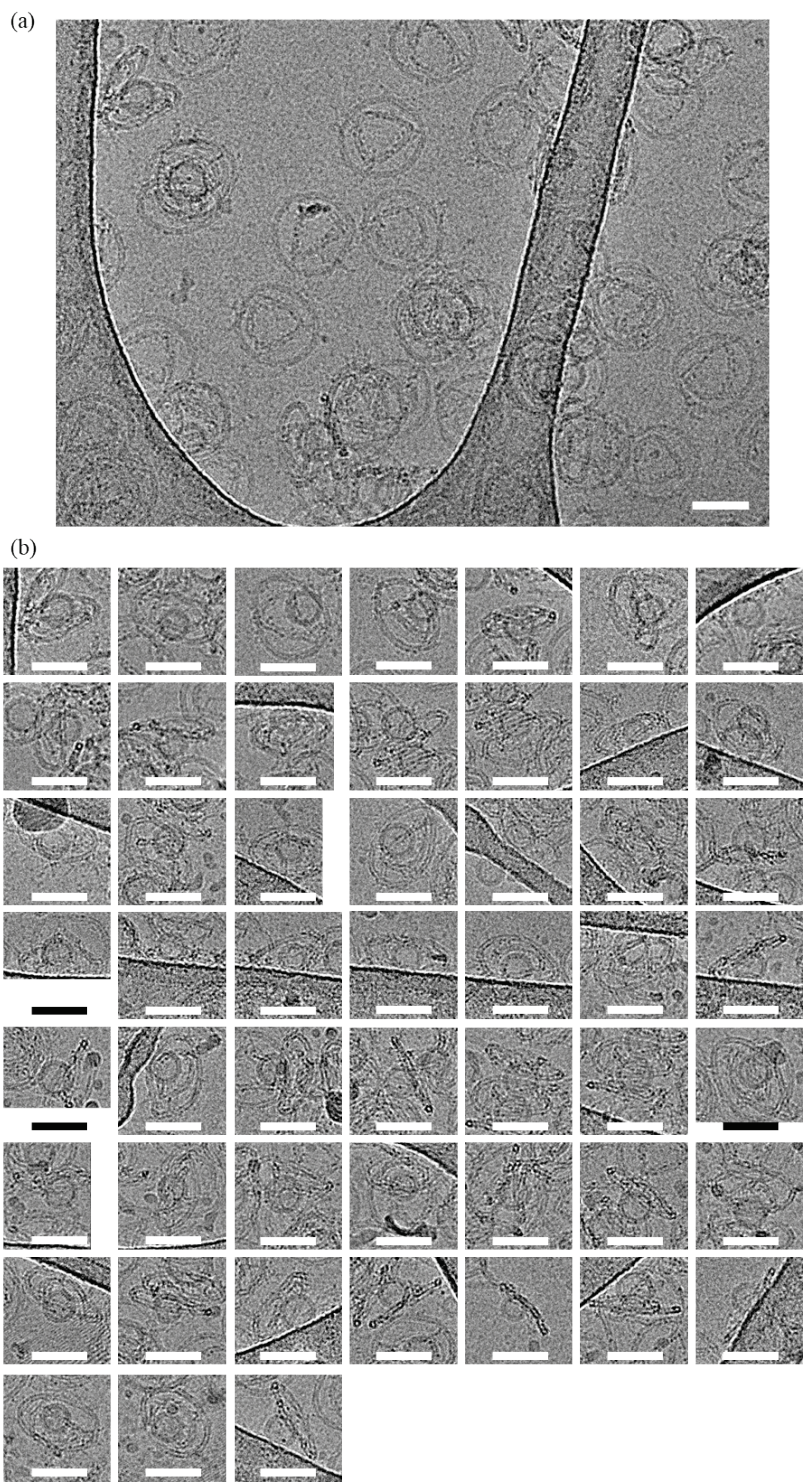

**Figure S19. Representative cryo-EM images of SRc-templated liposomes. (a)** A representative zoom-out image. A notably large number of vesicles ruptured during cryogenic processing. **(b)** selected zoom-ins including side views. Scale bars: 50 nm.

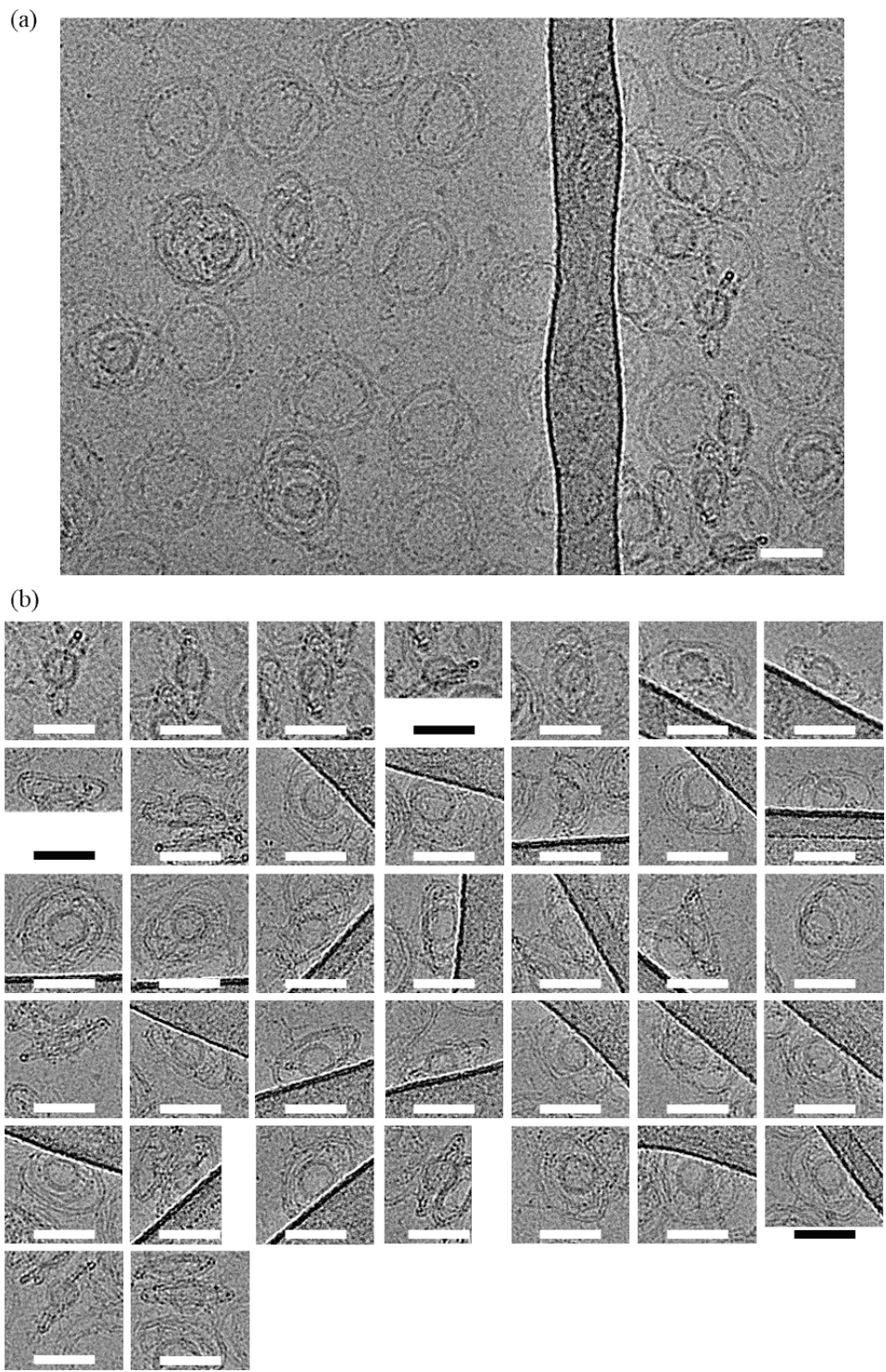

**Figure S20. Representative cryo-EM images of SRc-templated liposomes after ring dilation. (a)** A representative zoom-out image. A notably large number of vesicles ruptured during cryogenic processing. **(b)** selected zoom-ins including side views. Scale bars: 50 nm.

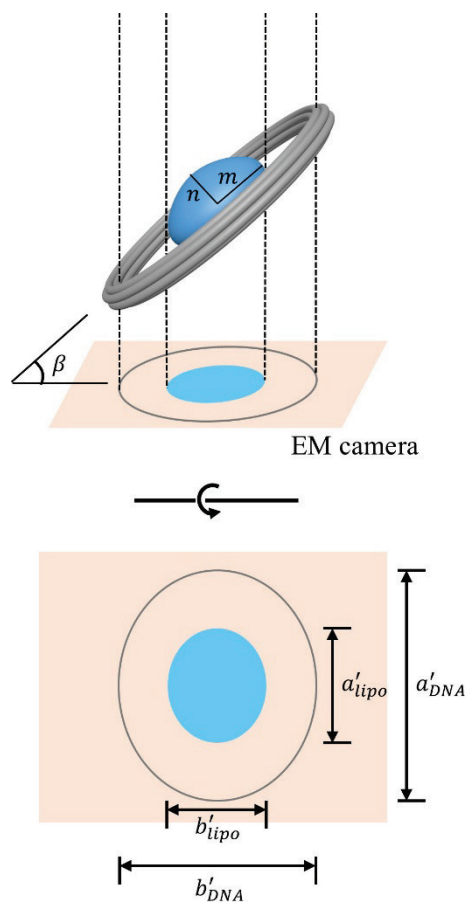

**Figure S21. Calibration of the liposome aspect ratio (AR) using the DNA outer ring as a reference.**
See “Aspect ratio (AR) of liposomes measured by Cryo-EM” in the Notes section for details.

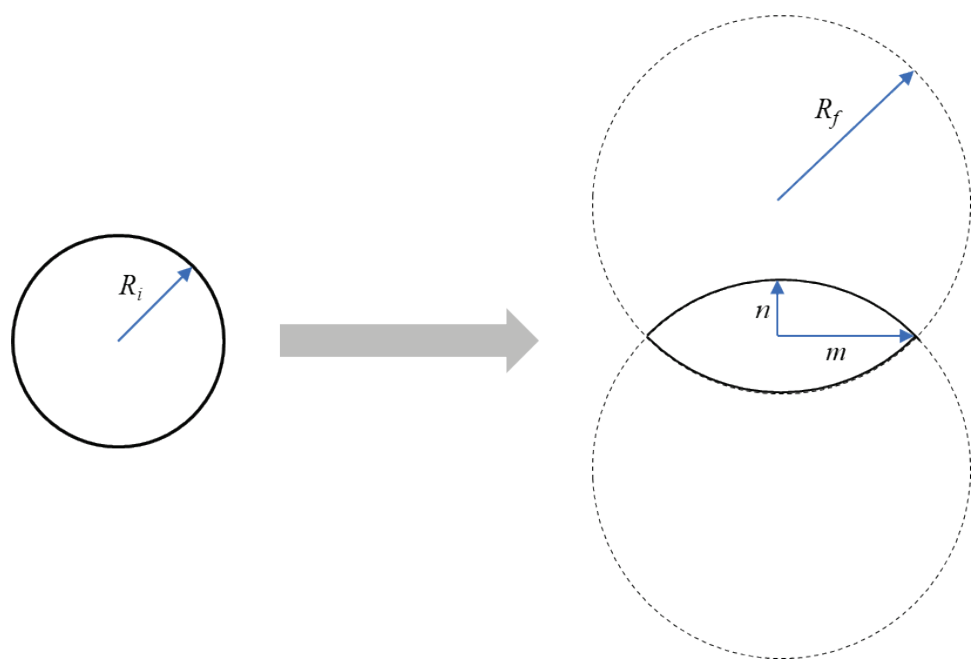

**Figure S22. Liposome models before (sphere) and after (spherical lens) expansion.** $R_i$ represents the radius of liposomes before expansion. $m$ and $n$ represent the semi-major and semi-minor axes of liposomes after expansion, respectively.

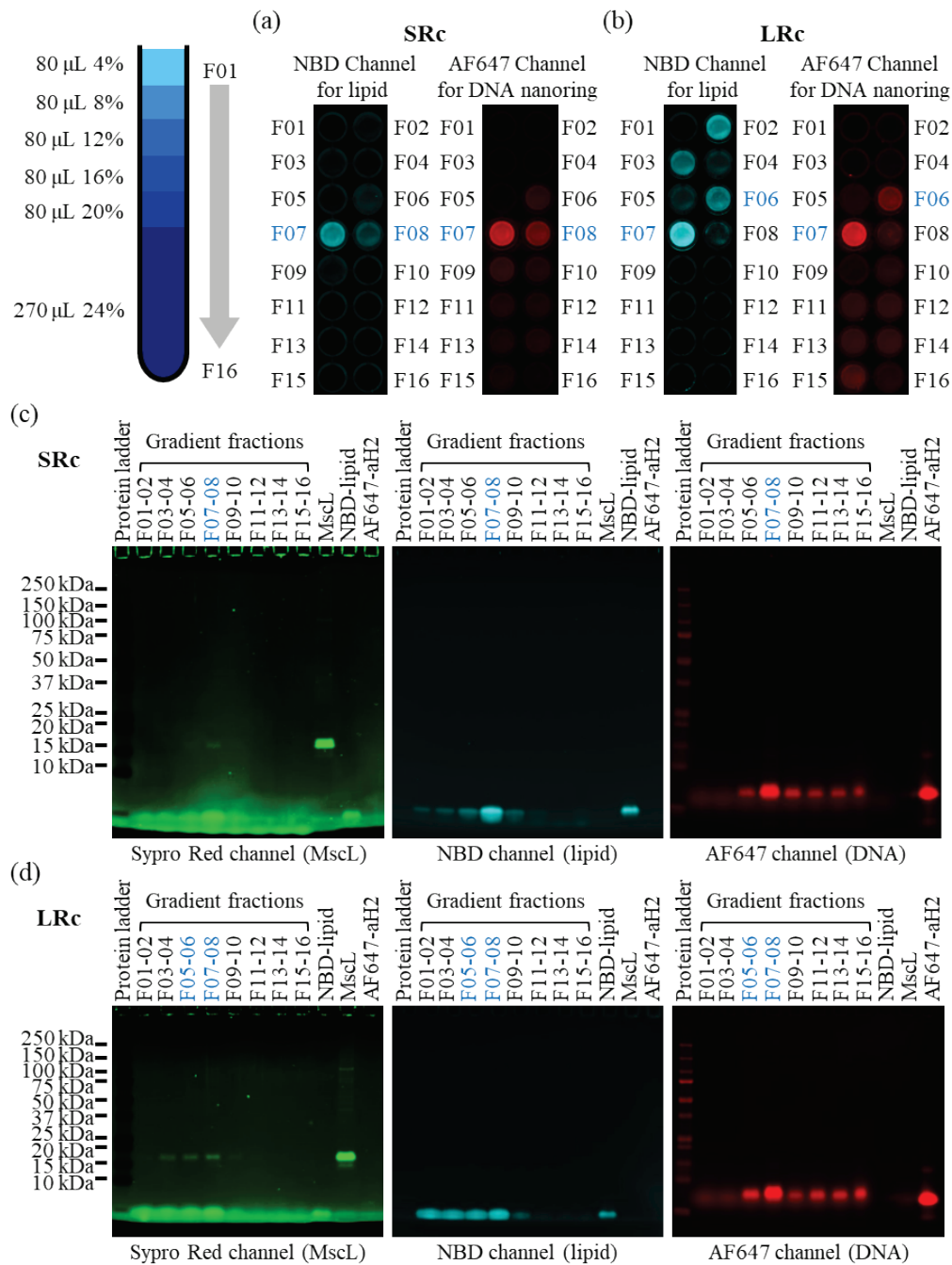

**Figure S23. Incorporating MscL into DNA nanoring-templated liposomes.** (a,b) Purification of DNA-nanoring templated, MscL-embedded SUVs by isopycnic centrifugation in iodixanol gradients. Fractions were sequentially numbered from F01 (top) to F16 (bottom) of the gradient. Fractions with both nitrobenzodiazole (NBD) and Alexa Fluor 647 (AF647) fluorescence signals enriched, specifically F07–08 for (a) SRc, and F06–07 for (b) LRc, were identified as containing DNA nanoring-templated liposomes. (c,d) Analysis of gradient fractions by 4–12% SDS-PAGE. Colocalization of AF647, NBD, and Sypro Red signals in these fractions indicated the co-presence of DNA nanorings, liposomes, and MscL protein, confirming the successful incorporation of MscL. aH: anti-handle.

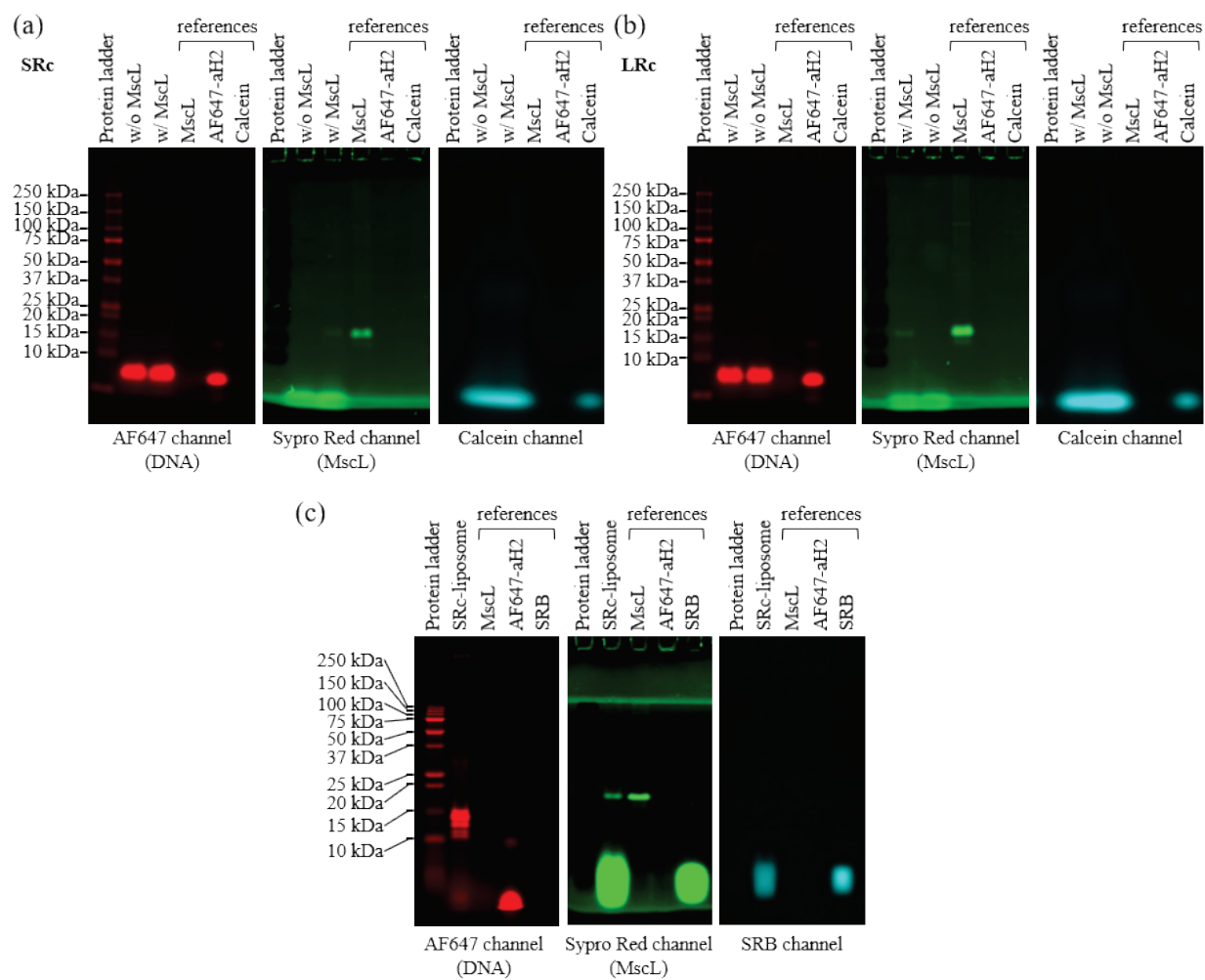

**Figure S24. Gel electrophoresis analysis confirming the formation of MscL-embedded, dye-loaded, DNA nanoring-templated liposomes.** SDS-PAGE was performed on calcein-loaded, (a) SRc- and (b) LRc- templated liposomes in the presence (w/ MscL) and absence (w/o MscL) of MscL using 4–12% gradient gels, and on (c) MscL-embedded, SRB-loaded, SRc-templated liposomes (SRc-liposome) using a 4%/10% stacking gel. aH: anti-handle.

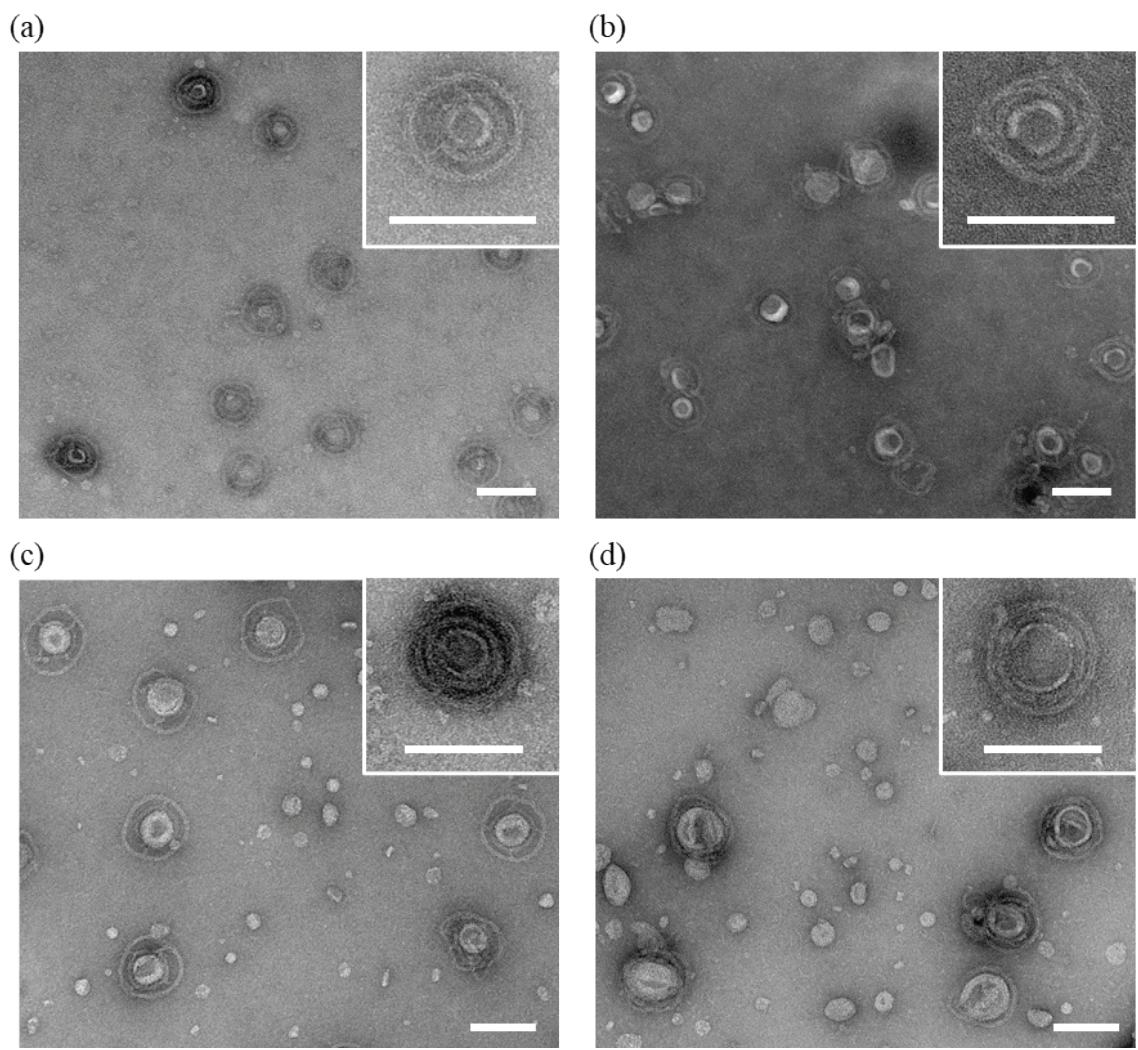

**Figure S25. Negative-stain TEM images of MscL-embedded, DNA nanoring-templated liposomes. (a) SRc + dT<sub>24</sub>, (b) SRc + open, (c) LRc + dT<sub>24</sub>, and (d) LRc + open.**

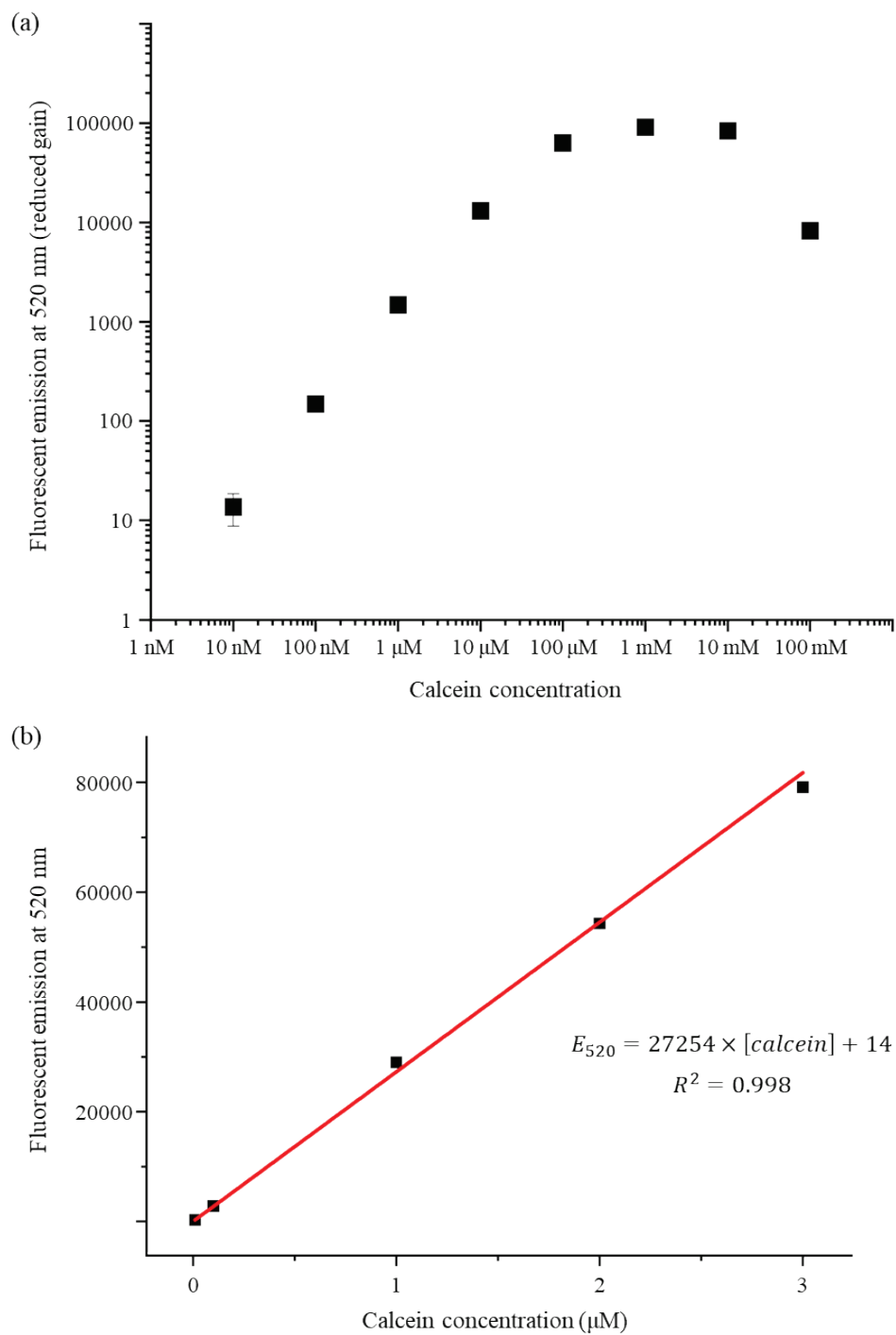

**Figure S26. Calcein fluorescence at 520 nm as a function of concentration.** Below 10 $\mu\text{M}$, calcein fluorescence increases linearly with concentration. Above 1 mM, calcein self-quenching is obvious. We set the liposome-encapsulated calcein concentration to 100 mM.

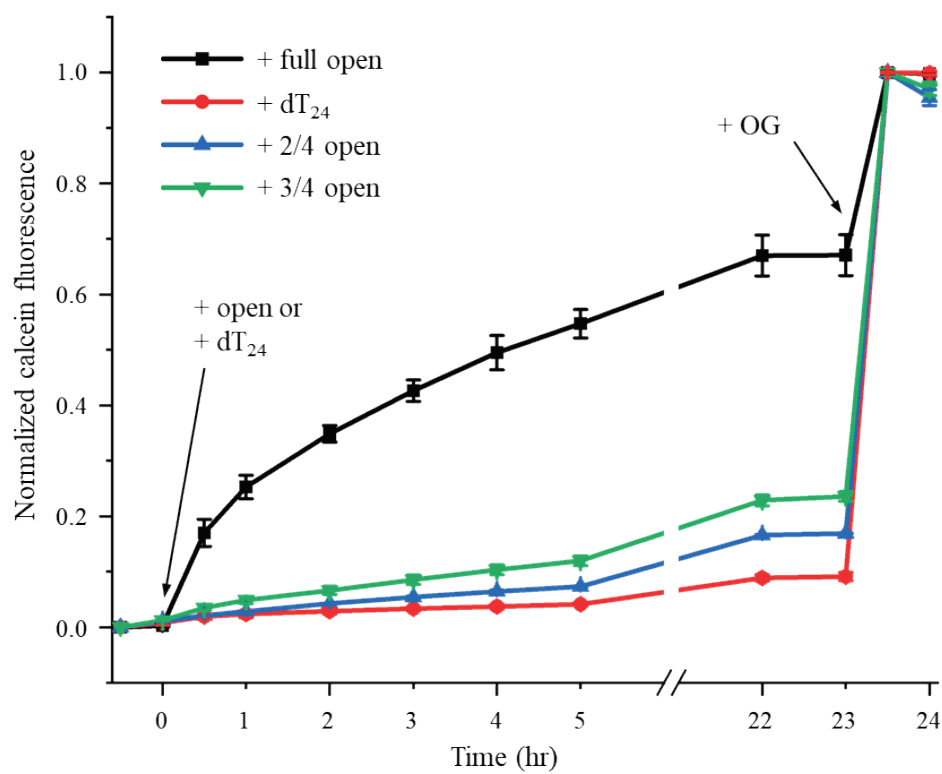

**Figure S27. MscL activation by partially dilated large DNA nanoring.** MscL-embedded liposomes were initially templated by LRc, encapsulating 100 mM calcein. Calcein fluorescence under four conditions were monitored; + full open, + 2/4 open, + 3/4 open: add ring-opening DNA strands that open the inner ring by rigidifying 4, 2 (opposite), or 3 out of the 4 groups of DNA tendons, respectively; + dT<sub>24</sub>: add an irrelevant strand (dT<sub>24</sub>). The black trace is the same as shown in **Figure 3c** (bottom) and shown here for comparison. MscL is known to open within milliseconds at ~12 mN/m of membrane tension<sup>8</sup>, which few SUVs in our setup could reach. The mean membrane tension of expanded SUVs (~6.1 mN/m) is thus consistent with the slower MscL activation kinetics observed. DNA ring dilation might also be slowed down by the SUV.

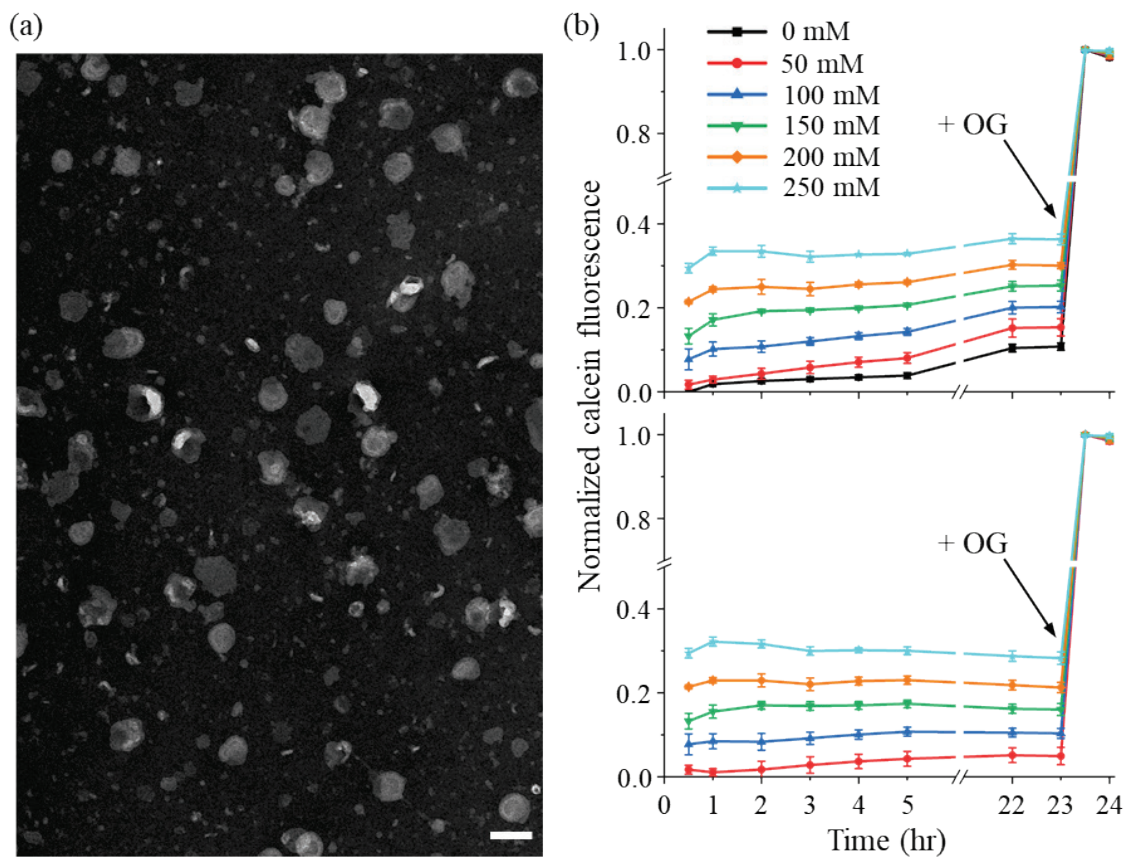

**Figure S28. MscL activation by osmotic pressure.** (a) A representative negative-stain TEM image of template-free MscL-embedded liposomes formed in an iso-osmotic buffer containing 290 mM KCl. Scale bar: 100 nm. (b) Calcein fluorescence traces following exposure of liposomes to hypotonic conditions caused by reducing [KCl] by 0 mM (black, control), 50 mM (red), 100 mM (blue), 150 mM (green), 200 mM (orange), and 250 mM (cyan) at 0 h, followed by the addition of OG to reach 1% OG (w/v) at 23 h (top). The slow rise of fluorescence in untreated liposomes (black trace) suggest spontaneous fusion that leaks calcein in the absence of DNA templates. Therefore, all other traces were corrected by subtracting the calcein fluorescence readings at the same time points without osmotic perturbation (0 mM KCl reduction, black) and replotted (bottom). Note that the fluorescence traces nearly plateaued at 30 min when the first data point was taken.

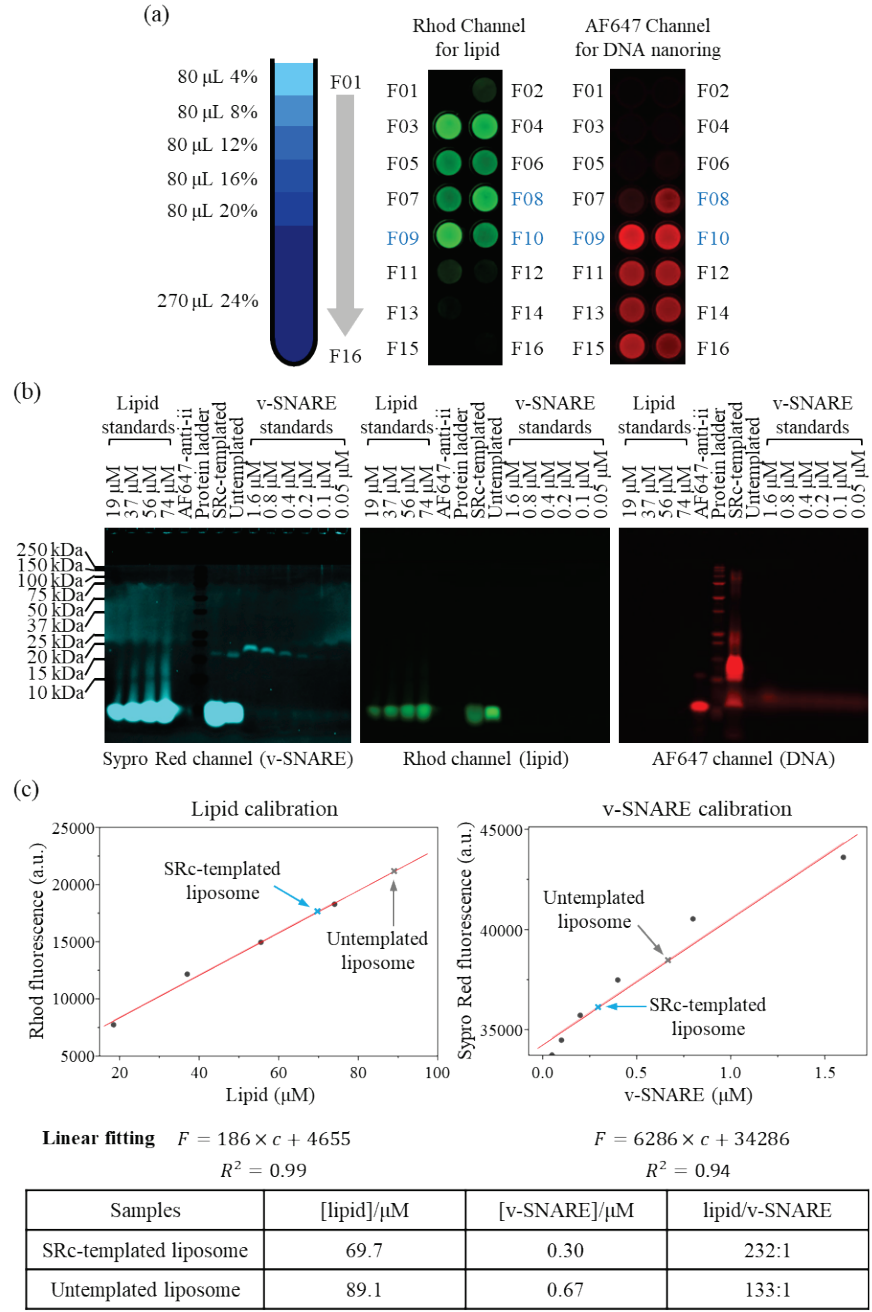

**Figure S29. Incorporation of v-SNARE into SRc-templated liposomes.** (a) Purification of DNA-nanoring templated, VAMP2-embedded SUVs by isopycnic centrifugation in iodixanol gradients. Fractions were sequentially numbered from F01 (top) to F16 (bottom) of the gradient. Fractions with both Rhodamine B (Rhod) and Alexa Fluor 647 (AF647) fluorescence signals enriched, specifically F08–10, were identified as containing DNA nanoring-templated liposomes. (b) SDS-PAGE (4–10%) analysis of both templated and untemplated liposomes. Co-presence of DNA nanorings, liposomes, and VAMP2 protein confirmed the successful incorporation of v-SNARE into DNA nanoring-templated liposomes. (c) Quantitative analysis of lipid and v-SNARE by comparing the band intensities from (b) with corresponding lipid and protein standards. The difference in the lipid:v-SNARE ratio may arise from the steric hindrance imposed by the DNA nanorings and curvature-sensitive VAMP2 incorporation<sup>9</sup>.

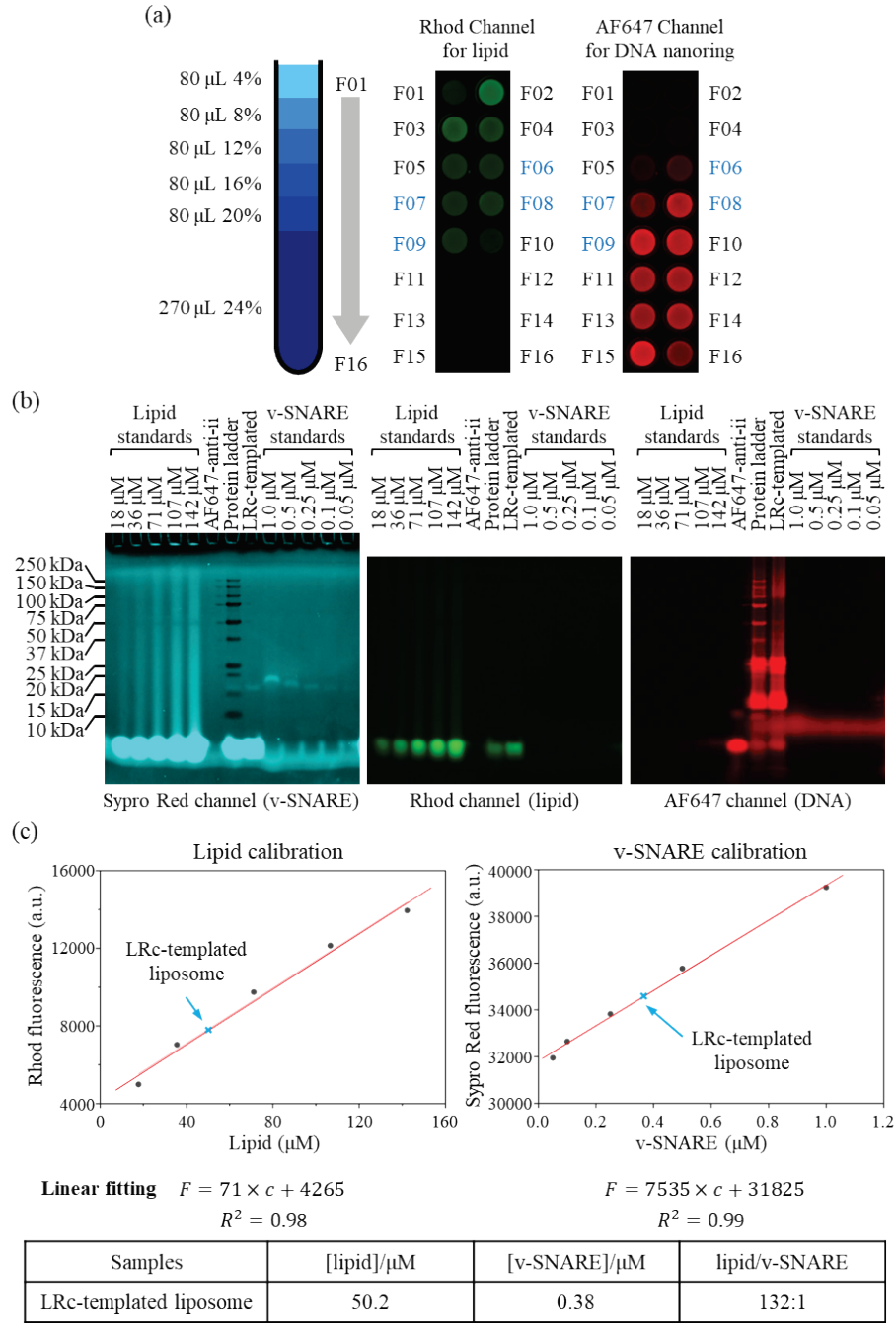

**Figure S30. Incorporation of v-SNARE into LRC-templated liposomes.** (a) Purification of DNA-nanoring templated, VAMP2-embedded SUVs by isopycnic centrifugation in iodixanol gradients. Fractions were sequentially numbered F01 (top)–F16 (bottom) of the gradient. Fractions with both Rhod and AF647 fluorescence signals enriched, specifically F06–09, were identified as containing DNA nanoring-templated liposomes. (b) SDS-PAGE (4–10%) analysis of LRC-templated liposomes. Co-presence of DNA nanorings, liposomes, and VAMP2 protein confirmed the successful incorporation of v-SNARE into DNA nanoring-templated liposomes. Note that the sample in the ‘LRC-templated’ well may have leaked into the ‘Protein ladder’ well, but this does not affect the quantitative analysis. (c) Quantitative analysis of lipid and v-SNARE by comparing the band intensities from (b) with corresponding lipid and protein standards.

**Figure S31. Purification and characterization of t-SNARE liposomes.** (a) Layout of the iodixanol gradient used to purify t-SNARE liposomes by ultracentrifugation. The t-SNARE liposomes were collected from the 0%/20% interface, as indicated by the arrow. (b) SDS-PAGE (4–12%) analysis of the purified t-SNARE liposomes. The gel was stained with Coomassie Blue. (c) A representative cryo-EM image of t-SNARE liposomes. Scale bar: 50 nm. (d) Diameter histogram of t-SNARE liposomes measured from 200 liposomes.

**Figure S32. Representative TEM images of the liposome-fusion experiments. (a)** v-SNARE liposomes formed in the small ring after incubation with liposomes bearing t-SNAREs for 16 h (**Figure 4b**). **(b)** Same as **(a)**, but for v-SNARE liposomes in the large ring (**Figure 4c**). White and yellow boxes highlight the presence and absence of SNARE-mediated liposome-liposome contact, respectively. Scale bars: 50 nm.

**Figure S33. Content-mixing between templated v- and t-SNARE liposomes.** VAMP2-SUVs were formed within SRc and loaded with the SRB dye; t-SNARE-SUVs were formed within LRc and used to fuse with expanded VAMP2-SUVs with or without expansion. **(a)** Normalized SRB fluorescence traces averaged over three independent trials. **(b)** Initial rate derived from the linear fit of normalized SRB fluorescence traces. **(c)** Representative TEM images of the samples in **(a)** at 16 hrs. White boxes highlight the SNARE-mediated liposome-liposome contact. Scale bars: 50 nm.

**Figure S34. Linear fit of NBD traces from lipid-mixing assays for small ring-templated VAMP2-SUVs (expanded).** (a-c) Three technical repeats and (d) their corresponding fit parameters.

**Figure S35. Linear fit of NBD traces from lipid-mixing assays for small ring-templated VAMP2-SUVs (relaxed).** (a-c) Three technical repeats and (d) their corresponding fit parameters.

**Figure S36. Linear fit of NBD traces from lipid-mixing assays for large ring-templated VAMP2-SUVs (expanded).** (a-c) Three technical repeats and (d) their corresponding fit parameters.

**Figure S37. Linear fit of NBD traces from lipid-mixing assays for large ring-templated VAMP2-SUVs (relaxed).** (a-c) Three technical repeats and (d) their corresponding fit parameters.

**Figure S38. Linear fit of SRB traces from content-mixing assays for small ring-templated VAMP2-SUVs (expanded).** (a-c) Three technical repeats and (d) their corresponding fit parameters.

**Figure S39. Linear fit of SRB traces from content-mixing assays for small ring-templated VAMP2-SUVs (relaxed).** (a-c) Three technical repeats and (d) their corresponding fitted parameters.

**Figure S40. Linear fit of SRB traces from content-mixing assays for large ring-templated VAMP2-SUVs (expanded).** (a-c) Three technical repeats and (d) their corresponding fit parameters.

**Figure S41. Linear fit of SRB traces from content-mixing assays for large ring-templated VAMP2-SUVs (relaxed).** (a-c) Three technical repeats and (d) their corresponding fit parameters.

**Figure S42. Linear fit of SRB traces from content-mixing assays for expanded VAMP2-SUVs in the small ring and expanded t-SNARE-SUVs in the large ring. (a-c) Three technical repeats and (d) their corresponding fit parameters.**

**Figure S43. Linear fit of SRB traces from content-mixing assays for expanded VAMP2-SUVs in the small ring and relaxed t-SNARE-SUVs in the large ring. (a-c) Three technical repeats and (d) their corresponding fitted parameters.**

**Figure S44. Purification of MscL by size-exclusion chromatography (SEC).** (a) SEC profile of MscL obtained using a Superdex 200 column on an ÄKTA chromatography system (Cytiva). Four major peaks (1–4) were collected for further analysis. (b) The identity of each peak was assessed by 4–12% SDS-PAGE, followed by Sypro Red staining. Prior to electrophoresis, samples were mixed with Laemmli sample buffer containing 10% (v/v) 2-mercaptoethanol and either incubated at room temperature (RT) or heated to 95 °C for 10 minutes. Based on electrophoretic analysis, the protein content under Peak 1 was identified as MscL.

### SUPPLEMENTARY TABLES

**Table S1. Lipid compositions.**

| Composition | I | II | III | IV | V |
| --- | --- | --- | --- | --- | --- |
| DOPC (mol%) | 82.2 | 82.2 | 83.0 | 80.0 | 85.0 |
| DOPS (mol%) | 15.0 | 15.0 | 15.0 | 15.0 | 15.0 |
| PEG2k-DOPE (mol%) | 2.0 | 2.0 | 2.0 | 2.0 | - |
| Rhodamine-DOPE (mol%) | 0.8 | - | - | 1.5 | - |
| NBD-DOPE (mol%) | - | 0.8 | - | 1.5 | - |
| lipid : SRc | 12675 : 1 |  |  |  | - |
| lipid : LRc | 27562 : 1 |  |  |  | 13230:1 |

DOPC: 1,2-dioleoyl-sn-glycero-3-phosphocholine

DOPS: 1,2-dioleoyl-sn-glycero-3-phospho-L-serine (sodium salt)

PEG2k-DOPE: 1,2-dioleoyl-sn-glycero-3-phosphoethanolamine-N-[methoxy(polyethylene glycol)-2000] (ammonium salt)

Rhodamine-DOPE: 1,2-dioleoyl-sn-glycero-3-phosphoethanolamine-N-(lissamine rhodamine B sulfonyl) (ammonium salt)

NBD-DOPE: 1,2-dioleoyl-sn-glycero-3-phosphoethanolamine-N-(7-nitro-2-1,3-benzoxadiazol-4-yl) (ammonium salt)

**Table S2. DNA sequences.**

| Sequence Name | Sequence |
| --- | --- |
| Scaffold p8064 | TGATAGACGGTTTTTCGCCCTTTGACGTTGGAGTCCACGTTCTTTAATAGTG<br>GACTCTTGTTCCAAACTGGAACAACACTCAACCCTATCTCGGGCTATTCTTT<br>TGATTTATAAGGGATTTTGCCGATTTCGGAACCACCATCAAACAGGATTTTC<br>GCCTGCTGGGGCAAACCAGCGTGGACCGCTTGCTGCAACTCTCTCAGGGCCA<br>GGCGGTGAAGGGCAATCAGCTGTTGCCCGTCTCACTGGTGAAAAGAAAAACC<br>ACCCTGGCGCCCAATACGCAAACCGCCTCTCCCCGCGCGTTGGCCGATTCAT<br>TAATGCAGCTGGCACGACAGGTTTCCCGACTGGAAAGCGGGCAGTGAGCGCA<br>ACGCAATTAATGTGAGTTAGCTCACTCATTAGGCACCCCAGGCTTTACACTT<br>TATGCTTCCGGCTCGTATGTTGTGTGGAATTGTGAGCGGATAACAATTTCAC<br>ACAGGAAACAGCTATGACCATGATTACGAATTCGAGCTCGGTACCCGGGGAT<br>CCTCAACTGTGAGGAGGCTCACGGACGCGAAGAACAGGCACGCGTGCTGGCA<br>GAAACCCCCGGTATGACCGTGAAAACGGCCCGCCGCATTCTGGCCGCAGCAC<br>CACAGAGTGCACAGGCGCGCAGTGACACTGCGCTGGATCGTCTGATGCAGGG<br>GGCACCGGCACCGCTGGCTGCAGGTAACCCGGCATCTGATGCCGTTAACGAT<br>TTGCTGAACACACCAGTGTAAGGGATGTTTATGACGAGCAAAGAAACCTTTA<br>CCCATTACCAGCCGCAGGGCAACAGTGACCCGGCTCATACCGCAACCGCGCC<br>CGGCGGATTGAGTGCGAAAGCGCCTGCAATGACCCCGCTGATGCTGGACACC<br>TCCAGCCGTAAGCTGGTTGCGTGGGATGGCACCACCGACGGTGCTGCCGTTG<br>GCATTCTTGCGGTTGCTGCTGACCAGACCAGCACCACGCTGACGTTCTACAA<br>GTCCGGCACGTTCCGTTATGAGGATGTGCTCTGGCCGGAGGCTGCCAGCGAC<br>GAGACGAAAAAACGGACCGCGTTTGCCGGAACGGCAATCAGCATCGTTTAAC<br>TTTACCCTTCATCACTAAAGGCCGCTGTGCGGCTTTTTTTTACGGGATTTTTT<br>TTATGTCGATGTACACAACCGCCCAACTGCTGGCGGCAAATGAGCAGAAATT<br>TAAGTTTGATCCGCTGTTTCTGCGTCTCTTTTTCCGTGAGAGCTATCCCTTC<br>ACCACGGAGAAAGTCTATCTCTCACAAATTCCGGGACTGGTAAACATGGCGC<br>TGTACGTTTCGCCGATTGTTTCCGGTGAGGTTATCCGTTCCCGTGGCGGCTC<br>CACCTCTGAAAGCTTGGCACTGGCCGTCGTTTTACAACGTCGTGACTGGGAA |
|  | AACCCTGGCGTTACCCAACTTAATCGCCTTGCAGCACATCCCCCTTTCGCCA<br>GCTGGCGTAATAGCGAAGAGGCCCGCACCGATCGCCCTTCCCAACAGTTGCG<br>CAGCCTGAATGGCGAATGGCGCTTTGCCTGGTTTCCGGCACCAGAAGCGGTG<br>CCGGAAAGCTGGCTGGAGTGCGATCTTCCTGAGGCCGATACTGTCGTCGTCC<br>CCTCAAACTGGCAGATGCACGGTTACGATGCGCCCATCTACACCAACGTGAC<br>CTATCCCATTACGGTCAATCCGCCGTTTGTTCCCACGGAGAATCCGACGGGT<br>TGTTACTCGCTCACATTTAATGTTGATGAAAGCTGGCTACAGGAAGGCCAGA<br>CGCGAATTATTTTTGATGGCGTTCCTATTGGTTAAAAAATGAGCTGATTTAA<br>CAAAAATTTAATGCGAATTTTAACAAAATATTAACGTTTACAATTTAAATAT<br>TTGCTTATACAATCTTCCTGTTTTTGGGGCTTTTCTGATTATCAACCGGGGT<br>ACATATGATTGACATGCTAGTTTTACGATTACCGTTCATCGATTCTCTGT<br>TGCTCCAGACTCTCAGGCAATGACCTGATAGCCTTTGTAGATCTCTCAAAAA<br>TAGCTACCCTCTCCGGCATTAATTTATCAGCTAGAACGGTTGAATATCATAT<br>TGATGGTGATTTGACTGTCTCCGGCCTTTCTCACCCTTTTGAATCTTTACCT<br>ACACATTACTCAGGCATTGCATTTAAAATATATGAGGGTTCTAAAAATTTTT<br>ATCCTTGCGTTGAAATAAAGGCTTCTCCCGCAAAAGTATTACAGGGTCATAA<br>TGTTTTTGGTACAACCGATTTAGCTTTATGCTCTGAGGCTTTATTGCTTAAT<br>TTTGCTAATTCTTTGCCTTGCCTGTATGATTTATTGGATGTTAATGCTACTA<br>CTATTAGTAGAATTGATGCCACCTTTTCAGCTCGCGCCCCAAATGAAAATAT<br>AGCTAAACAGGTTATTGACCATTTGCGAAATGTATCTAATGGTCAAACTAAA<br>TCTACTCGTTCGCAGAATTGGGAATCAACTGTTATATGGAATGAAACTTCCA<br>GACACCGTACTTTAGTTGCATATTTAAAACATGTTGAGCTACAGCATTATAT<br>TCAGCAATTAAGCTCTAAGCCATCCGCAAAAATGACCTCTTATCAAAAGGAG<br>CAATTAAAGGTACTCTCTAATCCTGACCTGTTGGAGTTTGCTTCCGGTCTGG<br>TTCGCTTTGAAGCTCGAATTAAAACGCGATATTTGAAGTCTTTCGGGCTTCC<br>TCTTAATCTTTTTGATGCAATCCGCTTTGCTTCTGACTATAATAGTCAGGGT<br>AAAGACCTGATTTTTGATTTATGGTCATTCTCGTTTTCTGAACTGTTTAAAG<br>CATTTGAGGGGGATTCAATGAATATTTATGACGATTCCGCAGTATTGGACGC<br>TATCCAGTCTAAACATTTTACTATTACCCCCTCTGGCAAAACTTCTTTTGCA<br>AAAGCCTCTCGCTATTTTGGTTTTTATCGTCGTCTGGTAAACGAGGGTTATG<br>ATAGTGTTGCTCTTACTATGCCTCGTAATTCCTTTTGGCGTTATGTATCTGC<br>ATTAGTTGAATGTGGTATTCCTAAATCTCAACTGATGAATCTTTCTACCTGT<br>AATAATGTTGTTCCGTTAGTTCGTTTTATTAACGTAGATTTTTCTTCCCAAC<br>GTCCTGACTGGTATAATGAGCCAGTTCTTAAAATCGCATAAGGTAATTCACA<br>ATGATTAAAGTTGAAATTAAACCATCTCAAGCCCAATTTACTACTCGTTCTG<br>GTGTTTCTCGTCAGGGCAAGCCTTATTCACTGAATGAGCAGCTTTGTTACGT<br>TGATTTGGGTAATGAATATCCGGTTCTTGTCAAGATTACTCTTGATGAAGGT<br>CAGCCAGCCTATGCGCCTGGTCTGTACACCGTTCATCTGTCCTCTTTCAAAG<br>TTGGTCAGTTCGGTTCCCTTATGATTGACCGTCTGCGCCTCGTTCCGGCTAA<br>GTAACATGGAGCAGGTCGCGGATTTCGACACAATTTATCAGGCGATGATACA<br>AATCTCCGTTGTACTTTGTTTCGCGCTTGGTATAATCGCTGGGGGTCAAAGA<br>TGAGTGTTTTAGTGTATTCTTTTGCCTCTTTCGTTTTAGGTTGGTGCCTTCG<br>TAGTGGCATTACGTATTTTACCCGTTTAATGGAAACTTCCTCATGAAAAAGT<br>CTTTAGTCCTCAAAGCCTCTGTAGCCGTTGCTACCCTCGTTCCGATGCTGTC<br>TTTCGCTGCTGAGGGTGACGATCCCGCAAAAGCGGCCTTTAACTCCCTGCAA<br>GCCTCAGCGACCGAATATATCGGTTATGCGTGGGCGATGGTTGTTGTCATTG<br>TCGGCGCAACTATCGGTATCAAGCTGTTTAAGAAATTCACCTCGAAAGCAAG<br>CTGATAAACCGATACAATTAAAGGCTCCTTTTGGAGCCTTTTTTTTGGAGAT<br>TTTCAACGTGAAAAAATTATTATTCGCAATTCCTTTAGTTGTTCCTTTCTAT<br>TCTCACTCCGCTGAAACTGTTGAAAGTTGTTTAGCAAAATCCCATACAGAAA<br>ATTCATTTACTAACGTCTGGAAAGACGACAAAACTTTAGATCGTTACGCTAA |
|  | CTATGAGGGCTGTCTGTGGAATGCTACAGGCGTTGTAGTTTGTACTGGTGAC<br>GAAACTCAGTGTTACGGTACATGGGTTCCTATTGGGCTTGCTATCCCTGAAA<br>ATGAGGGTGGTGGCTCTGAGGGTGGCGGTTCTGAGGGTGGCGGTTCTGAGGG<br>TGGCGGTACTAAACCTCCTGAGTACGGTGATACACCTATTCCGGGCTATACT<br>TATATCAACCCTCTCGACGGCACTTATCCGCCTGGTACTGAGCAAAACCCCG<br>CTAATCCTAATCCTTCTCTTGAGGAGTCTCAGCCTCTTAATACTTTCATGTT<br>TCAGAATAATAGGTTCCGAAATAGGCAGGGGGCATTAACTGTTTATACGGGC<br>ACTGTTACTCAAGGCACTGACCCCGTTAAAACTTATTACCAGTACACTCCTG<br>TATCATCAAAAGCCATGTATGACGCTTACTGGAACGGTAAATTCAGAGACTG<br>CGCTTTCCATTCTGGCTTTAATGAGGATTTATTTGTTTGTGAATATCAAGGC<br>CAATCGTCTGACCTGCCTCAACCTCCTGTCAATGCTGGCGGCGGCTCTGGTG<br>GTGGTTCTGGTGGCGGCTCTGAGGGTGGTGGCTCTGAGGGTGGCGGTTCTGA<br>GGGTGGCGGCTCTGAGGGAGGCGGTTCCGGTGGTGGCTCTGGTTCCGGTGAT<br>TTTGATTATGAAAAGATGGCAAACGCTAATAAGGGGGCTATGACCGAAAATG<br>CCGATGAAAACGCGCTACAGTCTGACGCTAAAGGCAAACTTGATTCTGTGCG<br>TACTGATTACGGTGCTGCTATCGATGGTTTCATTGGTGACGTTTCCGGCCTT<br>GCTAATGGTAATGGTGCTACTGGTGATTTTGCTGGCTCTAATTCCCAAATGG<br>CTCAAGTCGGTGACGGTGATAATTCACCTTTAATGAATAATTTCCGTCAATA<br>TTTACCTTCCCTCCCTCAATCGGTTGAATGTCGCCCTTTTGTCTTTGGCGCT<br>GGTAAACCATATGAATTTTCTATTGATTGTGACAAAATAAACTTATTCCGTG<br>GTGTCTTTGCGTTTCTTTTATATGTTGCCACCTTTATGTATGTATTTTCTAC<br>GTTTGCTAACATACTGCGTAATAAGGAGTCTTAATCATGCCAGTTCTTTTGG<br>GTATTCCGTTATTATTGCGTTTCCTCGGTTTCCTTCTGGTAACTTTGTTCGG<br>CTATCTGCTTACTTTTCTTAAAAAGGGCTTCGGTAAGATAGCTATTGCTATT<br>TCATTGTTTCTTGCTCTTATTATTGGGCTTAACTCAATTCTTGTGGGTTATC<br>TCTCTGATATTAGCGCTCAATTACCCTCTGACTTTGTTCAGGGTGTTCAGTT<br>AATTCTCCCGTCTAATGCGCTTCCCTGTTTTTATGTTATTCTCTCTGTAAAG<br>GCTGCTATTTTCATTTTTGACGTTAAACAAAAAATCGTTTCTTATTTGGATT<br>GGGATAAATAATATGGCTGTTTATTTTGTAACTGGCAAATTAGGCTCTGGAA<br>AGACGCTCGTTAGCGTTGGTAAGATTCAGGATAAAATTGTAGCTGGGTGCAA<br>AATAGCAACTAATCTTGATTTAAGGCTTCAAAACCTCCCGCAAGTCGGGAGG<br>TTCGCTAAAACGCCTCGCGTTCTTAGAATACCGGATAAGCCTTCTATATCTG<br>ATTTGCTTGCTATTGGGCGCGGTAATGATTCCTACGATGAAAATAAAAACGG<br>CTTGCTTGTTCTCGATGAGTGCGGTACTTGGTTTAATACCCGTTCTTGGAAT<br>GATAAGGAAAGACAGCCGATTATTGATTGGTTTCTACATGCTCGTAAATTAG<br>GATGGGATATTATTTTTCTTGTTCAGGACTTATCTATTGTTGATAAACAGGC<br>GCGTTCTGCATTAGCTGAACATGTTGTTTATTGTCGTCGTCTGGACAGAATT<br>ACTTTACCTTTTGTCGGTACTTTATATTCTCTTATTACTGGCTCGAAAATGC<br>CTCTGCCTAAATTACATGTTGGCGTTGTTAAATATGGCGATTCTCAATTAAG<br>CCCTACTGTTGAGCGTTGGCTTTATACTGGTAAGAATTTGTATAACGCATAT<br>GATACTAAACAGGCTTTTTCTAGTAATTATGATTCCGGTGTTTATTCTTATT<br>TAACGCCTTATTTATCACACGGTCGGTATTTCAAACCATTAAATTTAGGTCA<br>GAAGATGAAATTAACTAAAATATATTTGAAAAAGTTTTCTCGCGTTCTTTGT<br>CTTGCGATTGGATTTGCATCAGCATTTACATATAGTTATATAACCCAACCTA<br>AGCCGGAGGTTAAAAAGGTAGTCTCTCAGACCTATGATTTTGATAAATTCAC<br>TATTGACTCTTCTCAGCGTCTTAATCTAAGCTATCGCTATGTTTTCAAGGAT<br>TCTAAGGGAAAATTAATTAATAGCGACGATTTACAGAAGCAAGGTTATTCAC<br>TCACATATATTGATTTATGTACTGTTTCCATTAAAAAAGGTAATTCAAATGA<br>AATTGTTAAATGTAATTAATTTTGTTTTCTTGATGTTTGTTTCATCATCTTC<br>TTTTGCTCAGGTAATTGAAATGAATAATTCGCCTCTGCGCGATTTTGTAAC<br>TGGTATTCAAAGCAATCAGGCGAATCCGTTATTGTTTCTCCCGATGTAAAAG |
|  | GTACTGTTACTGTATATTCATCTGACGTTAAACCTGAAAATCTACGCAATTT<br>CTTTATTTCTGTTTTACGTGCAAATAATTTTGATATGGTAGGTTCTAACCCT<br>TCCATTATTCAGAAGTATAATCCAAACAATCAGGATTATATTGATGAATTGC<br>CATCATCTGATAATCAGGAATATGATGATAATTCCGCTCCTTCTGGTGGTTT<br>CTTTGTTCCGCAAAATGATAATGTTACTCAAACCTTTTAAAATTAATAACGTT<br>CGGGCAAAGGATTTAATACGAGTTGTCTGAATTGTTTGTAAGTCTAATACTT<br>CTAAATCCTCAAATGTATTATCTATTGACGGCTCTAATCTATTAGTTGTTAG<br>TGCTCCTAAAGATATTTTAGATAACCTTCCTCAATTCCTTTCAACTGTTGAT<br>TTGCCAACTGACCAGATATTGATTGAGGGTTTGATATTTGAGGTTTCAGCAAG<br>GTGATGCTTTAGATTTTTTCATTTGCTGCTGGCTCTCAGCGTGGCACTGTTGC<br>AGGCGGTGTTAATACTGACCGCCTCACCTCTGTTTTATCTTCTGCTGGTGGT<br>TCGTTCGGTATTTTTAATGGCGATGTTTTAGGGCTATCAGTTCGCGCATTAA<br>AGACTAATAGCCATTCAAAAATATTGTCTGTGCCACGTATTCTTACGCTTTC<br>AGGTCAGAAGGGTTCTATCTCTGTTGGCCAGAATGTCCCTTTTATTACTGGT<br>CGTGTGACTGGTGAATCTGCCAATGTAAATAATCCATTTCAGACGATTGAGC<br>GTCAAAATGTAGGTATTTCCATGAGCGTTTTTCCTGTTGCAATGGCTGGCGG<br>TAATATTGTTCTGGATATTACCAGCAAGGCCGATAGTTTGAGTTCTTCTACT<br>CAGGCAAGTGATGTTATTACTAATCAAAGAAGTATTGCTACAACGGTTAATT<br>TGCGTGATGGACAGACTCTTTTACTCGGTGGCCTCACTGATTATAAAAAACAC<br>TTCTCAGGATTCTGGCGTACCGTTCCCTGTCTAAAATCCCTTTAATCGGCCTC<br>CTGTTTAGCTCCCCTCTGATTCTAACGAGGAAAGCACGTTATACGTGCTCG<br>TCAAAGCAACCATAGTACGCGCCCTGTAGCGGCGCATTAAGCGCGGCGGGTG<br>TGGTGGTTACGCGCAGCGTGACCGCTACACTTGCCAGCGCCCTAGCGCCCGC<br>TCCTTTCGCTTTCTTCCCTTCCTTTCTCGCCACGTTCGCCGGCTTTCCCCGT<br>CAAGCTCTAAATCGGGGGCTCCCTTTAGGGTTCCGATTTAGTGCTTTACGGC<br>ACCTCGACCCCAAAAAACTTGATTTGGGTGATGGTTCACGTAGTGGGCCATC<br>GCCC |
| Small outer core_1 | CAAAAGCGTTGGTTCCGGCACCGCTTGGTTGCGTCTGTGG |
| Small outer core_2 | TTTAGGAGGAACAAGACGACAGTATCGGGGTAAAGCCTGCAT |
| Small outer core_3 | CAAGAATACATCAATAGGAACGCCATAGGCTATCAAAGAA |
| Small outer core_4 | TTTCAATTGAATACAATTCGCATTAAATCGATGAAACTAATA |
| Small outer core_5 | TTCTGACGTGATAATATAAGCAAATATTCCGGTTGATATATTTT |
| Small outer core_6 | ATATTTTTAAGAATCGGATCAAAACAGGAAGATTGATAAGGC |
| Small outer core_7 | GGTTGGGAAAGCCAGTTTACCAGTCCCGGCGGCCTTTCGGGAAAC |
| Small outer core_8 | TTTGGAAGTTATCCGCGTGCCGGACTTGTATTTC |
| Small outer core_9 | TCAAAGGCTCGAATAGCAACCGCAAGAATGGGGATGTATGTT |
| Small outer core_10 | TTCTGGCCGCGTCCGCCCACGCAACCAGCTGATCGGTAACATAT |
| Small outer core_11 | AGAATACCACGGTCTCATTGCAGGCG |
| Small outer core_12 | ATTTTTGGCCAGAATCAATCCGCCGGGCGCCTGGTGCTATACCA |
| Small outer core_13 | CCTAAAACCAGCGCCTGCGGCTGGTAATGCCTCAGGAAAACG |
| Small outer core_14 | GAAGATAGGTTACCTTCCCTTACACTGGTGAACCGTGCATCTGC |
| Small outer core_15 | AGGTCAGATTTCAACGTGAGAAAGGCCGGATGGGAACAACCCTC |
| Small outer core_16 | GAGGTCACAAAAACAACCGTTCTAGCTGACAACATTTGAAAG |
| Small outer core_17 | ATTGCAGGTCTTTACCCGGAAGGTTAGCCAG |
| Small outer core_18 | TAATGCTAATTAAGCTTGAGAGATCTACAACAAAAATGTAACAG |
| Small outer core_19 | CGGTGTCATTAACAAGCAAACAAGAGAATTTTTGTTCGCCTG |
| Small outer core_20 | TTCTGCGGGCGCGAGTCAATCATATGTACCTAAATTGTAAACGT |
| Small outer core_21 | GTTTGACGTTTAGCTAATCAGAAAAGCCCCAAAACTTAAATTTCTGCTCATG<br>AAACAGAAACAC |
| Small outer core_22 | ACCGCCTGCCAACGCATCGACATAAAAAAATCTCCGTTATCATA |
| Small outer core_23 | GTTTGCCTTTCCAGTAGTGATGAAGG |
| Small outer core_24 | TCCTGTTATTGCGTTACGATGCTGATTGCCAAACAATGGTATTC |
| Small outer core_25 | AATCAAAAGTGTAATCGTCGCTGGCAGCCTTTCAGACTTGCG |
| Small outer core_26 | ATGGTCACGTGGACAACGGAATACCCAAAAGAACCCAGCTACAATTTTAGTT<br>GGTCACGA |
| Small outer core_27 | CCTCACATAATAAAAACAAAGTTACCAGTAGAACCAGCTG |
| Small outer core_28 | GGGTTTCTGACCTGATTACCGAAGCCCTAACGCAACTGTTGG |
| Small outer core_29 | TGCTGCGAATGGCTGAGGCATAGTGCAAGAAACAAGACAAT |
| Small outer core_30 | CAGACGATCATCGCCATTCAACTAATGCCGTTAATAAGATCG |
| Small outer core_31 | GATGCCGAAACAGAACAGTGCCAAGATTCATCAGACCAGCA |
| Small outer core_32 | TAAAAATTAAGCAAAAAATATCGCGTTTATCACCTGGGATAG |
| Small outer core_33 | CCTGTAAGCTCCTTGATTGCATCAAAAATCTGGCGTCGGA |
| Small outer core_34 | TTAGCAAGTAGCTCTTAATTACATAATCAAAAATCTGAATA |
| Small outer core_35 | GTAGTAGCTGGAAGTGCAAAAGAAGATGACGGATTAAATCAG |
| Small outer core_36 | CATTTGGAACGAGTTTAATGGTTCAGAGGCGAATTTCCCAA |
| Small outer core_37 | GAGGCGGGCTGATTAAGAACGCGAGAAATAATTGAAAAAG |
| Small outer core_38 | CTGTCGAGCGGTCAACTATATGTAAATATACAAATGAGAGA |
| Small outer core_39 | CACATTATGATGGTCAGCCATATTTTTAACCTCCGCGAAAA |
| Small outer core_40 | AAGCATAAAGAATAGGCGTCTTTCCAGACTCCCGAGGTGGAG |
| Small outer core_41 | GCTGGTCTAAGTTGTAAGACTCCTTATAATAATTCCAACG |
| Small outer core_42 | GTCGGTGTATTACGAATACATACATAAAATAGCCGAGGGACA |
| Small outer core_43 | CCAGCATCTGCGCAAAGACACCACGGAAAGCTATCAAGCGTA |
| Small outer core_44 | CTTAGCGCCAATTTTGTCACCTGGCTCATCGGAAAC |
| Small outer core_45 | CGGGTCAGCCAGCTGAAGAAAAATCTAAGATACTGATAGC |
| Small outer core_46 | TGCTCGTCAGGGGACCATTATTACAGGTAGAACGCTGAGAGCCAGCACCTGC<br>AGGTGAGG |
| Small outer core_47 | TAAAGATTCCGTAATTGCTGAACCTCAAAGACTTCCTCCAAC |
| Small outer core_48 | ATCAATAAACAACCTCAGTTGGCAAATAAAGCGTTGATAA |
| Small outer core_49 | AGAGGGTACTTCCTGTATCTAAAATAAATATACAAATTCGC |
| Small outer core_50 | TGCCTGATTTAACCGGGAGAAACAATAATGAAATAAAGTA |
| Small outer core_51 | CGTAAAACTTGTTAACAAGTTACAAAATCGCGTGAAATACCGACCGTCTAAA<br>TAGATTTA |
| Small outer core_52 | GGGCGGTTCTCACGACTAGAAAAAGCCTCAAGACAGCCCTTC |
| Small outer core_53 | CGCACAGGAATTTGTTCTTACCAGTATTTATATCACGCTG |
| Small outer core_54 | GTAGCGCCATACGCTCAACAGGCTTATCCCGGCGAA |
| Small outer core_55 | GTCCGTTGGGAACGCGTTTTAGCGAACGCCTAACCTTATA |
| Small outer core_56 | ATCCTCATAAACGACCTTAAATCAAGATTATCCTGGTTCCAG |
| Small outer core_57 | GAAGGGCTACGGCTGCGCGTGCCTGTTCTTCAACAGAGAAAAGT |
| Small outer core_58 | CAGGCAATCGCACTGCGGCGGGCCGTTTTGTGGCACATGAAA |
| Small outer core_59 | CCACTATTTTGCACTGGCATGATGGTAACG |
| Small outer core_60 | CCAGGGTGAACGTCATTCCTGTGTGAAATTCAAGAGT |
| Small outer core_61 | GCGAAAGGCCAACGTCCCCGGGTACCGAGGCGAAAAACCGAG |
| Small outer core_62 | CACTCCACTGTTGCCAGTGTCACTGCGCGCCGCGAACATAACGC |
| Small outer core_63 | CAGTTTGATAAACAGCAGCCAGCGGTGCCACGAACCTTGAGA |
| Small outer core_64 | GCATCGTTGTTCAGCAAATCTGCAATGCCTGGTGTAAAAAATC |
| Small outer core_65 | GTCACGTTGAGTAATCCCTCATATATTTTAAAGTTAACGGCATCA |
| Small outer core_66 | TTCTCCGGACAGTCCGGGAGAAGCCTTTGATTAGAGAGGAAG |
| Small outer core_67 | CTTTCATTAAATTACTAAATCGGTTGTACTTTTTGCTATAGT |
| Small outer core_68 | GTCTGGCGCTATTTAATAAAGCCTCAGAGTTA |
| Small outer core_69 | CTCATTTGAGTCTGGTCCAATAAATCATACATGCAACCAAACAT |
| Small outer core_70 | TAATATTTAGCATGCTGAAAAGGTGGCATCAGTTGATATTCA |
| Small outer core_71 | AGACGCATTGCCGCTTGGGCGCCAGGGTGGTTTTCAAATGGTCAATAACCTC<br>ATTAGATTTCATC |
| Small outer core_72 | TAGACTTTCCCGTAATGCATTAATGAATCGGGCCCTGGCAAATC |
| Small outer core_73 | ACGTACAAAGTTAAGCGCTCACTGCCCGCCCAGCAGGGCTTA |
| Small outer core_74 | CCGCCACTTTTCGTCAGCCTGGGGTGCCTACAAAATCTTTGCCA |
| Small outer core_75 | CGTTGTAAACGGAACTCACAATTCCACACGAGTGTTAATCTT |
| Small outer core_76 | GTCAGGAGAATTACATTAGTCTTTAATGCTGTGCACGTATGAGC |
| Small outer core_77 | AACTAACATACCACATTAAAAATACCGAGGTGCCCGTTTCTT |
| Small outer core_78 | TAAAGCTAATTCGAGCTTTAACACCGGCAAATGGATGGGC |
| Small outer core_79 | GAATTGATGACTATGGATGGCTTAGAGCCATAAAGATGCCGG |
| Small outer core_80 | TACCTTTAACAAAAAACATGTTTTAAATAGGCAAGGCAGGTCAT |
| Small outer core_81 | ATTGCTTTACCTGATTCATTCCATATAACAATTCTCGGTAAT |
| Small outer core_82 | GTTAAAAGTTAATACATTTCGTCTTTTCACCAGTCAAAT |
| Small outer core_83 | TGCGTTGCTGATAGAGAGTTGCAGCATGCCAGCAAAAAGC |
| Small outer H2_1 | GAAACGCTACGCAGTGCTGCAAGGCGATTGGTCAGCTCGTAATC<br>CTTCACACCACACTCCATCTA |
| Small outer H2_2 | AAGCAGGGTGGCGCGGGCCTCTTCGCGTGCCATTGAGCCT<br>CTTCACACCACACTCCATCTA |
| Small outer H2_3 | TAGCAATTAAGTTTTTCGCCATTCAGGCAGCGGGGATACCGG<br>CTTCACACCACACTCCATCTA |
| Small outer H2_4 | CCCGAAATATCAAAACGGCGGATTGACAAAAGGGCAAGGA<br>CTTCACACCACACTCCATCTA |
| Small outer H2_5 | CAGAAGCCAACAGTAAATGTGAGCGAGTTGATATTCATTATGAC<br>CTTCACACCACACTCCATCTA |
| Small outer H2_6 | CAATCGGTTTAGGGTGAAGGGATAGCTGTGTACGCGGGGA<br>CTTCACACCACACTCCATCTA |
| Small outer H2_7 | GTTACAGCGAGGGATAACCTCACCGGGTTCCGGGCTAACT<br>CTTCACACCACACTCCATCTA |
| Small outer H2_8 | ACCAACGTTGAAGCGGCCAGTGCCAAGCTCCGGCCGAGCCGG<br>CTTCACACCACACTCCATCTA |
| Small outer H2_9 | CCACTATTAAAGAATAGCTGTGCGTGGT CTTCACACCACACTCCATCTA |
| Small outer H2_10 | AGCAAACGAAGGAAACCGTCTATCACAGGTTGAGGAGCAGCACC<br>CTTCACACCACACTCCATCTA |
| Small outer H2_11 | AAAAGATTTTAAGATAGAACCCTTCTGCCAGCAGAGGTGT<br>CTTCACACCACACTCCATCTA |
| Small outer H2_12 | CGGTCAGTATCAAAGCGAACCAGACCGGTTTAGAAGTGTAGG<br>CTTCACACCACACTCCATCTA |
| Small outer H2_13 | AATCAATAGATTAAGAGTACCTTTAATTTACTTTTGAAATCACC<br>CTTCACACCACACTCCATCTA |
| Small outer H2_14 | CGGAATCAACTTTTTGAGACGGGCAACATTTGCGTACAGCAGTT<br>CTTCACACCACACTCCATCTA |
| Small outer H2_15 | TAAGAACAAATAAAGGTTCCGAAATCGGATGAGTGACAAACGCG<br>CTTCACACCACACTCCATCTA |
| Small outer H2_16 | GGAGGTTCTAACGACCCGAGATAGGGTTAACATACAGAGCAC<br>CTTCACACCACACTCCATCTA |
| Small inner core_1 | GTGAATTTCAGAGAGATAAC |
| Small inner core_2 | TGAAAATAAGACTCTTATTAGCGTTTG |
| Small inner core_3 | GTAAATGAATTACTAAAACACT |
| Small inner core_4 | AGAACCGCCCAATAGTGAAT |
| Small inner core_5 | TGTTTAACGTCCCGGAATAGGTG |
| Small inner core_6 | GAAGCGCATTAGACGTATTCATTAAAGGAACCAAGTACAAAACAGG |
| Small inner core_7 | GACGACGATAAAAACCATGG |
| Small inner core_8 | GGAAACAGTACATTCCACAGACA |
| Small inner core_9 | CCAAAGACAAAAGGCTAA |
| Small inner core_10 | ACGAGAAACACCAGAAAGAAGTTGGAATTG |
| Small inner core_11 | GCCGTCAATAGTAACATTATCA |
| Small inner core_12 | GGTAATGAATAATGGAATCGTCGACACTGA |
| Small inner core_13 | ACGCCAACATGTAAAGAA |
| Small inner core_14 | GTAGAAACCAATTTGAGGCAGGT |
| Small inner core_15 | ACCAGTAGCCGCCCGGGATCGTCACCCT |
| Small inner core_16 | AATCAGTAGATTTCAACTTTAA |
| Small inner core_17 | AGACTGTAGCGCACAAAGCTG |
| Small inner core_18 | GATTATCAGATGCATAAGGGAAC |
| Small inner core_19 | CGCCTGTGCAGTCCGTATAAACAGTTAA |
| Small inner core_20 | AGGAGGCAATAATATTTTCATCGTAGGAATCATTACC |
| Small inner core_21 | AGCCGCTATCATTTCGAGAACAAGCAAGGCCT |
| Small inner core_22 | CCACCAACTTGAGAAATATTGACGGAAATGGAGAATTACCATCG |
| Small inner core_23 | AGAACCGAGCCAGCAACCGATTGAGGGACAAAGTCGTTGCGC |
| Small inner core_24 | TCCCTCAGAGCACCATTA |
| Small inner core_25 | CGGTCATTGCCTTTAAATTGGGCTTGAGAAAATAGTTCACGT |
| Small inner core_26 | CTTGACCCGAACGAGTATTAGACTTTACGCG |
| Small inner core_27 | ATAGGCTGGCTGACCTTCAGTTTGAGATAATA |
| Small inner core_28 | GTCAATATGGCAATGCACGTAAAACAGAAATAAAGAA |
| Small inner core_29 | CCGGAACCTGATTGAACCTACCATATCAGAG |
| Small inner core_30 | TCATACCGACAATAGAGAATATAAAGTACCCCT |
| Small inner core_31 | TTTACCGCTAATGGAGGCATTTTCGAGCCGA |
| Small inner core_32 | ATGGAAAGCTTATCAACA |
| Small inner core_33 | GGCTGAGACTCCTCAAGGTATAGCAAAAATGA |
| Small inner core_34 | TTAGCGCCGTCGACAGAGAGAATAACATACGCACTCACCAAGAA |
| Small inner core_35 | GAGGCTGACAACAAACTGAAC |
| Small inner core_36 | AGGCCGACCGATAAGAGGGTAATTGAGCGGCGACATTCAAAATC |
| Small inner core_37 | GAGGACTACTCCAAAAAA |
| Small inner core_38 | GAGGAATAATTTTCGAGAGGCTTTTGCAAACGAGTAGTAGCGTC |
| Small inner core_39 | CGGGTAAAACTAAATTGCCAGAGGGGGTATTGCCCTG |
| Small inner core_40 | CACTACGGAGTGAATGTTTAGACTGGATAAAACAATTCGACAATG |
| Small inner core_41 | TAAAACACAACTTATACTGCGGAATCGTCGAT |
| Small inner core_42 | AGAATACTTCTGTATCATTGAATCCCCCTCAAATGC |
| Small inner core_43 | TTTGTATCATCGCCTGACTGTAGCATAAATCA |
| Small inner core_44 | AATCCGCACCAGTTAACCTTGCTTCTGTAAAGGGTTAGTTTGGA |
| Small inner core_45 | GTGTACTTACCGTACTATTAATTAATTTTCGACAAAA |
| Small inner core_46 | TAACGGCAAGCCCATCCTTGAAAACATAGCAGT |
| Small inner core_47 | AGTAACCACCCTCTTAGATTAAGACGCTGTTTAGGCACAGAACG |
| Small inner H1_1 | TTAGAGGGTTGATATAAAGAAGGATTGAC<br>AAATTATCTACCACAACTCAC |
| Small inner H1_2 | CCCACGCATAACACCAGGCGGATAAGTGGGGTTTCACCAG<br>AAATTATCTACCACAACTCAC |
| Small inner H1_3 | GTCGATATATTCGGAGCCG AAATTATCTACCACAACTCAC |
| Small inner H1_4 | CGACAATTGCAGGTCAGAG AAATTATCTACCACAACTCAC |
| Small inner H1_5 | TATCTTAAACAGCTTGATCTTTTGACCCTC<br>AAATTATCTACCACAACTCAC |
| Small inner H1_6 | CGAATAAGTTTCCCATTTT AAATTATCTACCACAACTCAC |
| Small inner H1_7 | AATAAGGCATAGTAAAGAATAGAAAGGAACAATACGTCAACG<br>AAATTATCTACCACAACTCAC |
| Small inner H1_8 | TCCATCAACAGTTTCAGCGAAGGCGGATAT<br>AAATTATCTACCACAACTCAC |
| Small inner H1_9 | CATTTGAGATAAATATTGGGATTTTGCTAAGAAAGAAGTAAT<br>AAATTATCTACCACAACTCAC |
| Small inner H1_10 | TGAAACAAACTACAACGCTAAATTCAGACG<br>AAATTATCTACCACAACTCAC |
| Small inner H1_11 | GATACAGGATGTTACTTAG AAATTATCTACCACAACTCAC |
| Small inner H1_12 | TAGAAATAGGAACCCATGGGTAATTTTGAT<br>AAATTATCTACCACAACTCAC |
| Small inner H1_13 | TAGCATTTTCAGGGATAGGGTCAGTAAGCG<br>AAATTATCTACCACAACTCAC |
| Small inner H1_14 | GAGTACCCTCAGAGCCACAGTGCCTCTGAA<br>AAATTATCTACCACAACTCAC |
| Small inner H1_15 | AAATAGCACCGTTTTTCGGCTGTCTTTCCTCGCCAGCATTAGGA<br>AAATTATCTACCACAACTCAC |
| Small inner H1_16 | CCACCAGAACCACTGCTCA AAATTATCTACCACAACTCAC |
| Small inner H1_17 | CGGGTATTATGAATTATCACCGTCACCGCCCTCAGTCGCT<br>AAATTATCTACCACAACTCAC |
| Small inner H1_18 | ACCCTGAAGGGAAGGTCCATTTGGGAATTAGCCACCCGAGTTAA<br>AAATTATCTACCACAACTCAC |
| Small inner H1_19 | TTTACGACAGAATCAAGTTAGCCCCTTTTCAT<br>AAATTATCTACCACAACTCAC |
| Small inner H1_20 | TAGTTTTCATCGGATTAAA AAATTATCTACCACAACTCAC |
| Small inner H1_21 | TCATTACCCAAATAATGC AAATTATCTACCACAACTCAC |
| Small inner H1_22 | CTCATTCAGCTCGTATTAAATCCTTTGCAAGAACCACCAACC<br>AAATTATCTACCACAACTCAC |
| Small inner H1_23 | TTAGATTATTAATTTTAAAATCAAGGGCAAA<br>AAATTATCTACCACAACTCAC |
| Small inner H1_24 | ATATATGTAAATTATTTTCATCAATATAATCGAGGCGGTGTCGA<br>AAATTATCTACCACAACTCAC |
| Small inner H1_25 | GTTTCGTCGACCTGCTCCA AAATTATCTACCACAACTCAC |
| Small inner H1_26 | TTATACTTCAGTAATTCTGTCCAGACGAATGGCTAAGTTT<br>AAATTATCTACCACAACTCAC |
| Small inner H1_27 | AATAAAACAACATGTTCAGTTCCAGTGCCTTG<br>AAATTATCTACCACAACTCAC |
| Small close_1 | TAACAATTTCATTTGAATTACCTTTTTTATAAACAGTTCAGAAAACGAGA<br>AAGGAGTT |
| Small close_2 | AGAGCAACACTATCATAACCCTCGTTTACACAAGAATTGAGTTAAGCCCA<br>GAACGATA |
| Small close_3 | AGGGCTTAATTGAGAATCGCCATATTTAAGCCCAATAGCAAGCAAATCAG<br>TTGGTTGG |
| Small close_4 | ATCAATAGAAAATTCATATGGTTTACCAGATTGTGAATTACCTTATGCGA<br>ACCATGAG |
| Small close_5 | CTTTAGGAGCACTAACAACTAATAGATTATGCGTAGATTTTCAGGTTTAA<br>CTGTTTGC |
| Small close_6 | TTTATCCCAATCCAAATAAGAAACGATTTATCAAAATCATAGGTCTGAGA<br>GGAACCTA |
| Small anti-close_1 | AACTCCTT TCTCGTTTTCTGAACTGTTTATAAAAAAG |
| Small anti-close_2 | TATCGTTC TGGGCTTAACTCAATTCTTGTGTAAACGA |
| Small anti-close_3 | CCAACCAA CTGATTTGCTTGCTATTGGGCTTAAATAT |
| Small anti-close_4 | CTCATGGT TCGCATAAGGTAATTCACAATCTGGTAAA |
| Small anti-close_5 | GCAAACAG TTAAACCTGAAAATCTACGCATAATCTAT |
| Small anti-close_6 | TAGGTTCC TCTCAGACCTATGATTTTGATAAATCGTT |
| Small open_1 | TATCACCGTACTCAGGAGGTTTAGTACCGCCACCCTCAGAACCGCCACCCTC<br>CCAACATC |
| Small open_2 | CAGCAGCGAAAGACAGCATCGGAACGAGGGTAGCAACGGCTACAGAGGCTTT<br>TTAGAAGG |
| Small open_3 | CGAACTGACCAACTTTGAAAGAGGACAGATGAACGGTGTACAGACCAGGCGC<br>GCGAAAGT |
| Small open_4 | CAGACGATTGGCCTTGATATTCACAAACAAATAAATCCTCATTAAAGCCAGA<br>AAAGGTTT |
| Small open_5 | CCATTAGCAAGGCCGGAAACGTCACCAATGAAACCATCGATAGCAGCACCGT<br>TTGTAGGA |
| Small open_6 | CCATCTTTTCATAATCAAAATCACCGGAACCAGAGCCACCACCGGAACCGCC<br>CTATGATG |
| Small open_7 | GCCCTCATAGTTAGCGTAACGATCTAAAGTTTTGTCGTCTTTCCAGACGTTA<br>GATTACGA |
| Small open_8 | ATAGATAAGTCCTGAACAAGAAAAATAATATCCCATCCTAATTTACGAGCAT<br>ATGTTTGC |
| Small open_9 | CATCTTTGACCCCCAGCGATTATACCAAGCGCGAAACAAAGTACAACGGAGA<br>TCCTAGCA |
| Small open_10 | TGCCCCCTGCCTATTTCGGAACCTATTATTCTGAAACATGAAAGTATTAAGA<br>AGAACTGG |
| Small open_11 | AAGGCTCCAAAAGGAGCCTTTAATTGTATCGGTTTATCAGCTTGCTTTCGAG<br>CAGTGGTA |
| Small open_12 | TTTTGCGGAACAAAGAAACCACCAGAAGGAGCGGAATTATCATCATATTCCT<br>GCTAGGTA |
| Small anti-open_1 | GATGTTGG<br>GAGGGTGGCGGTTCTGAGGGTGGCGGTACTAAACCTCCTGAGTACGGTGATA |
| Small anti-open_2 | CCTTCTAA<br>AAAGCCTCTGTAGCCGTTGCTACCCTCGTTCCGATGCTGTCTTTCGCTGCTG |
| Small anti-open_3 | ACTTTCGC<br>GCGCCTGGTCTGTACACCGTTCATCTGTCCTCTTTCAAAGTTGGTCAGTTCG |
| Small anti-open_4 | GAACCTTT<br>TCTGGCTTTAATGAGGATTTATTTGTTTGTGAATATCAAGGCCAATCGTCTG |
| Small anti-open_5 | TCCTACAA<br>ACGGTGCTGCTATCGATGGTTTCATTGGTGACGTTTCCGGCCTTGCTAATGG |
| Small anti-open_6 | CATCATAG<br>GGCGGTTCCGGTGGTGGCTCTGGTTCCGGTGATTTTGATTATGAAAAGATGG |
| Small anti-open_7 | TCGTAATC<br>TAACGTCTGGAAAGACGACAAAACTTTAGATCGTTACGCTAACTATGAGGGC |
| Small anti-open_8 | GCAAACAT<br>ATGCTCGTAAATTAGGATGGGATATTATTTTTCTTGTTCAGGACTTATCTAT |
| Small anti-open_9 | TGCTAGGA<br>TCTCCGTTGTACTTTGTTTCGCGCTTGGTATAATCGCTGGGGGTCAAAGATG |
| Small anti-open_10 | CCAGTTCT<br>TCTTAATACTTTCATGTTTCAGAATAATAGGTTCCGAAATAGGCAGGGGGCA |
| Small anti-open_11 | TACCACTG<br>CTCGAAAGCAAGCTGATAAACCGATACAATTAAAGGCTCCTTTTGGAGCCTT |
| Small anti-open_12 | TACCTAGC<br>AGGAATATGATGATAATTCCGCTCCTTCTGGTGGTTTCTTTGTTCCGCAAAA |
| Large miniscaf_1 | CCGAGCGGGTTGATGGCTCAAGGGCGCTTCAAGTCAAGGGGTTAACCGT |
| Large miniscaf_2 | GCTCAACCACTACCGCACGGAATCTGGGCACGGTGCTACGACAGACTCG |
| Large miniscaf_3 | CGGCGGGTTGATTGGCAGCAAATACCGGCACTGGCACATGAGGCGAGCG |
| Large miniscaf_4 | CTCCGGCACTGCGCGGCGTTGTGTCTGCTGTCTCTTTTGAGGACTACCT |
| Large miniscaf_5 | TGTCGCTGCACAGTACGAGCTTCCAATCGCCGGGACAACCGCCCGGCAC |
| Large miniscaf_6 | TCGATCCTGAGCGAACTCACCTCCTCGGGCATGGTCGACGCTGCCGTTC |
| Large miniscaf_7 | TGGCCCGATGCTGGAGAGGATTGACGCGATGCAATACGCCGTCCTCCGT |
| Large miniscaf_8 | CGAGCAACCTACGCATCCAAACCCTAGTGTGGCCGGCCCAACGACGTTT |
| Large miniscaf_9 | GTCTTCTGAGGGCGCGCGGCTACCGGTACTAGACGGAGGACTGCTGGTC |
| Large miniscaf_10 | GCCTACCACCTCCCGTCGACCCATCTCGCCCGTAGCTCAAGAGAACAAG |
| Large miniscaf_11 | CCGTGCCCACCCACCCAGCGCTGATGAACCTTATGCAGACCCAGTTCAG |
| Large miniscaf_12 | CTGGGCGAGTGGGAGTCGCGGCGTTGTGAACACGACCTATGCAGGAGAC |
| Large miniscaf_13 | AATTTCGTGACGCGCAAGGCCCGGTCTCACTCTCCGTAGCGTCGGCCAC |
| Large miniscaf_14 | GGCACGCTCCCAGGCCGCAGACGACTTCGAACGTTTGAAGGACAGGAAT |
| Large miniscaf_15 | CACTGGGAGGTCGGTTGGTAGCGTGCTCGCGTGCCACTTAGGCCGTGAA |
| Large miniscaf_16 | CGGCGCACACCCAAGTGACAGCGTAATGGCAATGCCCACTGAACAAGTC |
| Large miniscaf_17 | AATAGAGGGCGCGTCGCGAGGATTGTGGCAGCCAGGGAATGCGGCTTCA |
| Large miniscaf_18 | CCTACGGCACGCTGCGAGCAGGCAGGACTGGAAGATTGACCCGACCGTG |
| Large miniscaf_19 | TTGTGGGTCTGTTCGCCTCAACTCTGCTGCGTCGGTGCGATGTGTAGGC |
| Large miniscaf_20 | TGAGGAGTCCCTCGAAAGCCGAGCCAACGTCCTCGCACTCAGGAAATAT |
| Large miniscaf_21 | CTCGATGTCGGGATGGTGACGCGCGGACACGCTTGAGAAGCGCGACCAG |
| Large miniscaf H2_1 | TCGGGCCCGTCATCGTCTCATATCCTCAGCAACTGCGGG<br>CTTCACACCACACTCCATCTA |
| Large miniscaf H2_2 | GGGCGACTGGCTACCGCTGCTACCCAGATATCACCGCAC<br>CTTCACACCACACTCCATCTA |
| Large miniscaf H2_3 | TCGTGGGCAAGATATATGTGGTCCGGGATGGGCGGTGCG<br>CTTCACACCACACTCCATCTA |
| Large miniscaf H2_4 | CGGCCCACAGGGTCAATAGCCGTCGCCTCACCCGACTTA<br>CTTCACACCACACTCCATCTA |
| Large miniscaf H2_5 | GAACCGCTAGCCTCGGCCGCGCATTTACACCTTGGTCCA<br>CTTCACACCACACTCCATCTA |
| Large miniscaf H2_6 | CGGTCCAGCACCATGGTCCCGTACCCTCACTCGCCATTG<br>CTTCACACCACACTCCATCTA |
| Large miniscaf H2_7 | GCACAACTCCTGAACAAGGCGCCCAGTGCAGTCGACGTC<br>CTTCACACCACACTCCATCTA |
| Large miniscaf H2_8 | CGACTAGAGCTGCTGTGCGTCCCGTTGCGTGAGTTAGGC<br>CTTCACACCACACTCCATCTA |
| Large miniscaf H2_9 | CCTACCCTCGGTGCCGAGTCCCTAGACGCTGGGCCAGTA<br>CTTCACACCACACTCCATCTA |
| Large miniscaf H2_10 | CGGCCACCGCTCGGATTATCCACAGATTATGGCGCGAGG<br>CTTCACACCACACTCCATCTA |
| Large miniscaf H2_11 | CCTTGAGCCGCATGCGCTACGTCTTGGCCTCCTACCCTC<br>CTTCACACCACACTCCATCTA |
| Large miniscaf H2_12 | GCGCAGTCCCGTTCTCGAGTGTGCCCACTTGCCCTCAAT<br>CTTCACACCACACTCCATCTA |
| Large miniscaf H2_13 | ACTCCCTGATAGGCGTCTTTCAGGCGGGCGGGACTCAGG<br>CTTCACACCACACTCCATCTA |
| Large miniscaf H2_14 | GGGACGCCAGACTGGGCTCACGAGTGATAGGCTTTCCGC<br>CTTCACACCACACTCCATCTA |
| Large miniscaf H2_15 | TTCGGCTTAGGTCCTTCTCGTGTGGTGCCCGCTGCCCTT<br>CTTCACACCACACTCCATCTA |
| Large miniscaf H2_16 | GCGAGCCACGCGGCGTGTAACCAACAGCATTTTCTCGCT<br>CTTCACACCACACTCCATCTA |
| Large miniscaf H2_17 | TGCGGCAGCATCAAGGACCTACCATCTTGACCGGTCGCC<br>CTTCACACCACACTCCATCTA |
| Large miniscaf H2_18 | ACTCCAGTTGCTAGAACGGTCTCGCGCACACACGGTGGG<br>CTTCACACCACACTCCATCTA |
| Large miniscaf H2_19 | CGTGCTTTGATCCGACTGACACTGCGTGGCCGATGGGCT<br>CTTCACACCACACTCCATCTA |
| Large miniscaf H2_20 | GTGCCCGTTGTGAGTCGGAGAACCTTGCGCGGCCTATTG<br>CTTCACACCACACTCCATCTA |
| Large miniscaf H2_21 | CGCAGGGACGCCGTCGAAACACAGCACTCCACATCGTAC<br>CTTCACACCACACTCCATCTA |
| Large outer core_1 | CAGAAGGCGTCTATCAATAGCAGCCTTCATCCTAATCCAC |
| Large outer core_2 | TTTGTATCAGATGATGGCGTTGAGCGTGCGCCTTGACTTGTCCT |
| Large outer core_3 | GAAGATTATACTTCTGACCCGCCGCGCACGTAGCACCGTTTGT |
| Large outer core_4 | TAACAAAATTATTTGCAGCCGGAGTAAGTATGTGCCAGTACCA |
| Large outer core_5 | CAAGATTTTCAGGTTTAAGCGACATGGACCAAAAGAGACTTGC |
| Large outer core_6 | ATGCTTTTACATCGGGAGGATCGACAATGGTTGTCCCGGACAG |
| Large outer core_7 | GGCTTGAATACCAAGTTCGGGCCAGACGTGTCGACCATGGATT |
| Large outer core_8 | GGGCGATAAAATCGCGTAGTATCATATATCAGCTCAGGAA |
| Large outer core_9 | AGGCCAGCTGGCGAAAGGTTGCTCGGCCTAGCGTATTGCAATTA |
| Large outer core_10 | AAAGTAACGCCAGGGTTAGAAGACTACTGGGGCCGGCCAAAGT |
| Large outer core_11 | TTGGCCAGTGCCAAGCTGGTAGGCCCTCGCCTCCGTCTAAACG |
| Large outer core_12 | AGAATAACCTCACCGGAGGCACGGGAGGGTGAGCTACGGGGAA |
| Large outer core_13 | GACTTTACCAGTCCCGGCGCCCAGATTGATCTGCATAAGCGCC |
| Large outer core_14 | TGCAGGCTTTGTGAGAGGGTAGCTATTCAGCATTGATCAT |
| Large outer core_15 | GTCCGGTTGCGGTATGAGCGAAATTCCTGATAGGTCGTGTCGGG |
| Large outer core_16 | GGTATGGGTAAAGGTTTGCGTGCCGCGGACTACGGAGAGGCTG |
| Large outer core_17 | TCGTGGTGTGTTCAGCACCCAGTGAAGGGTTCAAACGTTCTTA |
| Large outer core_18 | GGTACCTGCAGCCAGCGTGCGCCGAGCGAAAGTGGCACGCCGG |
| Large outer core_19 | TCTCAGCGCAGTGTCACCTCTATTGGCGAGTGGGCATTGGACG |
| Large outer core_20 | TGATTGCCGCGCCTGTGACAGAATCAACAAAGTTACCAGC |
| Large outer core_21 | ACCATGCAGCAAGCGGTCCCGTAGGCCCACTTCCCTGGCTAGAG |
| Large outer core_22 | GCGTCCTGTTTGATGGTCCCACAAAGCCCTCAATCTTCCGCGA |
| Large outer core_23 | AGGAATCAAAAGAATAGCTCCTCACAATACGCACCGACGCCTT |
| Large outer core_24 | TGGTTTGGAACAAGAGTCATCGAGGTACGAGTGCGAGGAGTTC |
| Large outer core_25 | GATTCAAAGGGCGAAAACGCTCGGCCCGCTTCTCAAGCGTCCA |
| Large outer core_26 | ATATAAGCGCCGGCCCGAACGGTTAACCTGAACCATGTAGAAACCACAT |
| Large outer core_27 | GATCGGAACAAAGAAATTACGAGAAGAAAA |
| Large outer core_28 | TTGCGTTATTAATTTTCCTT |
| Large outer core_29 | ATGGTATTAAATCCTTTGTAT |
| Large outer core_30 | TATATCAATATATGGAGCGA |
| Large outer core_31 | AAAATTAATGGAAACAGATAT |
| Large outer core_32 | GTATTTCATTTGAATTTTCT |
| Large outer core_33 | TCAGAAATTAATTACATTTTG |
| Large outer core_34 | TACAAACAAACATCAAATAA |
| Large outer core_35 | ACAACTGAGCAAAAGAAGAAT |
| Large outer core_36 | GCTGAATTATTCATTTATTA |
| Large outer core_37 | CGCTTCGCGTCGTTGTGCACGGAGGAAATTCGCACCAATAGGAACGTTC |
| Large outer core_38 | CGCTGCGCAACTGTTGTTTTTTAATTAAAT |
| Large outer core_39 | ATGTGCGCCATTCGCCACCAT |
| Large outer core_40 | TGGAGTGCCGGAAACCCTGG |
| Large outer core_41 | CCAGGTGTACATCGACATAAA |
| Large outer core_42 | ACGCCGCCAGCAGTTGTGTG |
| Large outer core_43 | AGAGCTTAAATTTCTGCAGGG |
| Large outer core_44 | CGGCGCAGAAACAGCGCAGT |
| Large outer core_45 | AATCAGCTCTCACGGAATGAT |
| Large outer core_46 | ATGTTTCTCCGTGGTGTGAT |
| Large outer core_47 | CCGCTCACAACACTGCGCGTCTCCTGAGAACCATTGAGGCAGGTCAGGT |
| Large outer core_48 | CGCAGCATCAGCGGGGCAGGAGGCCACCAG |
| Large outer core_49 | GGTCGCTTACGGCTGGAGACG |
| Large outer core_50 | GTAGGTGCCATCCCACACAA |
| Large outer core_51 | TGCTGTTGAGGATCCACGTCA |
| Large outer core_52 | CACCGTCCGTGAGCCTGGGA |
| Large outer core_53 | CGTTGCACGCGTGCCTGTCAC |
| Large outer core_54 | GTTCATACCGGGGGTTATTA |
| Large outer core_55 | CCGGCGGCGGGCCGTTTCACC |
| Large outer core_56 | ATCGTGGTGCTGCGGCGCAG |
| Large outer core_57 | CCTGGCCACAATGCCGCATGAAGCCGAGAAAAGAAACCGAGGAAACTTC |
| Large outer core_58 | AGTGTGAGACGGGCAACAGAAGGTAAGCAG |
| Large outer core_59 | GCTGGGTGGTTTTTCTTGCAA |
| Large outer core_60 | AAAGTTTGCGTATTGGA AAA |
| Large outer core_61 | TCCGGCAACGCGCGGGGGACT |
| Large outer core_62 | TCATTGAGTAACATTATATCA |
| Large outer core_63 | ATAGAGCCCCCGATTTGTCT |
| Large outer core_64 | GAGACTAAATCGGAACCGCCA |
| Large outer core_65 | CAGGGTCGAGGTGCCGCCAT |
| Large outer core_66 | CTATACCCAAATCAAGTCAAA |
| Large outer core_67 | ACGGGCCCACTACGTGGTTT |
| Large outer core_68 | AGAGTCGCCCCGAGTCTGCAGCTAAATTCCAAGAACGGGCCC |
| Large outer core_69 | GTACCCACGACGCTCGCCGTAAAGTAACTTTTTCAAATTACA |
| Large outer core_70 | GACTGGGCCGAGGTAGTCTTTTCGACTAAATTTAATGGTTAA |
| Large outer core_71 | TGGGCGGTTCGTGCCGGGCGCCATAAGGCGTTAAATAAGATG |
| Large outer core_72 | AGGTGGACCGGAACGGCACAGTATAGAAAAAGCCTGTTCAGA |
| Large outer core_73 | AGGCTAGTCGAAACGTCGTATAAGCTCCTCATATATTTTAAA |
| Large outer core_74 | CGGGGGTAGGGACCAGCACCCCGGTGTAAAGATTCAAATCAT |
| Large outer core_75 | GATGTGGCCGCTTGTTCTCGATGAAATCACCATCAATAAAAG |
| Large outer core_76 | ATCCTCAAGGCTGAACTGCTATCAGTTAATGCCGGAGAGATA |
| Large outer core_77 | ACCAGGGAGTGTGGCCGAAGAACCGAATAAATCCTACCCGTC |
| Large outer core_78 | GTCGCGTCCCATTCCTGTAGAGCCAAGAGCCAGCAAAATTCT |
| Large outer core_79 | CACAGCCGAATTCACGGCATTAGCGAGGCCGGAAACGTTCAC |
| Large outer core_80 | TACGGCTCGCGACTTGTTCTGTAGCCGTAATCAGTAGCGCAC |
| Large outer core_81 | CTGCTGGAGTCACGGTCGAACAATGCTGGCATGATTAAAGAG |
| Large outer core_82 | AGAAAGCACGGCCTACACTAACCCACAGAGCCTAATTTCTAA |
| Large outer core_83 | GGCCGGGCACATATTTCCTGAACAAATTTATCCCAATCTTTT |
| Large outer core_84 | CGCCCCTGCGCTGGTCGCAAACAGGGTCAAAAATGAAAGGGC |
| Large outer core_85 | AGAGAATAACATAAGCAGTTGCTGAGGATATGAGACCCTACAG |
| Large outer core_86 | ATAATATGATGACGCTTGAGCCATCAACCACAGCGGAATTATCATC |
| Large outer core_87 | ATACAAATTCTTACGCCGACTGCACTGGGCGCCTTGAAGCGTT |
| Large outer core_88 | TTTTGTTTTCAGGAAATCCTCTCCAGCATACCGGTGCGGGCCTCTT |
| Large outer core_89 | GAGATCTACAAAGGGGGGGCAAGTGGGCACACTCGAGCTTTGA |
| Large outer core_90 | AGCCGCCGAACGGGGCCGCGACTCCCACTAAGCTTTCGCACTCAAT |
| Large outer core_91 | CCTTTAGCGTCAGACACCGGTCAAGATGGTAGGTCCAAGTTTG |
| Large outer core_92 | ATAGCCGTTGATGCTCCTCGCGACGCGCCTGCCTTCACCGCCTGGC |
| Large outer core_93 | ATAAACAATAGATAAGTCCGTGATATCTGGGTAGCAGCGGTTTATCATCGGC |
| Large outer core_94 | TGTCTTTAAAAGTTCAATATAATCCTGAGCCC |
| Large outer core_95 | ATCTGCAGAACGCGCCTGTAGCCATTCCGTGCGGTAGTGAAT |
| Large outer core_96 | TAAATAAACAACATGTTTCCGCCCATCCCGGACCACATAGACGACAACCAAG |
| Large outer core_97 | TACCAACCAACTCGAAGGGTTAGAACCTGCCG |
| Large outer core_98 | GAAAATTCTGTCCAGACTATCTTGTTTGCTGCCAATCAAATA |
| Large outer core_99 | TTTAGTACCGACAAAAGTCCGGGTGAGGCGACGGCTATTAATATAAAGTTAA |
| Large outer core_100 | TTTCATCACCTTTTCAGAAATAAAGAAAAGCA |
| Large outer core_101 | GACGCCAGTAATAAGAGGACCCTGACAACGCCGCGCAGTCGT |
| Large outer core_102 | AAATTTAGGCAGAGGCACTCAAGGTGTAAATGCGCGGCCCATGTAATACCGA |
| Large outer core_103 | CCGTGTGGAAAACAATGAATATACAGTACGAT |
| Large outer core_104 | ATATTTAACAACGCCAAGAGGCTAAAGCTCGTACTGTGCACG |
| Large outer core_105 | AAAGCTTAATTGAGAATCGGCGAGTGAGGGTACGGGACCCAGTAGGCACCGG |
| Large outer core_106 | AATCATACAATTACTAACGGATTCGCCTCCCG |
| Large outer core_107 | CTAAAGCCAACGCTCAAATGGTGCAGGTGAGTTCGCTCAGAA |
| Large outer core_108 | CAATAATATTTTGTTAACGACTCACGCAACGGGACGCACTAAACGTAAATAA |
| Large outer core_109 | TTCGCGTAGGCAAAGCTGCAAGGCGATTCACT |
| Large outer core_110 | CCTAAATATTTAAATTGAGCAGCTGTTTGGATGCGTAGGGGG |
| Large outer core_111 | TGCAAACAGGAAGATTGTTGCCCAGCGTCTAGGGACTCGGCCCCAAAATGCC |
| Large outer core_112 | TGAGTAAGGCGGTTTCACGACGTTGTAAGTAC |
| Large outer core_113 | TAGTGATAATCAGAAAAGCACCGATAGCCGCGCGCCCTCTTC |
| Large outer core_114 | TGAGTCAATCATATGTAGTCGCCATAATCTGTGGATAATCTAGCATGAAAGG |
| Large outer core_115 | CCGGAGAGATCAAAGTGGAGCCGCCACGGCGA |
| Large outer core_116 | CAACGGTAATCGTAAAACCGAGCGGGGTCGACGGGAGGTTTC |
| Large outer core_117 | ATTGCAAACAAGAGAATCTTAGGAGGCCAAGACGTAGCGGTCTGGACAACCG |
| Large outer core_118 | TTCTAGCAAGGGATGGCGAAACGTACAGGTTC |
| Large outer core_119 | AAAGTCATTGCCTGAGACATGCGGAGCGCTGGGTGGGTGAAC |
| Large outer core_120 | ATTTCAGAGCCGCCACCCAGTCCCGCCCGCCTGAAAGACACCACCCGGCCTT |
| Large outer core_121 | GATATTCGCAACCAACTGTTGCCCTGCGTGAG |
| Large outer core_122 | ACACCACCCTCAGAGCCGCCTATCGGGCCTTGCGCGTCACCG |
| Large outer core_123 | CCGGAGCCGCCACCCTCCGAAGCCTATCACTCGTGAGCCTCCCTCAACTTGA |
| Large outer core_124 | GCCATTTCCTCACACGTCATAAACATCCCGAA |
| Large outer core_125 | ATTCCACCGGAACCGCCCAGTCTGGTCTGCGGCCTGGGACTT |
| Large outer core_126 | CAGAAATCACCGGAACCCCCAGCGGGCACCACACGAGAAATAATCATAGCAC |
| Large outer core_127 | CATTACCTCTGCCAAACGGCATCAGATGCGAG |
| Large outer core_128 | GCATTTGCCATCTTTTCGGACCTAGCTACCAACCGACCTAAT |
| Large outer core_129 | AATGTCATAGCCCCCTTCTGAAAATGCTGTTGGTTACACATTTTCGGAAACC |
| Large outer core_130 | ATCGATACAGAATGTGCCCCCTGCATCACCAT |
| Large outer core_131 | CACGCGTTTTCATCGGCGCCGCGTGCTGTCACTTGGGTGGTG |
| Large outer core_132 | TAACGAAGCCCTTTTTACACGTGTGTGCGCGAGACCGTTATCTTACTAACGG |
| Large outer core_133 | AATACCCGCGCCAGGTTTGCCCCAGCAGAGTC |
| Large outer core_134 | GAAAAATAGCAATAGCTCTAGCAACCTGCTCGCAGCGTGCAC |
| Large outer core_135 | CCTTAATAAGAGCAAGAGGATCGGCCACGCAGTGTCAGTAGCCCAATACGCT |
| Large outer core_136 | AACGAGCAGAGCTTAAATCGGCAAAATCCAGC |
| Large outer core_137 | TTCCAAGAATTGAGTTACGGATCAGTTGAGGCGAACAGAGGT |
| Large outer core_138 | GTTTAATATCAGAGAGAATGGCCGCGCAAGGTTCTCCGATGAGCGCACAAAA |
| Large outer core_139 | TAAACAGTAAAGCATAGGGTTGAGTGTTCGTT |
| Large outer core_140 | ATTAGTCAGAGGGTAATCTCACAATCGGCTTTCGAGGGACCC |
| Large outer core_141 | TAATTAACTGAACACCCTGATGTGGAGTGCTGTGTTTCGGGGAGAAGAAACG |
| Large outer core_142 | ATTTTTTAACCATCTAAAGAACGTGGACTGTC |
| Large outer core_143 | AACGAAGCGCATTAGACACGGCGTGCGTCACCATCCCGACCA |
| Large inner core_1 | GCGACACCCGCCGCGCCTAAAATATTATC |
| Large inner core_2 | AAAAGGAGTCAGAAGATTAGTTGCTATTTGGAA |
| Large inner core_3 | AATGCAATCAAAAGAGGTTTTGAAGCCTGCTA |
| Large inner core_4 | TAGGAATCCCGAAATAGCGAACCTCCCGACGGT |
| Large inner core_5 | CGATATCGTTTTATATTCTAAGAAC |
| Large inner core_6 | AACCATCGCGAACCCAGATATAGAAGGCTCTTT |
| Large inner core_7 | TTGCGCCTCCAACACCGCGCCCAATAGCAGAT |
| Large inner core_8 | AAACAGCGAGTACCTATTTTCATCGTAGGATAC |
| Large inner core_9 | AACTAACCCTCGTTGCGAAAGGAGCGGGCTAAA |
| Large inner core_10 | TACAGGCGAGCACCGCTGGCAAGTGTAGCTTG |
| Large inner core_11 | CAGTTGACGCGTACCTGCGCGTAACCACCAGGC |
| Large inner core_12 | GCGGGATTGAAAGGGCACTAACAACTAATAAGC |
| Large inner core_13 | AGCGAAGTCAGTTAGCCGTCAATAGATAAATC |
| Large inner core_14 | ACGAGGGCAAACCCGAGGATT |
| Large inner core_15 | GCTCCAATGCTGAAGTTATATAACTATATTAAT |
| Large inner core_16 | GAAAATTCAACATTTTTAACCTCCGGCTTTGA |
| Large inner core_17 | AATAATAACTAAAGTCATAGGTCTGAGAGGATT |
| Large inner core_18 | AACAACGTTTCATGTCACGTTGGGAATTTATC |
| Large inner core_19 | GCGGAGTGATTCCCCGGATTGACCGTAATTAAC |
| Large inner core_20 | ACAACTGTAGATTGGATTCTCCGTGGGAGGGG |
| Large inner core_21 | TCTGTATGATACATTGTGAGCGAGTAACATCAG |
| Large inner core_22 | CCATTAACACGCTGTTTCCCTTAGAATCCTAGG |
| Large inner core_23 | GTAATGTAACACCCATAGCGATAGCTTAACTA |
| Large inner core_24 | ACCAACCCAGAGGTCGCTGAGAAGAGTCA |
| Large inner core_25 | GCAAAAAACCACCTAGATGGGCGCATCGGGGA |
| Large inner core_26 | CACTCATCGCCATTCATCTGCCAGTTTGAACAA |
| Large inner core_27 | CGATTAGAACTGAACGACAGTATCGGCCACCC |
| Large inner core_28 | ACAAAGTGGCTATTATCGCAC |
| Large inner core_29 | TCATAGTAAAAGGTTTTTGCGGGAGAAGCATGA |
| Large inner core_30 | AGCATTTAATAGTAAAACATTATGACCCCTGA |
| Large inner core_31 | GTACAAACCAATAATAAAGCTAAATCGGTCGGT |
| Large inner core_32 | ATAGGAAAACGGGGACAGGAGTGTACTGGCATC |
| Large inner core_33 | TTCAGGAACAGTGCGTCATACATGGCTTGACT |
| Large inner core_34 | TCAGAGCTAATGCCCTGAATTTACCGTTCTGCT |
| Large inner core_35 | CTCCATGGAGATAGTAAAGTTAAACGATGTGTA |
| Large inner core_36 | ACGAGGAAAGGGACCGTTCCGGCAAACGTGTA |
| Large inner core_37 | TCATAAGCCAGTCATTTTTCGTCTCGTCGTCAG |
| Large inner core_38 | CCAACTTTATTTACCTCCGGCCAGAGCATACAAAATTAAGCAA |
| Large inner core_39 | GATGAACCGCTCAATAACGGAACGTGCCGTTGA |
| Large inner core_40 | GGCGCAATGGAAAAGAACGTCAGCGTGGCAGT |
| Large inner core_41 | CTTCATCATTGCAATGGTCAG |
| Large inner core_42 | CCGTACTGACTCCTGGGAGGGAAGGTAAAGCTG |
| Large inner core_43 | GCCCGGGGATTAGAAGGGCGACATTCAAATCC |
| Large inner core_44 | AGAGGGTGTACCAGGGTTTACCAGCGCCACATA |
| Large inner core_45 | TTTTGCCAATCGTCCCACGGAATAAGTTTAGTG |
| Large inner core_46 | GAGGCTAATCCCCCATATAAAAGAAACGCGTT |
| Large inner core_47 | GATAAAAACAGTTCAATACATACATAAAGCCAG |
| Large inner core_48 | GCTGCTCAGTAGAACTGTGTGAAATTGTTCCGA |
| Large inner core_49 | GGCTTGAGTAATACACAATTCCACACAAAAGA |
| Large inner core_50 | CACCAGATGTAGCACCGGAAGCATAAAGTATTC |
| Large inner core_51 | TGGGCTTCCATCACTGGGGTGCCTAATGATTT |
| Large inner core_52 | TTTCAACGCCACCGAACTCACATTAATTGCAAA |
| Large inner core_53 | AATTACAGTGTTTCTCACTGCCCGCTTTGTGG |
| Large inner core_54 | AAGAACTCGGTACGGAAACCT |
| Large inner core_55 | ATATGCGGATAAGAGACTGGATAGCCACAATCA |
| Large inner H1_1 | GAAAAGAATCAGAGCGGGCGTTAATAGGCATA<br>AAATTATCTACCACAACTCAC |
| Large inner H1_2 | GGGGTATAACGTGCTTTGGAACAAACGCC<br>AAATTATCTACCACAACTCAC |
| Large inner H1_3 | CACGTATGGTTGCTTTGATAGAAAGTTCAACT<br>AAATTATCTACCACAACTCAC |
| Large inner H1_4 | GTTTGACTTCAAATATCGATTCGGTCGCTTT<br>AAATTATCTACCACAACTCAC |
| Large inner H1_5 | AGGAAATTGAGGAAGGTTAAAGGCCGCATAAC<br>AAATTATCTACCACAACTCAC |
| Large inner H1_6 | TAGGGCAAATCAACAGTCGTCACGACAAC<br>AAATTATCTACCACAACTCAC |
| Large inner H1_7 | ATTTTCAATCAATATCTGAGACAGCCCGATAG<br>AAATTATCTACCACAACTCAC |
| Large inner H1_8 | AAGGAAGTGCACCCCCCTGACTATTATAATTACGAAAACG<br>AAATTATCTACCACAACTCAC |
| Large inner H1_9 | TCAAGCAAAGCGGATTGCGATACATACATTAT<br>AAATTATCTACCACAACTCAC |
| Large inner H1_10 | CGGAGATTAAGAGGAAGACCACAATTCAT<br>AAATTATCTACCACAACTCAC |
| Large inner H1_11 | GGAGTTATTTAATGCGCCGCTACAGGGGAGAGGCTTGCAG<br>AAATTATCTACCACAACTCAC |
| Large inner H1_12 | CGGATTCGAGCTTCAAAGCCCACGCTTTT<br>AAATTATCTACCACAACTCAC |
| Large inner H1_13 | AAATAGACCGGAAGCAAACGACAATCCTCAGC<br>AAATTATCTACCACAACTCAC |
| Large inner H1_14 | ATTAGGTCAGGATTAGATTGATAATCGGA<br>AAATTATCTACCACAACTCAC |
| Large inner H1_15 | TAATAGAGCCAGCAGCAACATGAGGCCTTTAA<br>AAATTATCTACCACAACTCAC |
| Large inner H1_16 | AAAGCCTGCAACAGTGCACGGGTAAAAAG<br>AAATTATCTACCACAACTCAC |
| Large inner H1_17 | AAGAGAGGCGGTCACTATCCACTACTCACGTT<br>AAATTATCTACCACAACCTCAC |
| Large inner H1_18 | ATAGTTGAGCAGAAGATAAAATAAACATTGCG<br>AAATTATCTACCACAACCTCAC |
| Large inner H1_19 | CGTGAAAAATACCGAACGGAATACAAGAAAGG<br>AAATTATCTACCACAACTCAC |
| Large inner H1_20 | ACGTAGCCCTAAAACATCTTTGAGTTTCA<br>AAATTATCTACCACAACTCAC |
| Large inner H1_21 | GAAGAGTCTTTAATGCGCTACCAAGTTGCTAA<br>AAATTATCTACCACAACTCAC |
| Large inner H1_22 | TCGCTATGTAAATGGCTTAGAGCTTAATAAGGAGAAGTT<br>AAATTATCTACCACAACCTCAC |
| Large inner H1_23 | TTGGTATAATGCTGTAGCCTCCAAAAAATAC<br>AAATTATCTACCACAACCTCAC |
| Large inner H1_24 | CCTGTTTTAATATGCAATTTTTGAAGGC<br>AAATTATCTACCACAACCTCAC |
| Large inner H1_25 | AAAATACGGTGTCTGGAATAAAGGAGAAAGAG<br>AAATTATCTACCACAACTCAC |
| Large inner H1_26 | TAGTCCATATAACAGTTGAGAATCTAAAA<br>AAATTATCTACCACAACTCAC |
| Large inner H1_27 | ACGGAATTCTGCGAACGATTCAACACCCCCAG<br>AAATTATCTACCACAACTCAC |
| Large inner H1_28 | GTCTAGTTTGACCATTAGGGATTGCGGAA<br>AAATTATCTACCACAACCTCAC |
| Large inner H1_29 | AGGGAACCTTCTGACCTAATCCGCAACGATC<br>AAATTATCTACCACAACCTCAC |
| Large inner H1_30 | TTGCATTCTGGCCAACATTACTTCAGCCC<br>AAATTATCTACCACAACTCAC |
| Large inner H1_31 | CCGTCACGACCAGTAATACGCAGACCGCCTGT<br>AAATTATCTACCACAACTCAC |
| Large inner H1_32 | TAAAGCCCTGGCAGCATTGGCAGATTGAGGAACCTCACCA<br>AAATTATCTACCACAACCTCAC |
| Large inner H1_33 | CTCATCGTCTGAAATGGATTGAAAGTACCGTA<br>AAATTATCTACCACAACTCAC |
| Large inner H1_34 | TGTTACCTACATTTTGAGGTGTAAGCCCA<br>AAATTATCTACCACAACCTCAC |
| Large inner H1_35 | GGTCCAGGAAAAACGCTCTAGGCTGCCTCATT<br>AAATTATCTACCACAACTCAC |
| Large inner H1_36 | TTTAGTGCTTTATTTGGGGCGCGAGCTGTAGCGTGACCTG<br>AAATTATCTACCACAACTCAC |
| Large inner H1_37 | ATACGGCATCAATTCTACCCACAGAAGCCGGA<br>AAATTATCTACCACAACTCAC |
| Large inner H1_38 | CCAAGTAGCATTAAACATCTACAAGGTCAA<br>AAATTATCTACCACAACCTCAC |
| Large inner H1_39 | AGCAATCATACAGGCAAGAGTTTCGGAACCTGA<br>AAATTATCTACCACAACCTCAC |
| Large inner H1_40 | ACACTGGCAAAGAATTAGATAAGTTTTCCCATGAGGACA<br>AAATTATCTACCACAACTCAC |
| Large inner H1_41 | TGATTCACTGCCTTGAGTGATAGCACAGACCA<br>AAATTATCTACCACAACCTCAC |
| Large inner H1_42 | AAGCCCGTATAAACAGTCACCACGCTGAC<br>AAATTATCTACCACAACTCAC |
| Large inner H1_43 | TTTCGAACTCAAACCTATCTCAACGTGTTTAGT<br>AAATTATCTACCACAACCTCAC |
| Large inner H1_44 | GCTACATCACTTGCCTGATTCAGGTATCA<br>AAATTATCTACCACAACTCAC |
| Large inner H1_45 | CGAGATACTTCTTTGATTCCCTGACAAGTATA<br>AAATTATCTACCACAACTCAC |
| Large inner H1_46 | ATAGAAAGTAAAGCCGCAAATTAACCGTACGAGTCCGTCG<br>AAATTATCTACCACAACTCAC |
| Large inner H1_47 | AGCTAGTAAAAGAGTCTGTGAGATGGGTAATA<br>AAATTATCTACCACAACTCAC |
| Large inner H1_48 | GCGTTATAATCAGTGAGTTTAATAAGAAG<br>AAATTATCTACCACAACTCAC |
| Large inner H1_49 | TCGGCCAGAATCCTGAGACTTATGCATAGCGA<br>AAATTATCTACCACAACTCAC |
| Large inner H1_50 | GGTCATATATTGACTATTAAGAGGCTGACAGGAGAACAAA<br>AAATTATCTACCACAACTCAC |
| Large inner H1_51 | TTGACAAGAGAAGGATTAAATAGGTTGAATAA<br>AAATTATCTACCACAACTCAC |
| Large inner H1_52 | CAACGGGGTTTTGCTCATGATATGAGAAA<br>AAATTATCTACCACAACTCAC |
| Large inner H1_53 | GTAAAATGTTTTGAGTAAAT AAATTATCTACCACAACTCAC |
| Large inner H1_54 | TGTGTCCAATACTGCGGAGAGGGGTTTAA<br>AAATTATCTACCACAACTCAC |
| Large inner H1_55 | GACAATAAATATTCATTGTTTGCAACATTGTG<br>AAATTATCTACCACAACTCAC |
| Large inner H1_56 | CAACTCAAATGCTTTAAACCAAAGATTTT<br>AAATTATCTACCACAACTCAC |
| Large close_1 | GTGAATAACCTTGCTTCTGTAAATGAAGTATTAGACTTTACA<br>CACGACTC |
| Large close_2 | CTCATCGAGAACAAGCAAGCCGTTGATGCAAATCCAATCGCA<br>AACAGCCT |
| Large close_3 | CCGTAAAAAAAGCCGCACAGGCGGCAGCCAGCTTTCCGGCAC<br>CTCGTCGT |
| Large close_4 | TGTAGCCAGCTTTCATCAACATTAAACGCAAGGATAAAAATT<br>AGAAGGAA |
| Large close_5 | GGTACCGAGCTCGAATTCGTAATCGCAACCGCAAGAATGCCA<br>GCCTCAAT |
| Large close_6 | TAAAGCCAGAATGGAAAGCGCAGTAAATTATTCATTAAAGGT<br>AGCAGCTG |
| Large close_7 | GGGGAAAGCCGGCGAACGTGGCGACGTGCCAGCTGCATTAAT<br>GAACAGAG |
| Large close_8 | TACGCAGTATGTTAGCAAACGTAGCTACAATTTTATCCTGAA<br>ACTTCCGG |
| Large anti-close_1 | GAGTCGTG<br>TGTAAAGTCTAATACTTCATTTACAGAAGCAAGGTTATTCAC |
| Large anti-close_2 | AGGCTGTT<br>TGCGATTGGATTTGCATCAACGGCTTGCTTGTTCTCGATGAG |
| Large anti-close_3 | ACGACGAG<br>GTGCCGGAAAGCTGGCTGCCGCCTGTGCGGCTTTTTTTACGG |
| Large anti-close_4 | TTCCTTCT<br>AATTTTTATCCTTGCGTTTAATGTTGATGAAAGCTGGCTACA |
| Large anti-close_5 | ATTGAGGC<br>TGGCATTCTTGCGGTTGCGATTACGAATTCGAGCTCGGTACC |
| Large anti-close_6 | CAGCTGCT<br>ACCTTTAATGAATAATTTACTGCGCTTTCCATTCTGGCTTTA |
| Large anti-close_7 | CTCTGTTC<br>ATTAATGCAGCTGGCACGTCGCCACGTTCGCCGGCTTTCCCC |
| Large anti-close_8 | CCGGAAGT<br>TTCAGGATAAAATTGTAGCTACGTTTGCTAACATACTGCGTA |
| Large open_1 | GAGGCTTACCTTGCTGAACCTCAAATATTAGCAACATTTCTT<br>CATTTACG |
| Large open_2 | TTTAATTGCTCCTTGAGGTGAGGCTACA AGCAGCGG |
| Large open_3 | GCATATGC ATGAAAAATCTAAAGCATCTGAGGACATCAGCT |
| Large open_4 | TGCTTTCTTGATAAGAGGTCATTTTTGCGGATG GGCCTAG |
| Large open_5 | CTGCTCGC TTGTATCGGTTTTAAAGACTTTTT |
| Large open_6 | TATCATCGCACAGACAATATTTTTGAATACAACGGTGAATTT<br>TCAATTCC |
| Large open_7 | TTCGCAAATGGTCATAGTAAAAGATTTG TGACCACC |
| Large open_8 | GGCGCTAT GAAAGCGTAAGAATACGTGGCCTGATGTCTTTC |
| Large open_9 | CAGACGTATAACCTGTTTAGCTATATTTTCATT TCACGCAG |
| Large open_10 | AGTTGATG TAAAGTTTTGTCAAATTGTGTCGA |
| Large open_11 | CAAGAACAGAACAATATTACCGCCAGCCAAGAGTAGCCACCC<br>TCGCTGGC |
| Large open_12 | CCCTGCCTATTTCGCAGAACCATCTTGA AGGGCGGA |
| Large open_13 | ACGGGCTG GGCCTTGCTGGTAATATCCCGGATATAGAACCG |
| Large open_14 | CCACCCTGAACCTATTATTCTGAAACATGAAAG ATGCCCGG |
| Large open_15 | CGCGGCCC ACCGCCACCCTCTCATTACCCAAA |
| Large open_16 | GTCAGGATAAAGGGATTTTAGACAGGAAGGCTCATAGACGAC<br>CGCGCGAC |
| Large open_17 | AGAAAACGAGAATGGTTTACCTATACCA CCGACAGG |
| Large open_18 | ACACAACA AGCTAAACAGGAGGCCGATCGTTGGGCTATCAT |
| Large open_19 | AACCCTCACCATAAATCAAAAATCAGGTCTTTA GCGGTGCA |
| Large open_20 | CCTCAACC GTAAGAGCAACAAAGAAAAATCTA |
| Large anti-open_1 | CGTAAATG<br>AAGAAATGTTGCTAATATTTGAGGTTCAGCAAGGTAAGCCTC |
| Large anti-open_2 | CCGCTGCT TGTAGCCTCACCTCAAGGAGCAATTAAA |
| Large anti-open_3 | AGCTGATGTCCTCAGATGCTTTAGATTTTTCAT GCATATGC |
| Large anti-open_4 | CTAGTGCC CATCCGCAAAAATGACCTCTTATCAAGAAAGCA |
| Large anti-open_5 | AAAAAGTCTTTAAAACCGATACAA GCGAGCAG |
| Large anti-open_6 | GGAATTGA<br>AAATTCACCGTTGTATTCAAAAATATTGTCTGTGCGATGATA |
| Large anti-open_7 | GGTGGTCA CAAATCTTTTACTATGACCATTTGCGAA |
| Large anti-open_8 | GAAAGACATCAGGCCACGTATTCTTACGCTTTC ATAGCGCC |
| Large anti-open_9 | CTGCGTGA AATGAAAATATAGCTAAACAGGTTATACGTCTG |
| Large anti-open_10 | TCGACACAATTTGACAAAACTTTA CATCAACT |
| Large anti-open_11 | GCCAGCGA<br>GGGTGGCTACTCTTGGCTGGCGGTAATATTGTTCTGTTCTTG |
| Large anti-open_12 | TCCGCCCT TCAAGATGGTTCTGCGAAATAGGCAGGG |
| Large anti-open_13 | CGGTTCTATATCCGGGATATTACCAGCAAGGCC CAGCCCGT |
| Large anti-open_14 | CCGGGCAT CTTTCATGTTTCAGAATAATAGGTTCAGGGTGG |
| Large anti-open_15 | TTTGGGTAATGAGAGGGTGGCGGT GGGCCGCG |
| Large anti-open_16 | GTCGCGCG<br>GTCGTCTATGAGCCTTCCTGTCTAAAATCCCTTTATCCTGAC |
| Large anti-open_17 | CCTGTCGG TGGTATAGGTAAACCATTCTCGTTTTCT |
| Large anti-open_18 | ATGATAGCCCAACGATCGGCCTCCTGTTTAGCT TGTGTGT |
| Large anti-open_19 | TGCACCGC TAAAGACCTGATTTTTGATTTATGGTGAGGGTT |
| Large anti-open_20 | TAGATTTTTCTTTGTTGCTCTTAC GGTTGAGG |
| 5'-chol anti-H1 | /5CholTEG/GTGAGTTGTGGTAGATAATTT |
| 5'-AF647 anti-H2 | /5Alex647N/TAGATGGAGTGTGGTGTGAAG |
